# Computational deconvolution of gene expression in leukemic cell hierarchies

**DOI:** 10.1101/521864

**Authors:** Linnea Jäarvstråt, Ram Ajore, Anna-Karin Wihlborg, Urban Gullberg, Björn Nilsson

**Author notes:** Correspondence to: Björn Nilsson.

## Abstract

Leukemias and some solid tumors are organized as cell hierarchies, sustained by cancer stem cells. We developed a computational method to study gene expression cancer cell hierarchies. Unlike traditional approaches based on physical cell sorting, our method extracts cell type-specific gene expression signals from gene expression profiles of unsorted tumor cells by deconvolution. We apply our method in the context of acute myeloid leukemia, and recover markers for acute myeloid leukemia stem cells (AML-LSC).

## Introduction

Leukemias and some solid tumors are organized as cell hierarchies, sustained by a population of cancer stem cells (CSCs) at the apex.^1,2^ The hierarchical model has generated considerable interest as CSCs have properties that make them clinically relevant, including an ability to survive commonly employed therapies and to cause recurrences.^1^

An important step towards characterizing CSCs is to define their gene expression patterns in order to identify markers for directed therapy.^3,4,5,6^ Most attempts at this have relied on isolation and analysis of putative CSCs from tumor samples by cell sorting, which is logistically and technically demanding.

## Method

As an alternative approach, we explored the possibility of extracting CSC-associated gene expression patterns from gene expression profiles of unsorted tumor cells. Several thousands of such data have been published, and potentially allow detection of cell type-specific gene expression signals without physical cell sorting.

For this, we developed a computational method to extract cell type-specific gene expression signals from gene expression profiles of mixed blood cell populations. As described in detail in **Supplementary Methods**, our model that views a cancer cell hierarchy as a perturbed version of the normal cell hierarchy in the tissue-of-origin. To recover cell type-associated gene expression patterns using gene expression profiles of unsorted tumor samples where the proportions of cell types are unknown, we solve the optimization problem

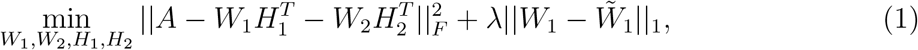

where *A* represents gene expression profiles of unsorted tumor cells, *W*_1_ gene expression profiles of each cell types in the tumor, *W*̃_1_ known reference gene expression profiles of the corresponding normal cell types (**Fig. 1a**), and *W*_2_ additional gene expression vectors to absorb additional variation (*e.g.*, batch effects). The columns of *H*_1_ and *H*_2_ represent non-negative mixing weights. Of note, this formulation differs from most previous deconvolution algorithms, which typically estimate either *W* or *H* only, not both at the same time (**Supplementary Methods**).

**Figure 1:**
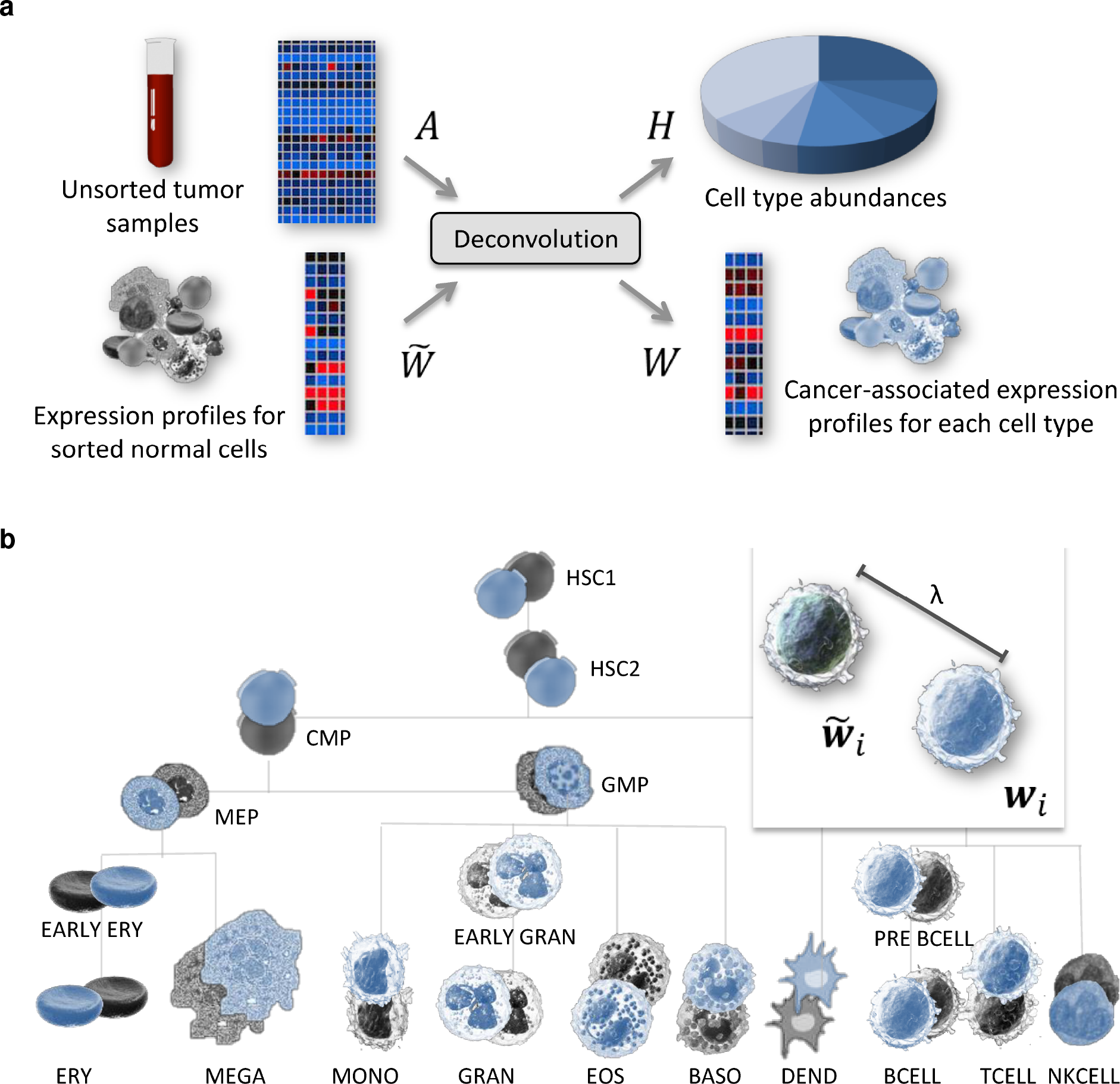
(a) Using gene expression profiles of unsorted tumor cells (*A*) and reference vectors for normal cells *W*̃, the developed method extracts cell type-specific gene expres-sion signatures (*W*) and their abundances (*H*). **(b, c)**We applied our method to 2,799 AML samples. To test whether any identified patterns represented AML-LSCs, we used gene expression profiles of functionally annotated AML cell fractions to annotate genes with AML-LSC activity (Supplementary Methods). Throughout, we observed enrichments of high AML-LSC activity scores among genes upregulated in leukemic HSC1 stem cell population, but not for other cell types (**Supplementary Figure 3**). Moreover, some of these genes were known AML-LSC markers.

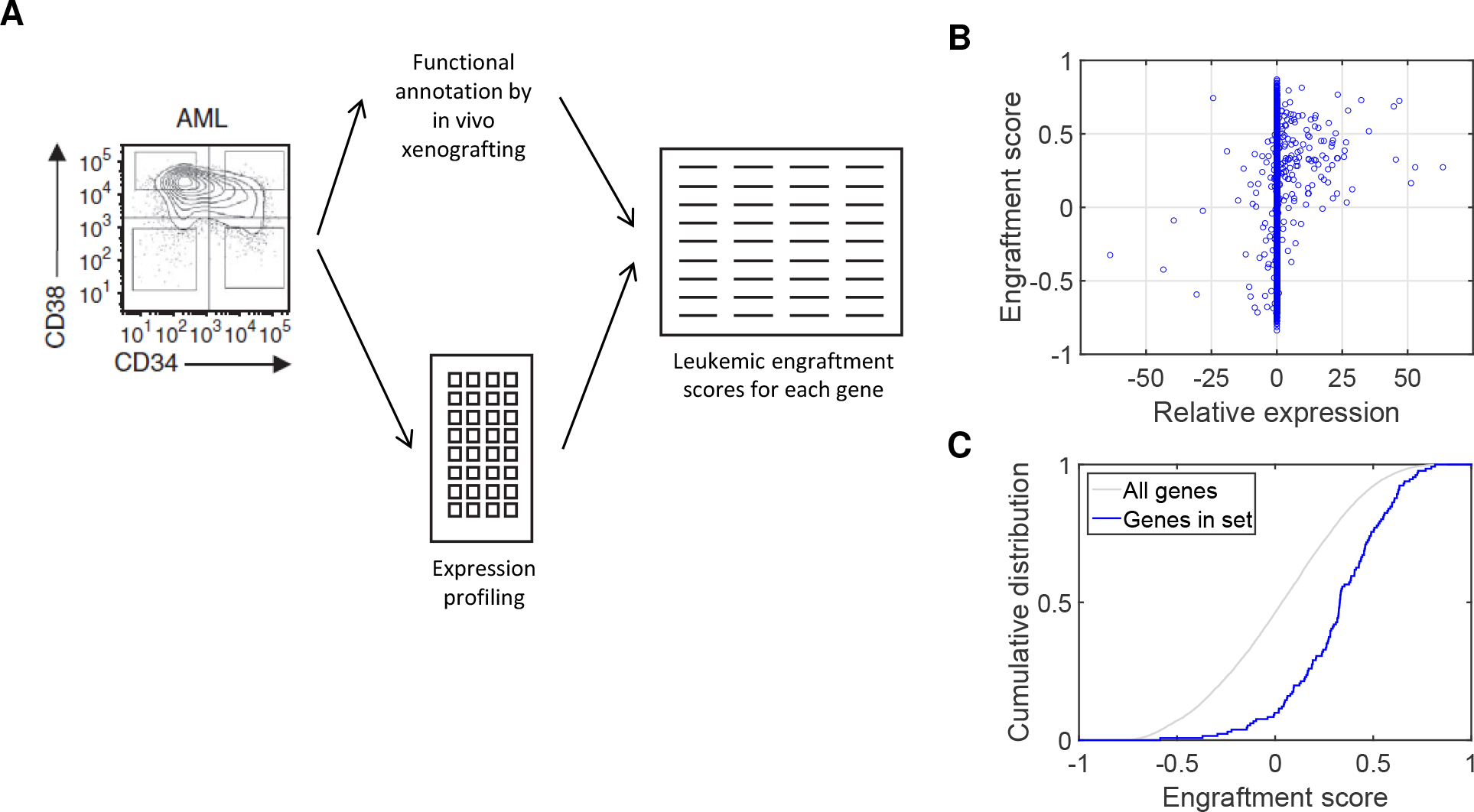

To solve the optimization problem, we adopted an efficient block-coordinate descent algorithm, ^7,8^ and added a bootstrapping step to increase robustness (**Supplementary Methods**). During the optimization, the second term in the objective function plays a key role. It guides the deconvolution process by keeping the extracted patterns in *W*_1_ close to the reference patterns in *W*̃_1_. This ensures that the extracted patterns can be interpreted as cell type-associated, rather than purely statistically driven entities. The term also imposes sparsity in that *W*_1_ and *W*̃_1_ will differ only at a few elements. The parameter λ > 0 controls the distance between the estimated and the reference patterns and the degree of sparsity.

## Results and Discussion

We tested our model on hematologic malignancies, which are marked by perturbed blood cell development, both in terms of cell type frequencies and gene expression. Several hematologic malignancies are hierarchically organized, including acute myeloid leukemia (AML), myelodysplastic syndrome, and chronic myeloid leukemia. ^9,10,11,12^

To illustrate the ability of our approach to identify CSC-relevant signals, we focused on leukemic stem cells in AML (AML-LSCs). For this, we retrieved gene expression data for 2,799 unsorted blood and bone marrow samples from patients with AML from ten previous studies (Supplementary Methods). As reference patterns, we used gene expression profiles of normal blood cells from D-Map,^13^ including data for two stem cell-enriched populations (”HSC1” and”HSC2” representing lineage-negative CD34^dim^38^−^133^+^ and CD34^+^38^−^ cells) (**Supplementary Fig. 1**).

We applied our method to the pooled set of 2,799 AML samples, to each AML data set individually, and to 1,554 examples of other hematologic malignancies.^14^ To test whether any of the extracted patterns represented AML-LSCs, we used gene expression profiles of sorted AML cell fractions (CD34^+^38^−^, CD34^+^38^+^, CD34^−^38^+^ and CD34^−^38^−^ cells), which were gene expression-profiled and assayed for AML-LSC activity using a limiting dilution assay based on *in vivo* xenografting).^9^ Using these data, we calculated engraftment scores that quantitatively reflect the relevance of each gene with leukemic engraftment.^9^ We then tested for enrichments of high scores among genes found to be differently expressed in the cell type-specific gene expression profiles of unsorted AML samples.

Throughout, we observed enrichments of genes with high AML-LSC relevance scores among genes with increased expression in leukemic HSC1 cells (*i.e.*, the most primitive cell type in the model) (**Fig. 1b,c** and **Supplementary Fig. 2**). By contrast, we saw no such enrichments for any other cell type-specific patterns deconvolved using AML data (**Supplementary Fig. 3**), nor in any patterns deconvolved using data from other hematologic malignancies (**Supplementary Fig. 4**). Moreover, the set of genes up-regulated in the AML-HSC1 pattern included multiple known AML-LSC markers, including *IL3RA*, *CD96*, and *IL1RAP* (**Supplementary Table 1**).^4,5^

In summary, we developed a computational approach to infer cell type-specific gene expression patterns through deconvolution of gene expression profiles of unsorted cells. Compared to physical cell sorting and single-cell approaches, deconvolution represents a complementary method that is easy to use, but has a limitation in that small cell populations may be hard to detect. Nevertheless, our results illustrate that signals representing CSCs can be extracted, probably because of power gained by analyzing thousands of samples, and because reference vectors to guide the analysis. The described method has been implemented as a software tool that is available for Windows and Linux, and accepts input data in standard tab-delimited format.

## Acknowledgements

The study was supported by research grants from Knut and Alice Wallenberg’s Foundation (no. 2012.0193), Vetenskapsrådet (no. 2012-1753), the Swedish Children’s Cancer Fund (no. PR2015-0028), the Swedish Cancer Fund (no. 2014/482), the Swedish Foundation for Strategic Research (no. ICA08-0057 and KF10-0009), and ALF grants from Region Skåne.

## Author contributions

L.J. and B.N. developed the method and carried out the experiments. R.A. carried out additional analyses. U.G. contributed the to experimental design and the conception of the project, with additional input from A.K.W. All authors contributed to the final manuscript.

## Competing interests

The authors declare no competing financial interests.

## Notes

Conicts-of-interest: none.

## References

1. Kreso A, Dick J. E. Evolution of the cancer stem cell model․. Cell Stem Cell 2014; 14: 275–291.

2. Meacham C. E, Morrison S. J. Tumour heterogeneity and cancer cell plasticity․. Nature 2013; 501: 328–337.

3. Jin L, Hope K. J, Zhai Q, Smadja-Joffe F, Dick J. E. Targeting of CD44 eradicates human acute myeloid leukemic stem cells․. Nat Med 2006; 12: 1167–1174.

4. Jin L, Lee E. M, Ramshaw H. S, Busfield S. J, Peoppl A. G, Wilkinson L, Guthridge M. A, Thomas D, Barry E. F, Boyd A, Gearing D. P, Vairo G, Lopez A. F, Dick J. E, Lock R. B. Monoclonal antibody-mediated targeting of CD123, IL-3 receptor alpha chain, eliminates human acute myeloid leukemic stem cells․. Cell Stem Cell 2009; 5: 31–42.

5. Majeti R. Monoclonal antibody therapy directed against human acute myeloid leukemia stem cells․. Oncogene 2011; 30: 1009–1019.

6. Ågerstam H, Karlsson C, Hansen N, Sandén C, Askmyr M, von Palffy S, Högberg C, Rissler M, Wunderlich M, Juliusson G, Richter J, Sjöstrom K, Bhatia R, Mulloy J. C, Järås M, Fioretos T. Antibodies targeting human IL1RAP (IL1R3) show therapeutic effects in xenograft models of acute myeloid leukemia․. Proc Natl Acad Sci U S A 2015; 112: 10786–10791.

7. Taslaman L, Nilsson B. A framework for regularized non-negative matrix factorization, with application to the analysis of gene expression data․. PLoS One 2012; 7: e46331.

8. Benthem, M.H. Keenan M. Fast algorithm for the solution of large-scale non-negativity-constrained least squares problems. Journal of Chemometrics 2004; 18: 441–450.

9. Eppert K, Takenaka K, Lechman E. R, Waldron L, Nilsson B, van Galen P, Metzeler K. H, Poeppl A, Ling V, Beyene J, Canty A. J, Danska J. S, Bohlander S. K, Buske C, Minden M. D, Golub T. R, Jurisica I, Ebert B. L, Dick J. E. Stem cell gene expression programs influence clinical outcome in human leukemia․. Nat Med 2011; 17: 1086–1093.

10. Larochelle A, Vormoor J, Hanenberg H, Wang J. C, Bhatia M, Lapidot T, Moritz T, Murdoch B, Xiao X. L, Kato I, Williams D. A, Dick J. E. Identification of primitive human hematopoietic cells capable of repopulating NOD/SCID mouse bone marrow: implications for gene therapy․. Nat Med 1996; 2: 1329–1337.

11. Tehranchi R et al. Persistent malignant stem cells in del(5q) myelodysplasia in remission․. N Engl J Med 2010; 363: 1025–1037.

12. Woll P. S et al. Myelodysplastic syndromes are propagated by rare and distinct human cancer stem cells in vivo․. Cancer Cell 2014; 25: 794–808.

13. Novershtern N, Subramanian A, Lawton L. N, Mak R. H, Haining W. N, McConkey M. E, Habib N, Yosef N, Chang C. Y, Shay T, Frampton G. M, Drake A. C. B, Leskov I, Nilsson B, Preffer F, Dombkowski D, Evans J. W, Liefeld T, Smutko J. S, Chen J, Friedman N, Young R. A, Golub T. R, Regev A, Ebert B. L. Densely interconnected transcriptional circuits control cell states in human hematopoiesis․. Cell 2011; 144: 296–309.

14. Haferlach T et al. Clinical utility of microarray-based gene expression profiling in the diagnosis and subclassification of leukemia: report from the International Microarray Innovations in Leukemia Study Group․. J Clin Oncol 2010; 28: 2529–2537.

