## Supplementary Figure 1 for "Computational deconvolution of gene expression in leukemic cell hierarchies"

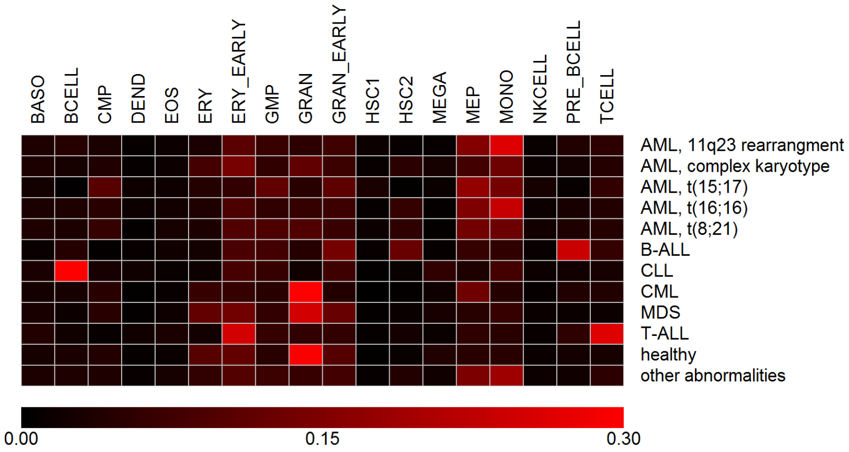


Our deconvolution approach estimates both cell type-associated gene expression patterns and proportions of the different cell types in the mix. To exemplify the ability of our approach to recover cell type proportions in addition to cell type-specific signals, we applied our deconvolution method to a set of 2,096 samples from different conditions[1], including Acute Lymphoblastic Leukemia (ALL), Acute Myeloid Leukemia (AML), Chronic Lymphocytic Leukemia (CLL), Chronic Myeloid Leukemia (CML), and Myelodysplastic Syndrome (MDS).

We plotted a heatmap of the average mixing weights ($H_1$ values) for cell type in each condition. As shown, we recovered changes in cell type proportions that are characteristic for the different conditions. For example, we observed an abundance of T cells in T-ALL, an abundance of pre-B cells in B-ALL (where the tumor cells typically show a pre-B phenotype), and an abundance of B cells in CLL (where the tumor cells typically show a mature B cell phenotype). In the AML samples, we observed enrichments of early myeloid progenitors (MEPs and CMPs), as well as an enrichment of monocytes, particularly in AML with 11q23 rearrangement. These changes are consistent with typical AML phenotypes. In the CML samples, we observed showed an enrichment of MEPs and granulocytes, which is consistent with the myeloid expansion seen in CML. Finally, the MDS samples showed cell type proportions similar to that of healthy individuals. In summary, this example illustrates the ability of our approach to recover relevant cell type proportions. In this example, we used $\lambda = 0.1$. Similar results were obtained with other reasonable $\lambda$ values.

### References

1. Haferlach, T. *et al.* Clinical utility of microarray-based gene expression profiling in the diagnosis and subclassification of leukemia: report from the international microarray innovations in leukemia study group. *J Clin Oncol* **28**, 2529–2537 (2010). URL `http://dx.doi.org/10.1200/JCO.2009.23.4732`.
