## Supplementary Figure 2 for "Computational deconvolution of gene expression in leukemic cell hierarchies"

Enrichment plots of leukemic engraftment scores in sets of genes found to be up-regulated compared to the normal for the HSC1 cell type in each of the ten AML data sets tested. In most of them, we detected significant enrichments of positive leukemic engraftment scores for HSC1 cells. The numbers of up-regulated genes and p values for enrichment are indicated above each panel. Similar results were seen with other reasonable $\lambda$ values.

**$\lambda$=0.10**

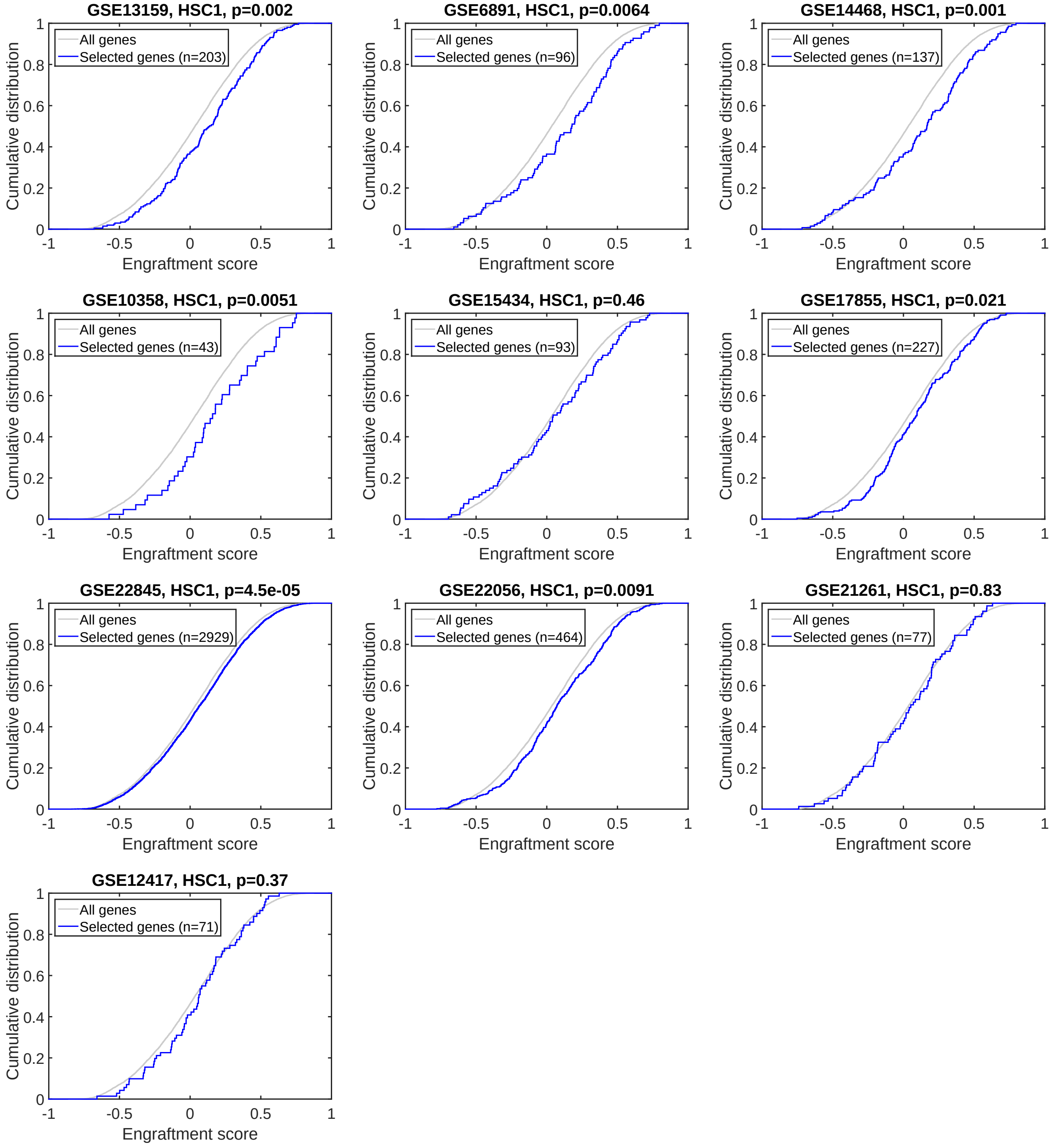
