## Supplementary Figure 3 for "Computational deconvolution of gene expression in leukemic cell hierarchies"

Enrichment plots of leukemic engraftment scores in sets of genes found to be up-regulated compared to the normal for the different cell types in the deconvolution model with different $\lambda$ values. As shown, the only consistent enrichment of positive leukemic engraftment scores was detected for HSC1 cells (*i.e.*, the most primitive stem cell population in the model).

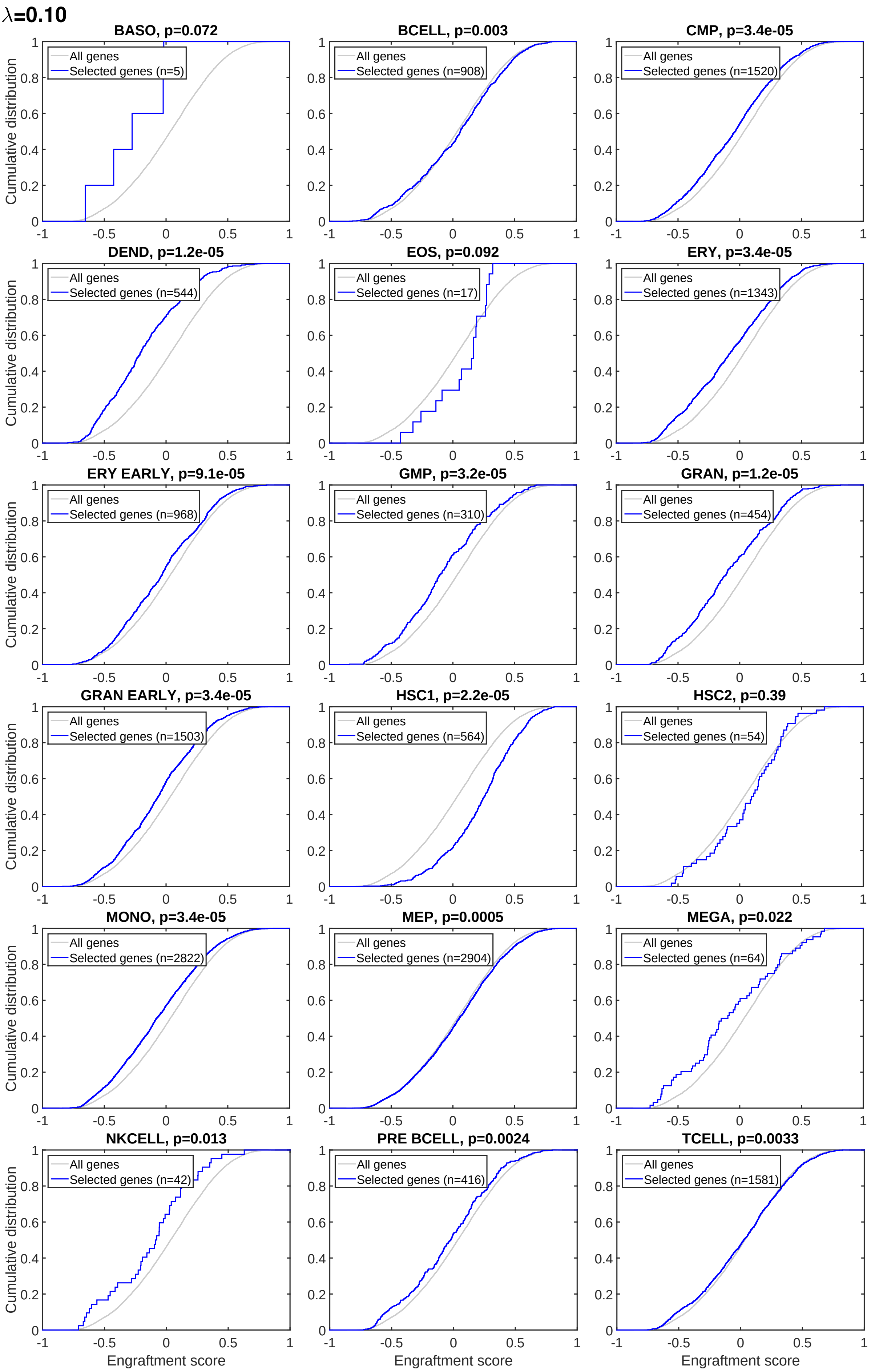
λ=0.10
BASO, p=0.072
All genes
Selected genes (n=5)
BCELL, p=0.003
All genes
Selected genes (n=908)
CMP, p=3.4e-05
All genes
Selected genes (n=1520)
DEND, p=1.2e-05
All genes
Selected genes (n=544)
EOS, p=0.092
All genes
Selected genes (n=17)
ERY, p=3.4e-05
All genes
Selected genes (n=1343)
ERY EARLY, p=9.1e-05
All genes
Selected genes (n=968)
GMP, p=3.2e-05
All genes
Selected genes (n=310)
GRAN, p=1.2e-05
All genes
Selected genes (n=454)
GRAN EARLY, p=3.4e-05
All genes
Selected genes (n=1503)
HSC1, p=2.2e-05
All genes
Selected genes (n=564)
HSC2, p=0.39
All genes
Selected genes (n=54)
MONO, p=3.4e-05
All genes
Selected genes (n=2822)
MEP, p=0.0005
All genes
Selected genes (n=2904)
MEGA, p=0.022
All genes
Selected genes (n=64)
NKCELL, p=0.013
All genes
Selected genes (n=42)
PRE BCELL, p=0.0024
All genes
Selected genes (n=416)
TCELL, p=0.0033
All genes
Selected genes (n=1581)
Cumulative distribution
Engraftment score

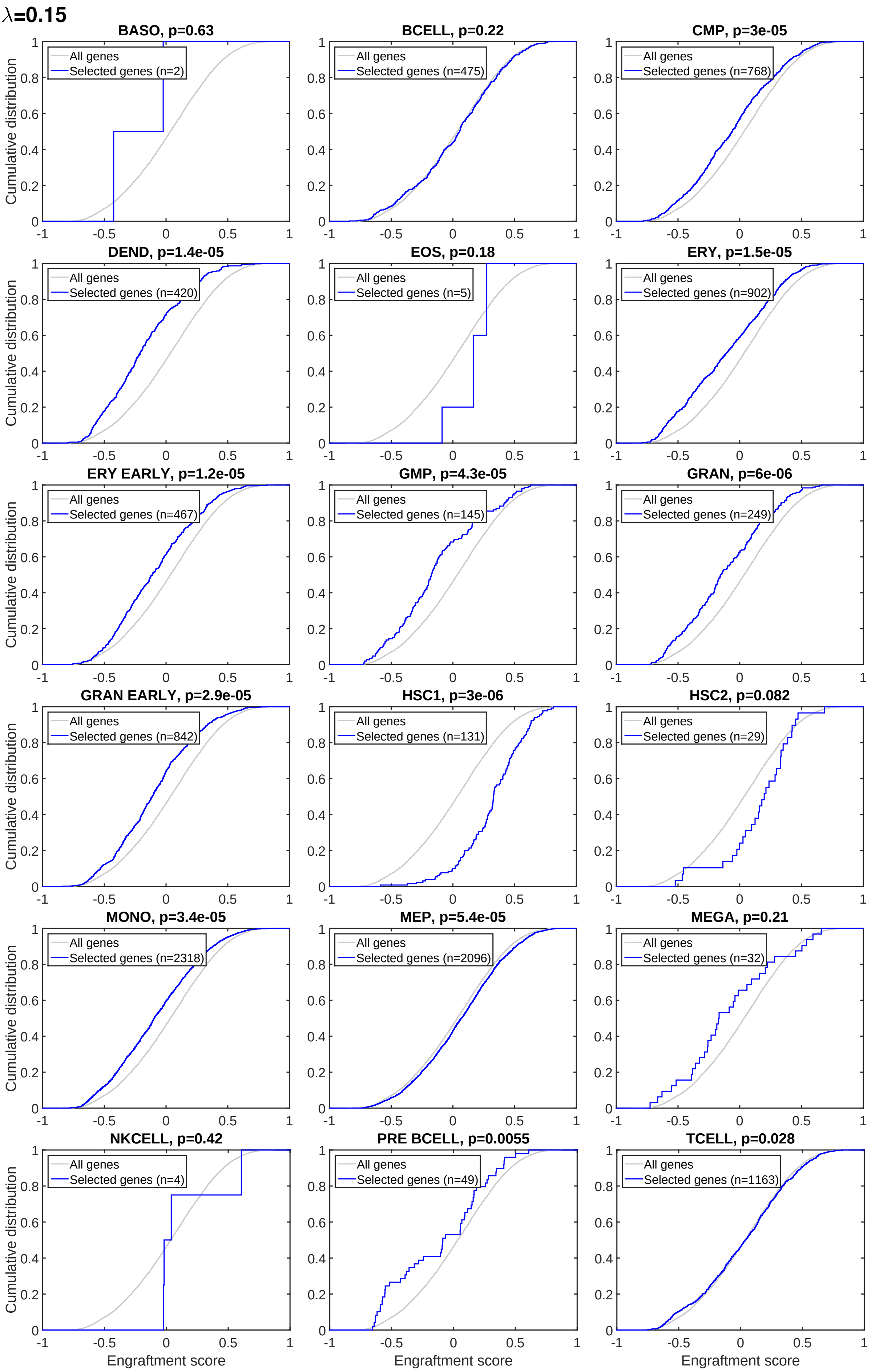

λ=0.15
BASO, p=0.63
BCELL, p=0.22
CMP, p=3e-05
DEND, p=1.4e-05
EOS, p=0.18
ERY, p=1.5e-05
ERY EARLY, p=1.2e-05
GMP, p=4.3e-05
GRAN, p=6e-06
GRAN EARLY, p=2.9e-05
HSC1, p=3e-06
HSC2, p=0.082
MONO, p=3.4e-05
MEP, p=5.4e-05
MEGA, p=0.21
NKCELL, p=0.42
PRE BCELL, p=0.0055
TCELL, p=0.028
All genes
Selected genes (n=2)
Selected genes (n=475)
Selected genes (n=768)
Selected genes (n=420)
Selected genes (n=5)
Selected genes (n=902)
Selected genes (n=467)
Selected genes (n=145)
Selected genes (n=249)
Selected genes (n=842)
Selected genes (n=131)
Selected genes (n=29)
Selected genes (n=2318)
Selected genes (n=2096)
Selected genes (n=32)
Selected genes (n=4)
Selected genes (n=49)
Selected genes (n=1163)
Cumulative distribution
Engraftment score

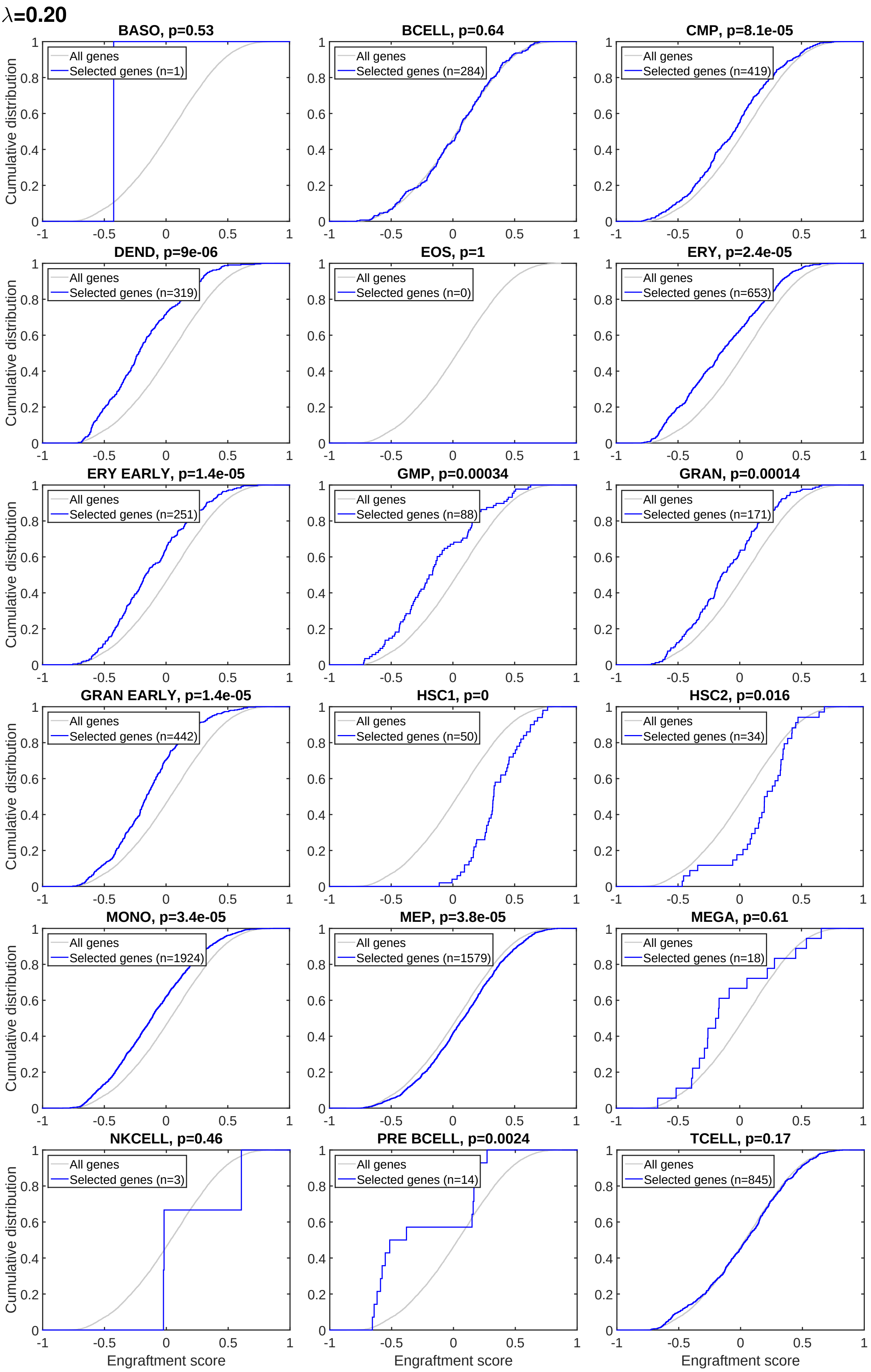
λ=0.20
BASO, p=0.53
BCELL, p=0.64
CMP, p=8.1e-05
DEND, p=9e-06
EOS, p=1
ERY, p=2.4e-05
ERY EARLY, p=1.4e-05
GMP, p=0.00034
GRAN, p=0.00014
GRAN EARLY, p=1.4e-05
HSC1, p=0
HSC2, p=0.016
MONO, p=3.4e-05
MEP, p=3.8e-05
MEGA, p=0.61
NKCELL, p=0.46
PRE BCELL, p=0.0024
TCELL, p=0.17
All genes
Selected genes (n=1)
Selected genes (n=284)
Selected genes (n=419)
Selected genes (n=319)
Selected genes (n=0)
Selected genes (n=653)
Selected genes (n=251)
Selected genes (n=88)
Selected genes (n=171)
Selected genes (n=442)
Selected genes (n=50)
Selected genes (n=34)
Selected genes (n=1924)
Selected genes (n=1579)
Selected genes (n=18)
Selected genes (n=3)
Selected genes (n=14)
Selected genes (n=845)
Cumulative distribution
Engraftment score

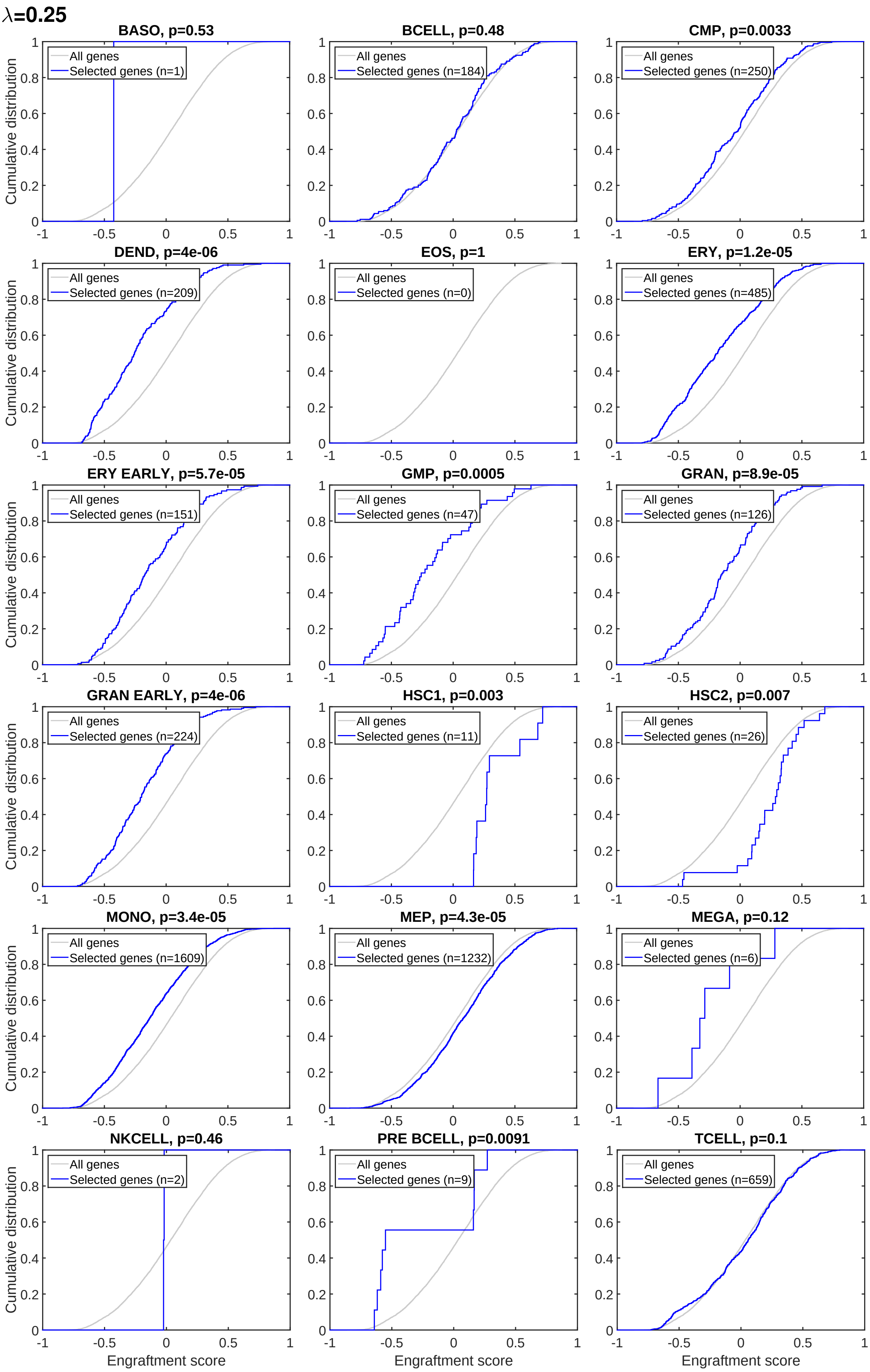

λ=0.25
BASO, p=0.53
All genes
Selected genes (n=1)
BCELL, p=0.48
All genes
Selected genes (n=184)
CMP, p=0.0033
All genes
Selected genes (n=250)
DEND, p=4e-06
All genes
Selected genes (n=209)
EOS, p=1
All genes
Selected genes (n=0)
ERY, p=1.2e-05
All genes
Selected genes (n=485)
ERY EARLY, p=5.7e-05
All genes
Selected genes (n=151)
GMP, p=0.0005
All genes
Selected genes (n=47)
GRAN, p=8.9e-05
All genes
Selected genes (n=126)
GRAN EARLY, p=4e-06
All genes
Selected genes (n=224)
HSC1, p=0.003
All genes
Selected genes (n=11)
HSC2, p=0.007
All genes
Selected genes (n=26)
MONO, p=3.4e-05
All genes
Selected genes (n=1609)
MEP, p=4.3e-05
All genes
Selected genes (n=1232)
MEGA, p=0.12
All genes
Selected genes (n=6)
NKCELL, p=0.46
All genes
Selected genes (n=2)
PRE BCELL, p=0.0091
All genes
Selected genes (n=9)
TCELL, p=0.1
All genes
Selected genes (n=659)
Cumulative distribution
Engraftment score
