## Supplementary Figure 4 for "Computational deconvolution of gene expression in leukemic cell hierarchies"

Enrichment plots of leukemic engraftment scores in sets of genes found to be up-regulated compared to the normal for the HSC1 cell type in 1,554 non-AML samples[1]. In contrast to AML samples (Supplementary Figure 1 and 2), we detected no consistent enrichments of positive leukemic engraftment scores for HSC1 cells with non-AML samples. The numbers of up-regulated genes and p values for enrichment are indicated above each panel.

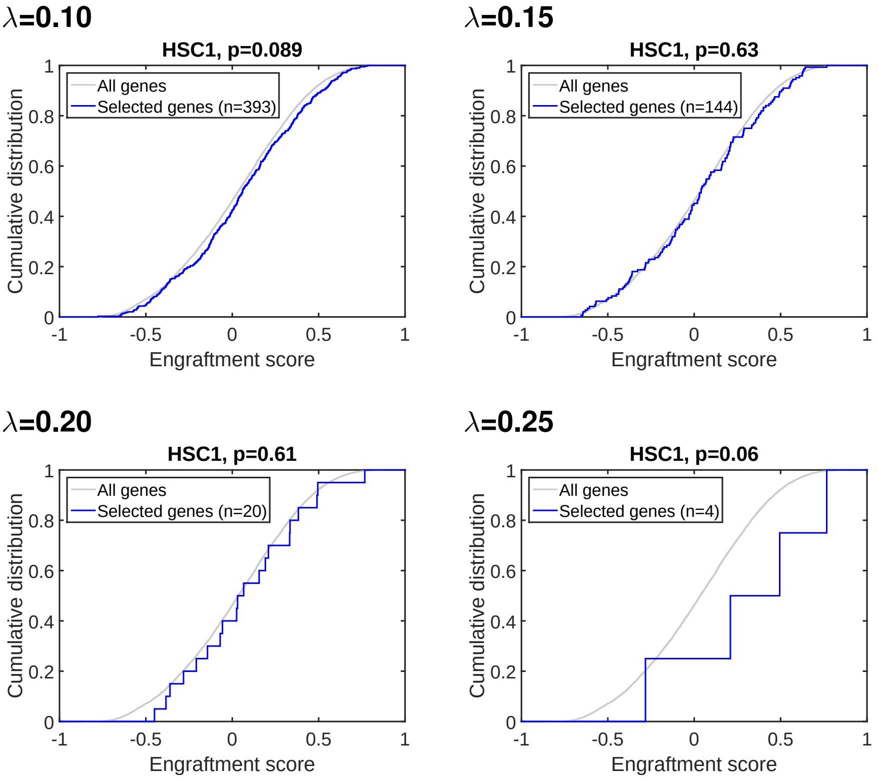


### References

1. Haferlach, T. *et al.* Clinical utility of microarray-based gene expression profiling in the diagnosis and subclassification of leukemia: report from the international microarray innovations in leukemia study group. *J Clin Oncol* **28**, 2529–2537 (2010). URL `http://dx.doi.org/10.1200/JCO.2009.23.4732`.
