## Supplementary Figure 5 for "Computational deconvolution of gene expression in leukemic cell hierarchies"

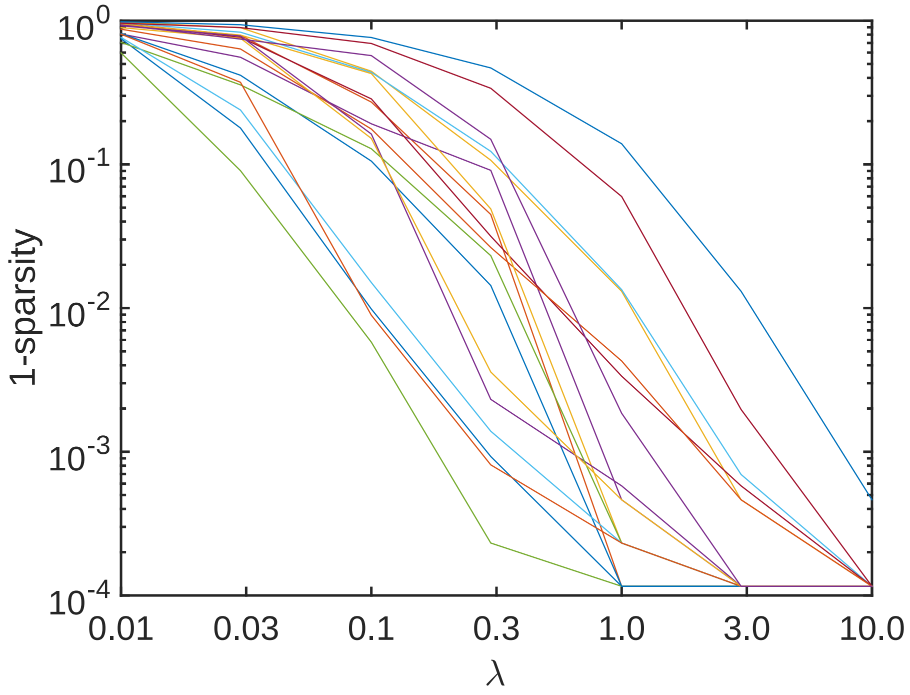


Sparsity of $W_1 - \tilde{W}_1$ per cell type, as a function of $\lambda$. In our model, the parameter $\lambda$ controls the distance between the estimated patterns in $W_1$ and the reference patterns in $\tilde{W}_1$, as well as the sparsity of the difference matrix $W_1 - \tilde{W}_1$. A low $\lambda$ allows greater freedom in the estimation, while a high $\lambda$ will keep the estimated patterns closer to the reference patterns. This figure illustrates the sparsity of $W_1 - \tilde{W}_1$ per column (cell type) for the pooled set of 2,799 AML samples.
