## Supplementary Table 1 for "Computational deconvolution of gene expression in leukemic cell hierarchies"

Difference between extracted and reference HSC1 vectors for the 2,799 AML samples for different values of λ. Only genes with non-zero difference shown.

| | | Relative expression for HSC1 | | | |
|---|---|---|---|---|---|
| **GeneID** | **GeneSymbol** | **λ=0.10** | **λ=0.15** | **λ=0.20** | **λ=0.25** |
| 29064 | *FAM30A* | 41.982469 | 38.692869 | 31.516392 | 23.045118 |
| 9834 | *KIAA0125* | 37.381947 | 32.987672 | 26.938371 | 18.356502 |
| 1791 | *DNTT* | 30.662212 | 22.884979 | 10.804217 | 0.000000 |
| 56654 | *NPDC1* | 27.447258 | 20.101661 | 9.476397 | 0.551442 |
| 27287 | *VENTX* | 26.347776 | 23.994494 | 12.806278 | 1.682255 |
| 2620 | *GAS2* | 25.878834 | 22.296992 | 15.690719 | 4.204950 |
| 9805 | *SCRN1* | 24.209533 | 20.096722 | 12.498550 | 4.551375 |
| 7098 | *TLR3* | 23.860083 | 21.357251 | 15.767367 | 5.825480 |
| 54762 | *GRAMD1C* | 22.590506 | 18.607444 | 12.477669 | 4.249300 |
| 79870 | *BAALC* | 22.482438 | 17.390707 | 10.428391 | 2.732282 |
| 2982 | *GUCY1A3* | 21.833249 | 17.100435 | 10.622427 | 2.263355 |
| 3728 | *JUP* | 20.858127 | 16.354875 | 8.566455 | 1.615124 |
| 3563 | *IL3RA* | 20.653463 | 19.393127 | 16.078647 | 8.126556 |
| 81615 | *TMEM163* | 20.130805 | 17.005314 | 10.622882 | 1.536283 |
| 6192 | *RPS4Y1* | 19.943206 | 35.821280 | 104.702466 | 135.768501 |
| 3617 | *IMPG1* | 19.285433 | 15.636070 | 10.942500 | 5.929242 |
| 3212 | *HOXB2* | 18.758877 | 15.334269 | 9.100464 | 0.953251 |
| 6335 | *SCN9A* | 18.561249 | 15.066636 | 10.261285 | 1.681084 |
| 55351 | *STK32B* | 18.492339 | 15.243634 | 8.300268 | 0.000000 |
| 55713 | *ZNF334* | 18.217932 | 12.916452 | 8.995219 | 2.406024 |
| 3213 | *HOXB3* | 17.860660 | 13.766579 | 5.202923 | 0.000000 |
| 8287 | *USP9Y* | 17.545574 | 18.585242 | 41.771147 | 52.041407 |
| 11156 | *PTP4A3* | 17.378177 | 15.180075 | 10.027333 | 2.654390 |
| 79839 | *CCDC102B* | 16.476058 | 13.565662 | 9.191089 | 2.433015 |
| 4883 | *NPR3* | 16.446908 | 10.239864 | 0.000000 | 0.000000 |
| 7552 | *ZNF711* | 16.391477 | 10.319276 | 6.002023 | 0.000000 |
| 58189 | *WFDC1* | 16.268422 | 14.735299 | 8.313601 | 2.016175 |
| 2273 | *FHL1* | 16.238708 | 12.609076 | 5.570401 | 0.000000 |
| 1795 | *DOCK3* | 16.175998 | 10.387777 | 2.554465 | 0.000000 |
| 3445 | *IFNA8* | 16.084258 | 12.921448 | 8.421084 | 1.826542 |
| 23015 | *GOLGA8A* | 15.903786 | 10.875053 | 3.210699 | 0.000000 |
| 4192 | *MDK* | 15.712135 | 12.597313 | 4.247330 | 0.000000 |
| 4885 | *NPTX2* | 15.576666 | 12.411242 | 7.921605 | 1.361587 |
| 3642 | *INSM1* | 15.411624 | 13.255774 | 13.601881 | 9.282459 |
| 586 | *BCAT1* | 15.362933 | 12.803651 | 8.107247 | 1.576660 |
| 55415 | *PRO2964* | 15.306280 | 10.516626 | 4.651588 | 0.000000 |
| 1043 | *CD52* | 15.254058 | 13.669877 | 8.852977 | 3.794841 |
| 3075 | *CFH* | 15.118782 | 10.583680 | 1.987221 | 0.000000 |
| 4068 | *SH2D1A* | 15.057037 | 11.120678 | 4.497509 | 0.000000 |
| 51678 | *MPP6* | 15.041422 | 11.830817 | 6.092625 | 0.000000 |

| | | | | | |
|---|---|---|---|---|---|
| 10225 | *CD96* | 15.005079 | 12.155250 | 6.701106 | 0.657204 |
| 5891 | *RAGE* | 14.754651 | 12.596922 | 8.815629 | 3.087843 |
| 5874 | *RAB27B* | 14.415216 | 11.155767 | 7.741661 | 2.630572 |
| 4038 | *LRP4* | 14.315493 | 11.674968 | 8.410671 | 5.220715 |
| 26281 | *FGF20* | 14.247240 | 9.945811 | 3.413513 | 0.000000 |
| 80723 | *TMEM22* | 14.163095 | 9.389456 | 5.978592 | 0.613178 |
| 55655 | *NLRP2* | 13.900645 | 7.933475 | 0.000000 | 0.000000 |
| 284 | *ANGPT1* | 13.868512 | 9.847847 | 5.351740 | 0.000000 |
| 9289 | *GPR56* | 13.619996 | 10.828027 | 7.457569 | 1.812503 |
| 653483 | *LOC653483* | 13.551050 | 10.419294 | 2.892094 | 0.000000 |
| 8876 | *VNN1* | 13.428097 | 9.295478 | 6.813717 | 2.838799 |
| 79957 | *PAQR6* | 13.346826 | 11.150060 | 7.991633 | 4.085587 |
| 1396 | *CRIP1* | 13.334948 | 11.575575 | 7.107814 | 3.217286 |
| 1191 | *CLU* | 13.144812 | 10.364637 | 2.102059 | 0.000000 |
| 56938 | *ARNTL2* | 13.097699 | 10.282863 | 5.597111 | 0.000000 |
| 9053 | *MAP7* | 13.019501 | 9.085744 | 4.720742 | 1.073364 |
| 9379 | *NRXN2* | 12.962914 | 9.037114 | 0.664007 | 0.000000 |
| 64221 | *ROBO3* | 12.892303 | 11.088181 | 7.763651 | 1.437766 |
| 54435 | *HCG4* | 12.746324 | 9.737109 | 4.485006 | 0.000000 |
| 23682 | *RAB38* | 12.590604 | 10.626138 | 3.091657 | 0.000000 |
| 1287 | *COL4A5* | 12.141442 | 9.216493 | 2.791421 | 0.000000 |
| 4330 | *MN1* | 12.124319 | 8.869518 | 4.219415 | 0.000000 |
| 2940 | *GSTA3* | 12.097321 | 8.910521 | 0.806795 | 0.000000 |
| 57730 | *KIAA1641* | 12.042144 | 6.689845 | 2.062325 | 0.000000 |
| 51237 | *MGC29506* | 11.992304 | 9.791677 | 5.740940 | 0.000000 |
| 26049 | *KIAA0888* | 11.982317 | 5.693404 | 0.000000 | 0.000000 |
| 28965 | *SLC27A6* | 11.808326 | 7.962302 | 3.966421 | 0.358870 |
| 7503 | *XIST* | 11.783191 | 0.000000 | -30.984569 | -49.530604 |
| 440270 | *GOLGA8B* | 11.770227 | 6.201873 | 0.000000 | 0.000000 |
| 64111 | *NPVF* | 11.654639 | 7.810163 | 4.552779 | 0.000000 |
| 2769 | *GNA15* | 11.514055 | 10.659197 | 8.054065 | 4.924851 |
| 9808 | *KIAA0087* | 11.388464 | 5.971199 | 0.000000 | 0.000000 |
| 3339 | *HSPG2* | 11.346348 | 7.220116 | 3.730137 | 0.000000 |
| 8835 | *SOCS2* | 11.257680 | 8.248814 | 2.917136 | 0.000000 |
| 1007 | *CDH9* | 11.198415 | 8.067194 | 2.945669 | 0.000000 |
| 8284 | *JARID1D* | 11.146166 | 14.359293 | 49.211460 | 64.021631 |
| 55304 | *SPTLC3* | 11.067287 | 5.804966 | 0.795253 | 0.000000 |
| 1612 | *DAPK1* | 11.063465 | 5.709455 | 0.000000 | 0.000000 |
| 7432 | *VIP* | 10.834995 | 6.318493 | 0.611823 | 0.000000 |
| 79841 | *AGBL2* | 10.761966 | 7.917653 | 2.239165 | 0.000000 |
| 2262 | *GPC5* | 10.690484 | 7.324820 | 1.475206 | 0.000000 |
| 7163 | *TPD52* | 10.678847 | 5.342783 | 0.000000 | 0.000000 |
| 26289 | *AK5* | 10.678245 | 6.893812 | 5.137628 | 1.582035 |
| 225 | *ABCD2* | 10.667587 | 7.095266 | 6.826152 | 2.566138 |
| 4744 | *NEFH* | 10.647312 | 8.187677 | 5.388843 | 0.334696 |
| 5881 | *RAC3* | 10.511229 | 4.943090 | 0.000000 | 0.000000 |
| 3357 | *HTR2B* | 10.340547 | 7.499084 | 2.990083 | 0.000000 |
| 9122 | *SLC16A4* | 10.329328 | 5.171779 | 0.000000 | 0.000000 |

| | | | | | |
|---|---|---|---|---|---|
| 10740 | *RFPL1S* | 10.271519 | 7.704768 | 4.171454 | 0.000000 |
| 55410 | *CYorf14* | 10.270865 | 9.736191 | 19.440442 | 21.426605 |
| 6524 | *SLC5A2* | 10.227602 | 6.676428 | 1.696704 | 0.000000 |
| 9124 | *PDLIM1* | 10.134693 | 7.418663 | 1.925338 | 0.000000 |
| 54541 | *DDIT4* | 10.119938 | 7.755161 | 5.506767 | 0.942679 |
| 10040 | *TOM1L1* | 10.082056 | 8.580372 | 5.548014 | 0.000000 |
| 4345 | *CD200* | 10.074270 | 4.593930 | 0.000000 | 0.000000 |
| 4267 | *CD99* | 10.039710 | 6.995551 | 0.708655 | 0.000000 |
| 23101 | *MCF2L2* | 10.036177 | 4.820883 | 0.000000 | 0.000000 |
| 374977 | *TTC4* | 9.973784 | 6.100700 | 0.000000 | 0.000000 |
| 84679 | *SLC9A7* | 9.942676 | 5.840967 | 1.261242 | 0.000000 |
| 1466 | *CSRP2* | 9.888563 | 5.570451 | 0.194200 | 0.000000 |
| 150759 | *LOC150759* | 9.721534 | 5.777426 | 0.816281 | 0.000000 |
| 91120 | *ZNF682* | 9.709292 | 5.171664 | 0.000000 | 0.000000 |
| 285 | *ANGPT2* | 9.555957 | 5.585873 | 0.408308 | 0.000000 |
| 80070 | *ADAMTS20* | 9.545399 | 4.901182 | 0.258692 | 0.000000 |
| 4922 | *NTS* | 9.444836 | 4.150146 | 0.138990 | 0.000000 |
| 5090 | *PBX3* | 9.366077 | 5.946428 | 0.000000 | 0.000000 |
| 55506 | *H2AFY2* | 9.268204 | 6.569005 | 2.944277 | 0.000000 |
| 10942 | *PRSS21* | 9.140530 | 4.672445 | 0.000000 | 0.000000 |
| 79070 | *KDELC1* | 9.125915 | 4.784205 | 0.000000 | 0.000000 |
| 9715 | *KIAA0773* | 9.100530 | 3.520641 | 0.000000 | 0.000000 |
| 22986 | *SORCS3* | 9.082493 | 3.201768 | 0.000000 | 0.000000 |
| 10850 | *CCL27* | 9.034405 | 4.655291 | 0.275346 | 0.000000 |
| 976 | *CD97* | 8.968839 | 8.275178 | 4.259477 | 0.194922 |
| 56895 | *AGPAT4* | 8.952689 | 5.442419 | 0.000000 | 0.000000 |
| 2770 | *GNAI1* | 8.945998 | 4.898433 | 0.426735 | 0.000000 |
| 5367 | *PMCH* | 8.942874 | 4.379153 | 0.000000 | 0.000000 |
| 2921 | *CXCL3* | 8.849623 | 5.258226 | 1.394194 | 0.000000 |
| 1809 | *DPYSL3* | 8.829213 | 5.363827 | 1.140532 | 0.000000 |
| 7441 | *VPREB1* | 8.744751 | 3.566288 | 0.000000 | 0.000000 |
| 23414 | *ZFPM2* | 8.735792 | 3.001373 | 0.000000 | 0.000000 |
| 81575 | *APOLD1* | 8.698523 | 5.012082 | 0.000000 | 0.000000 |
| 5790 | *PTPRCAP* | 8.697905 | 5.296362 | 1.317561 | 0.000000 |
| 51149 | *LOC51149* | 8.622240 | 4.711901 | 0.000000 | 0.000000 |
| 79057 | *PRRG3* | 8.610908 | 3.569247 | 0.000000 | 0.000000 |
| 3556 | *IL1RAP* | 8.549695 | 4.916280 | 1.165786 | 0.000000 |
| 4072 | *TACSTD1* | 8.519568 | 7.270650 | 8.911170 | 7.754502 |
| 223117 | *SEMA3D* | 8.488704 | 4.351951 | 0.928409 | 0.000000 |
| 947 | *CD34* | 8.460900 | 3.496274 | 0.000000 | 0.000000 |
| 23439 | *ATP1B4* | 8.362224 | 3.399387 | 0.582869 | 0.000000 |
| 10626 | *TRIM16* | 8.329653 | 4.682934 | 0.000000 | 0.000000 |
| 84719 | *MGC5457* | 8.296845 | 3.256643 | 0.000000 | 0.000000 |
| 3592 | *IL12A* | 8.246255 | 5.153265 | 1.150685 | 0.000000 |
| 7293 | *TNFRSF4* | 8.202653 | 5.164945 | 0.916798 | 0.000000 |
| 57523 | *KIAA1305* | 8.151407 | 5.585308 | 0.905816 | 0.000000 |
| 5152 | *PDE9A* | 8.135955 | 3.546912 | 0.000000 | 0.000000 |
| 56521 | *DNAJC12* | 8.051336 | 4.261985 | 0.000000 | 0.000000 |

| | | | | | |
|---|---|---|---|---|---|
| 9365 | *KL* | 8.041794 | 2.681672 | 0.000000 | 0.000000 |
| 481 | *ATP1B1* | 8.039977 | 3.246107 | 0.000000 | 0.000000 |
| 55353 | *LAPTM4B* | 8.028044 | 1.987717 | 0.000000 | 0.000000 |
| 3355 | *HTR1F* | 8.023521 | 5.547827 | 2.739964 | 0.000000 |
| 4071 | *TM4SF1* | 7.997031 | 3.726323 | 0.000000 | 0.000000 |
| 79864 | *C11orf63* | 7.962055 | 3.777462 | 0.000000 | 0.000000 |
| 3655 | *ITGA6* | 7.960003 | 3.248299 | 0.169990 | 0.000000 |
| 3397 | *ID1* | 7.918514 | 3.557412 | 0.000000 | 0.000000 |
| 5243 | *ABCB1* | 7.909534 | 2.625400 | 0.000000 | 0.000000 |
| 64754 | *SMYD3* | 7.908919 | 7.762218 | 4.929112 | 1.846062 |
| 8853 | *DDEF2* | 7.903859 | 3.965729 | 0.000000 | 0.000000 |
| 696 | *BTN1A1* | 7.816712 | 4.406536 | 2.758146 | 0.000000 |
| 55786 | *ZNF415* | 7.734692 | 1.902030 | 0.000000 | 0.000000 |
| 8800 | *PEX11A* | 7.713229 | 3.197269 | 0.000000 | 0.000000 |
| 8707 | *B3GALT2* | 7.709676 | 3.215779 | 0.000000 | 0.000000 |
| 2218 | *FCMD* | 7.681298 | 3.739868 | 0.000000 | 0.000000 |
| 1015 | *CDH17* | 7.649188 | 3.752136 | 0.000000 | 0.000000 |
| 1793 | *DOCK1* | 7.583369 | 4.911108 | 1.129780 | 0.000000 |
| 55296 | *TBC1D19* | 7.409842 | 3.586790 | 0.090012 | 0.000000 |
| 147172 | *DKFZp667M2411* | 7.406757 | 2.394008 | 0.000000 | 0.000000 |
| 150 | *ADRA2A* | 7.242067 | 2.339504 | 0.000000 | 0.000000 |
| 79720 | *VPS37B* | 7.240034 | 4.216800 | 0.605174 | 0.000000 |
| 7798 | *LUZP1* | 7.223716 | 3.725216 | 0.000000 | 0.000000 |
| 63920 | *LOC63920* | 7.214116 | 1.974504 | 0.000000 | 0.000000 |
| 8821 | *INPP4B* | 7.209603 | 1.782289 | 0.000000 | 0.000000 |
| 10243 | *GPHN* | 7.170965 | 2.180389 | 0.000000 | 0.000000 |
| 7177 | *TPSAB1* | 7.119765 | 2.714554 | 0.000000 | 0.000000 |
| 26105 | *DKFZP434C153* | 7.056197 | 1.900854 | 0.000000 | 0.000000 |
| 51152 | *LOC51152* | 7.049920 | 1.758481 | 0.000000 | 0.000000 |
| 29044 | *C9orf38* | 7.046479 | 0.992257 | 0.000000 | 0.000000 |
| 1950 | *EGF* | 7.041090 | 4.395837 | 3.264221 | 0.106607 |
| 79946 | *C10orf95* | 7.024970 | 4.110997 | 0.000000 | 0.000000 |
| 7294 | *TXK* | 7.015178 | 2.167726 | 0.000000 | 0.000000 |
| 1789 | *DNMT3B* | 6.970701 | 3.294466 | 0.000000 | 0.000000 |
| 79642 | *ARSJ* | 6.922933 | 1.718963 | 0.000000 | 0.000000 |
| 2913 | *GRM3* | 6.917015 | 3.037839 | 0.000000 | 0.000000 |
| 2150 | *F2RL1* | 6.913784 | 2.428295 | 0.000000 | 0.000000 |
| 8407 | *TAGLN2* | 6.906917 | 4.704419 | 0.000000 | 0.000000 |
| 1310 | *COL19A1* | 6.731083 | 1.677538 | 0.000000 | 0.000000 |
| 25827 | *FBXL2* | 6.729082 | 2.073670 | 0.000000 | 0.000000 |
| 71 | *ACTG1* | 6.712214 | 6.801710 | 1.512311 | 0.123055 |
| 79703 | *FLJ22531* | 6.673088 | 1.111002 | 0.000000 | 0.000000 |
| 284244 | *LOC284244* | 6.660942 | 1.996857 | 0.000000 | 0.000000 |
| 1938 | *EEF2* | 6.656234 | 4.945749 | 0.770219 | 0.000000 |
| 23209 | *MLC1* | 6.652070 | 2.591210 | 0.000000 | 0.000000 |
| 23251 | *KIAA1024* | 6.634311 | 2.004602 | 0.000000 | 0.000000 |
| 3202 | *HOXA5* | 6.613599 | 2.279998 | 0.000000 | 0.000000 |
| 5157 | *PDGFRL* | 6.602714 | 2.115939 | 0.000000 | 0.000000 |

| | | | | | |
|---|---|---|---|---|---|
| 81697 | *OR2B2* | 6.594805 | 2.810811 | 0.000000 | 0.000000 |
| 3709 | *ITPR2* | 6.582148 | 1.929176 | 0.000000 | 0.000000 |
| 28992 | *LRP16* | 6.536931 | 2.353453 | 0.000000 | 0.000000 |
| 4430 | *MYO1B* | 6.526522 | 2.503704 | 0.000000 | 0.000000 |
| 5645 | *PRSS2* | 6.462183 | 4.012959 | 1.149444 | 0.000000 |
| 9955 | *HS3ST3A1* | 6.441698 | 1.415588 | 0.000000 | 0.000000 |
| 3399 | *ID3* | 6.391998 | 0.036892 | 0.000000 | 0.000000 |
| 7035 | *TFPI* | 6.326482 | 2.346496 | 0.000000 | 0.000000 |
| 5187 | *PER1* | 6.307447 | 3.669063 | 1.474017 | 0.000000 |
| 26872 | *STEAP1* | 6.301468 | 0.501689 | 0.000000 | 0.000000 |
| 10562 | *OLFM4* | 6.299090 | 0.505114 | 0.000000 | 0.000000 |
| 9331 | *B4GALT6* | 6.294708 | 1.802741 | 0.000000 | 0.000000 |
| 57051 | *NAG18* | 6.259762 | 0.125647 | 0.000000 | 0.000000 |
| 3574 | *IL7* | 6.253506 | 1.490667 | 0.000000 | 0.000000 |
| 65982 | *ZNF447* | 6.246081 | 1.049397 | 0.000000 | 0.000000 |
| 5350 | *PLN* | 6.178631 | 1.474488 | 0.000000 | 0.000000 |
| 9723 | *SEMA3E* | 6.161991 | 0.774566 | 0.000000 | 0.000000 |
| 100 | *ADA* | 6.157616 | 4.498335 | 1.340551 | 0.000000 |
| 5357 | *PLS1* | 6.110484 | 3.930984 | 0.000000 | 0.000000 |
| 57084 | *SLC17A6* | 6.048081 | 1.206406 | 0.000000 | 0.000000 |
| 27018 | *NGFRAP1* | 6.041263 | 2.378105 | 0.000000 | 0.000000 |
| 25825 | *BACE2* | 6.021251 | 3.592779 | 1.201943 | 0.000000 |
| 2887 | *GRB10* | 5.994282 | 1.910106 | 0.000000 | 0.000000 |
| 54507 | *ADAMTSL4* | 5.903042 | 1.708381 | 0.000000 | 0.000000 |
| 10216 | *PRG4* | 5.901083 | 1.196170 | 0.000000 | 0.000000 |
| 4058 | *LTK* | 5.843981 | 1.718109 | 0.000000 | 0.000000 |
| 10203 | *CALCRL* | 5.813085 | 1.784346 | 0.000000 | 0.000000 |
| 65250 | *FLJ13231* | 5.796539 | 0.278500 | 0.000000 | 0.000000 |
| 23764 | *MAFF* | 5.685726 | 3.737988 | 0.982784 | 0.000000 |
| 8139 | *GAN* | 5.677625 | 1.098573 | 0.000000 | 0.000000 |
| 11030 | *RBPMS* | 5.604109 | 1.827282 | 0.000000 | 0.000000 |
| 10152 | *ABI2* | 5.586779 | 1.649292 | 0.000000 | 0.000000 |
| 51554 | *CCRL1* | 5.509546 | 0.812951 | 0.000000 | 0.000000 |
| 50859 | *SPOCK3* | 5.500200 | 0.010672 | 0.000000 | 0.000000 |
| 5834 | *PYGB* | 5.486600 | 2.134129 | 0.000000 | 0.000000 |
| 10391 | *CORO2B* | 5.465187 | 0.000000 | 0.000000 | 0.000000 |
| 10215 | *OLIG2* | 5.455273 | 1.120459 | 0.000000 | 0.000000 |
| 613 | *BCR* | 5.433243 | 2.057473 | 0.000000 | 0.000000 |
| 8645 | *KCNK5* | 5.417123 | 1.103927 | 0.000000 | 0.000000 |
| 10748 | *KLRA1* | 5.411862 | 0.120273 | 0.000000 | 0.000000 |
| 2037 | *EPB41L2* | 5.394966 | 2.356770 | 0.000000 | 0.000000 |
| 81796 | *SLCO5A1* | 5.340362 | 0.000000 | 0.000000 | 0.000000 |
| 23635 | *SSBP2* | 5.333010 | 2.769168 | 0.000000 | 0.000000 |
| 23683 | *PRKD3* | 5.319921 | 0.000000 | 0.000000 | 0.000000 |
| 81839 | *VANGL1* | 5.314157 | 0.974000 | 0.000000 | 0.000000 |
| 81616 | *ACSBG2* | 5.311469 | 0.000000 | 0.000000 | 0.000000 |
| 856 | *CATR1* | 5.299223 | 0.417065 | 0.000000 | 0.000000 |
| 858 | *CAV2* | 5.298308 | 0.376522 | 0.000000 | 0.000000 |

| | | | | | |
|---|---|---|---|---|---|
| 9136 | *RRP9* | 5.285973 | 1.173129 | 0.000000 | 0.000000 |
| 4222 | *MEOX1* | 5.284926 | 1.375575 | 0.000000 | 0.000000 |
| 80052 | *FLJ12331* | 5.281238 | 0.174496 | 0.000000 | 0.000000 |
| 54558 | *SPATA6* | 5.277550 | 1.230032 | 0.000000 | 0.000000 |
| 54810 | *GIPC2* | 5.245139 | 1.240044 | 0.000000 | 0.000000 |
| 79640 | *CTA-216E10.6* | 5.216216 | 1.004046 | 0.000000 | 0.000000 |
| 10321 | *CRISP3* | 5.203484 | 0.569797 | 0.000000 | 0.000000 |
| 26053 | *AUTS2* | 5.198790 | 0.000000 | 0.000000 | 0.000000 |
| 4363 | *ABCC1* | 5.188603 | 0.671468 | 0.000000 | 0.000000 |
| 8718 | *TNFRSF25* | 5.178139 | 1.658996 | 0.000000 | 0.000000 |
| 9780 | *FAM38A* | 5.168903 | 2.760751 | 0.000000 | 0.000000 |
| 26051 | *PPP1R16B* | 5.111586 | 2.048375 | 0.000000 | 0.000000 |
| 1193 | *CLIC2* | 5.096089 | 1.477652 | 0.000000 | 0.000000 |
| 2254 | *FGF9* | 5.075642 | 1.123981 | 0.000000 | 0.000000 |
| 6935 | *TCF8* | 5.046677 | 0.024098 | 0.000000 | 0.000000 |
| 54941 | *RNF125* | 5.016329 | 2.844465 | 0.062352 | 0.000000 |
| 3741 | *KCNA5* | 5.014475 | 0.153667 | 0.000000 | 0.000000 |
| 3730 | *KAL1* | 5.004040 | 2.689025 | 2.273026 | 1.032519 |
| 55068 | *RP11-301I17.1* | 4.968780 | 0.929976 | 0.000000 | 0.000000 |
| 2568 | *GABRP* | 4.965896 | 0.000000 | 0.000000 | 0.000000 |
| 9729 | *KIAA0408* | 4.950404 | 0.082824 | 0.000000 | 0.000000 |
| 7757 | *ZNF208* | 4.930981 | 0.233671 | 0.000000 | 0.000000 |
| 7525 | *YES1* | 4.928595 | 0.425394 | 0.000000 | 0.000000 |
| 51161 | *C3orf18* | 4.921851 | 0.115231 | 0.000000 | 0.000000 |
| 7367 | *UGT2B17* | 4.915942 | 0.000000 | 0.000000 | 0.000000 |
| 79986 | *ZNF702* | 4.902431 | 0.687961 | 0.000000 | 0.000000 |
| 155400 | *NSUN5B* | 4.872869 | 1.498722 | 0.000000 | 0.000000 |
| 51314 | *TXNDC3* | 4.872410 | 1.281863 | 0.000000 | 0.000000 |
| 55510 | *DDX43* | 4.863516 | 1.118343 | 0.000000 | 0.000000 |
| 6624 | *FSCN1* | 4.836405 | 1.520808 | 0.000000 | 0.000000 |
| 29922 | *NME7* | 4.833658 | 1.171683 | 0.000000 | 0.000000 |
| 79683 | *ZDHHC14* | 4.831207 | 0.715845 | 0.000000 | 0.000000 |
| 23090 | *ZNF423* | 4.821179 | 0.725644 | 0.000000 | 0.000000 |
| 55800 | *SCN3B* | 4.818801 | 0.000000 | 0.000000 | 0.000000 |
| 2827 | *GPR3* | 4.818564 | 0.109788 | 0.000000 | 0.000000 |
| 638 | *BIK* | 4.790911 | 0.595563 | 0.000000 | 0.000000 |
| 26031 | *OSBPL3* | 4.750514 | 1.104486 | 0.000000 | 0.000000 |
| 79154 | *MGC4172* | 4.747805 | 0.832396 | 0.000000 | 0.000000 |
| 26229 | *B3GAT3* | 4.745601 | 1.758254 | 0.000000 | 0.000000 |
| 29967 | *LRP12* | 4.741880 | 0.000000 | 0.000000 | 0.000000 |
| 8349 | *HIST2H2BE* | 4.741447 | 1.213156 | 0.000000 | 0.000000 |
| 5565 | *PRKAB2* | 4.719073 | 0.938565 | 0.000000 | 0.000000 |
| 3543 | *IGLL1* | 4.707672 | 1.405993 | 0.000000 | 0.000000 |
| 79698 | *ZMAT4* | 4.700822 | 0.000000 | 0.000000 | 0.000000 |
| 54885 | *TBC1D8B* | 4.683038 | 1.126605 | 0.000000 | 0.000000 |
| 3852 | *KRT5* | 4.651291 | 0.194808 | 0.000000 | 0.000000 |
| 79583 | *FLJ22167* | 4.650532 | 0.846728 | 0.000000 | 0.000000 |
| 91355 | *LRP5L* | 4.643473 | 0.622742 | 0.000000 | 0.000000 |

| | | | | | |
|---|---|---|---|---|---|
| 8544 | *PIR* | 4.643083 | 0.000000 | 0.000000 | 0.000000 |
| 59348 | *ZNF350* | 4.635838 | 0.043629 | 0.000000 | 0.000000 |
| 7349 | *UCN* | 4.624483 | 0.000000 | 0.000000 | 0.000000 |
| 730051 | *LOC730051* | 4.618974 | 0.070440 | 0.000000 | 0.000000 |
| 799 | *CALCR* | 4.614602 | 0.000000 | 0.000000 | 0.000000 |
| 8820 | *HESX1* | 4.599633 | 0.931421 | 0.000000 | 0.000000 |
| 2796 | *GNRH1* | 4.584809 | 0.375147 | 0.000000 | 0.000000 |
| 7424 | *VEGFC* | 4.577640 | 0.355095 | 0.000000 | 0.000000 |
| 29091 | *STXBP6* | 4.573797 | 0.769390 | 0.000000 | 0.000000 |
| 54996 | *MOSC2* | 4.564713 | 1.438971 | 0.000000 | 0.000000 |
| 3950 | *LECT2* | 4.537332 | 0.000000 | 0.000000 | 0.000000 |
| 1558 | *CYP2C8* | 4.526771 | 0.000000 | 0.000000 | 0.000000 |
| 22866 | *CNKSR2* | 4.525802 | 0.000000 | 0.000000 | 0.000000 |
| 2731 | *GLDC* | 4.494752 | 0.164259 | 0.000000 | 0.000000 |
| 3386 | *ICAM4* | 4.461442 | 1.289638 | 0.000000 | 0.000000 |
| 644450 | *LOC644450* | 4.461008 | 0.000000 | 0.000000 | 0.000000 |
| 1359 | *CPA3* | 4.459210 | 0.297255 | 0.000000 | 0.000000 |
| 54986 | *ULK4* | 4.438933 | 0.211678 | 0.000000 | 0.000000 |
| 1761 | *DMRT1* | 4.381109 | 0.000000 | 0.000000 | 0.000000 |
| 6414 | *SEPP1* | 4.376372 | 0.913514 | 0.000000 | 0.000000 |
| 9256 | *BZRAP1* | 4.359761 | 2.153587 | 0.000000 | 0.000000 |
| 81563 | *C1orf21* | 4.333951 | 0.000000 | 0.000000 | 0.000000 |
| 91752 | *ZNF804A* | 4.312007 | 0.000000 | 0.000000 | 0.000000 |
| 9705 | *ST18* | 4.297625 | 0.000000 | 0.000000 | 0.000000 |
| 894 | *CCND2* | 4.273820 | 0.000000 | 0.000000 | 0.000000 |
| 152098 | *IGHA1* | 4.268483 | 0.000000 | 0.000000 | 0.000000 |
| 10653 | *SPINT2* | 4.215644 | 2.344982 | 0.000000 | 0.000000 |
| 7915 | *ALDH5A1* | 4.209156 | 0.202542 | 0.000000 | 0.000000 |
| 10244 | *RABEPK* | 4.205633 | 0.771149 | 0.000000 | 0.000000 |
| 216 | *ALDH1A1* | 4.195046 | 1.109703 | 0.000000 | 0.000000 |
| 2059 | *EPS8* | 4.180383 | 1.132157 | 0.000000 | 0.000000 |
| 7318 | *UBE1L* | 4.165010 | 2.304429 | 0.666267 | 0.000000 |
| 2165 | *F13B* | 4.142915 | 0.000000 | 0.000000 | 0.000000 |
| 9720 | *KIAA0565* | 4.126548 | 0.000000 | 0.000000 | 0.000000 |
| 79789 | *CLMN* | 4.112149 | 0.000000 | 0.000000 | 0.000000 |
| 4145 | *MATK* | 4.104639 | 0.807861 | 0.000000 | 0.000000 |
| 1429 | *CRYZ* | 4.102173 | 0.383834 | 0.000000 | 0.000000 |
| 154796 | *AMOT* | 4.094222 | 0.000000 | 0.000000 | 0.000000 |
| 51299 | *NRN1* | 4.083962 | 0.179017 | 0.000000 | 0.000000 |
| 22835 | *ZFP30* | 4.058048 | 0.000000 | 0.000000 | 0.000000 |
| 303 | *ANXA2P1* | 4.037021 | 0.000000 | 0.000000 | 0.000000 |
| 55063 | *ZCWPW1* | 4.025692 | 0.914877 | 0.000000 | 0.000000 |
| 63895 | *FAM38B* | 3.994119 | 0.000000 | 0.000000 | 0.000000 |
| 4784 | *NFIX* | 3.982301 | 0.458157 | 0.000000 | 0.000000 |
| 23619 | *ZIM2* | 3.979682 | 0.293966 | 0.000000 | 0.000000 |
| 23621 | *BACE1* | 3.976477 | 0.629813 | 0.000000 | 0.000000 |
| 25940 | *FAM98A* | 3.971631 | 0.000000 | 0.000000 | 0.000000 |
| 22802 | *CLCA4* | 3.967587 | 0.000000 | 0.000000 | 0.000000 |

| | | | | | |
|---|---|---|---|---|---|
| 54621 | *FLJ20674* | 3.942938 | 0.337862 | 0.000000 | 0.000000 |
| 9618 | *TRAF4* | 3.904328 | 0.185897 | 0.000000 | 0.000000 |
| 9022 | *CLIC3* | 3.895404 | 0.011316 | 0.000000 | 0.000000 |
| 5276 | *SERPINI2* | 3.878718 | 0.000000 | 0.000000 | 0.000000 |
| 5638 | *PRRG1* | 3.869657 | 0.000000 | 0.000000 | 0.000000 |
| 9934 | *P2RY14* | 3.862337 | 0.000000 | 0.000000 | 0.000000 |
| 6522 | *SLC4A2* | 3.855737 | 0.487217 | 0.000000 | 0.000000 |
| 5644 | *PRSS1* | 3.851136 | 0.000000 | 0.000000 | 0.000000 |
| 389 | *RHOC* | 3.845160 | 1.005801 | 0.000000 | 0.000000 |
| 7423 | *VEGFB* | 3.824997 | 0.647214 | 0.000000 | 0.000000 |
| 10279 | *PRSS16* | 3.803339 | 0.000000 | 0.000000 | 0.000000 |
| 6691 | *SPINK2* | 3.793963 | 5.043591 | 5.845169 | 3.599097 |
| 3216 | *HOXB6* | 3.790153 | 0.000000 | 0.000000 | 0.000000 |
| 594 | *BCKDHB* | 3.783643 | 0.013204 | 0.000000 | 0.000000 |
| 645644 | *FLJ42627* | 3.783279 | 0.316379 | 0.000000 | 0.000000 |
| 126231 | *ZNF573* | 3.775819 | 0.000000 | 0.000000 | 0.000000 |
| 50810 | *HDGFRP3* | 3.774735 | 0.000000 | 0.000000 | 0.000000 |
| 57824 | *HMHB1* | 3.744978 | 0.000000 | 0.000000 | 0.000000 |
| 83442 | *SH3BGRL3* | 3.738963 | 3.507651 | 0.000000 | 0.000000 |
| 64595 | *TTTY15* | 3.732661 | 4.321210 | 15.430436 | 18.741931 |
| 8323 | *FZD6* | 3.707812 | 0.000000 | 0.000000 | 0.000000 |
| 3487 | *IGFBP4* | 3.691227 | 0.000000 | 0.000000 | 0.000000 |
| 84900 | *TMEM118* | 3.671923 | 0.496131 | 0.000000 | 0.000000 |
| 63910 | *C20orf59* | 3.660250 | 0.253606 | 0.000000 | 0.000000 |
| 8763 | *CD164* | 3.650665 | 0.908288 | 0.000000 | 0.000000 |
| 51559 | *NT5DC3* | 3.605565 | 0.062562 | 0.000000 | 0.000000 |
| 3769 | *KCNJ13* | 3.605135 | 0.000000 | 0.000000 | 0.000000 |
| 317762 | *C14orf65* | 3.594190 | 0.000000 | 0.000000 | 0.000000 |
| 79608 | *RIC3* | 3.587041 | 0.000000 | 0.000000 | 0.000000 |
| 57198 | *ATP8B2* | 3.583078 | 0.775781 | 0.000000 | 0.000000 |
| 56271 | *BEXL1* | 3.579263 | 1.727583 | 0.000000 | 0.000000 |
| 26750 | *RPS6KC1* | 3.571784 | 0.562719 | 0.000000 | 0.000000 |
| 152185 | *CCDC52* | 3.570058 | 0.000000 | 0.000000 | 0.000000 |
| 6674 | *SPAG1* | 3.540765 | 1.138825 | 0.000000 | 0.000000 |
| 5991 | *RFX3* | 3.528493 | 0.386213 | 0.000000 | 0.000000 |
| 5797 | *PTPRM* | 3.509328 | 0.000000 | 0.000000 | 0.000000 |
| 2920 | *CXCL2* | 3.497867 | 0.000000 | 0.000000 | 0.000000 |
| 26952 | *SMR3A* | 3.491228 | 0.000000 | 0.000000 | 0.000000 |
| 85368 | *KIAA1654* | 3.478153 | 0.000000 | 0.000000 | 0.000000 |
| 5174 | *PDZK1* | 3.465822 | 0.000000 | 0.000000 | 0.000000 |
| 3911 | *LAMA5* | 3.462259 | 0.000000 | 0.000000 | 0.000000 |
| 55259 | *CASC1* | 3.460365 | 0.000000 | 0.000000 | 0.000000 |
| 137872 | *ADHFE1* | 3.426692 | 0.038472 | 0.000000 | 0.000000 |
| 56892 | *C8orf4* | 3.418696 | 0.000000 | 0.000000 | 0.000000 |
| 11238 | *CA5B* | 3.390824 | 0.000000 | 0.000000 | 0.000000 |
| 79977 | *GRHL2* | 3.383178 | 0.000000 | 0.000000 | 0.000000 |
| 388567 | *ZNF749* | 3.376615 | 0.000000 | 0.000000 | 0.000000 |
| 7043 | *TGFB3* | 3.368018 | 0.000000 | 0.000000 | 0.000000 |

| | | | | | |
|---|---|---|---|---|---|
| 81789 | *TIGD6* | 3.366280 | 0.000000 | 0.000000 | 0.000000 |
| 79973 | *ZNF442* | 3.363156 | 0.000000 | 0.000000 | 0.000000 |
| 10322 | *SMYD5* | 3.336522 | 0.706844 | 0.000000 | 0.000000 |
| 8292 | *COLQ* | 3.290911 | 0.000000 | 0.000000 | 0.000000 |
| 147657 | *ZNF480* | 3.287087 | 0.000000 | 0.000000 | 0.000000 |
| 57326 | *PBXIP1* | 3.284397 | 0.000000 | 0.000000 | 0.000000 |
| 26035 | *GLCE* | 3.239485 | 0.000000 | 0.000000 | 0.000000 |
| 80117 | *ARL14* | 3.226529 | 0.000000 | 0.000000 | 0.000000 |
| 1244 | *ABCC2* | 3.221160 | 0.000000 | 0.000000 | 0.000000 |
| 8204 | *NRIP1* | 3.199634 | 0.579021 | 0.000000 | 0.000000 |
| 3576 | *IL8* | 3.197564 | 2.030299 | 0.000000 | 0.000000 |
| 9623 | *TCL1B* | 3.196931 | 0.000000 | 0.000000 | 0.000000 |
| 23564 | *DDAH2* | 3.196630 | 1.260106 | 0.000000 | 0.000000 |
| 6296 | *ACSM3* | 3.181343 | 0.000000 | 0.000000 | 0.000000 |
| 55779 | *WDR52* | 3.149879 | 0.000000 | 0.000000 | 0.000000 |
| 11273 | *ATXN2L* | 3.129396 | 0.000000 | 0.000000 | 0.000000 |
| 6564 | *SLC15A1* | 3.115293 | 0.000000 | 0.000000 | 0.000000 |
| 23508 | *TTC9* | 3.114368 | 0.000000 | 0.000000 | 0.000000 |
| 2515 | *ADAM2* | 3.111162 | 0.000000 | 0.000000 | 0.000000 |
| 80152 | *CENPT* | 3.106144 | 0.000000 | 0.000000 | 0.000000 |
| 6575 | *SLC20A2* | 3.093622 | 0.000000 | 0.000000 | 0.000000 |
| 9532 | *BAG2* | 3.089848 | 0.000000 | 0.000000 | 0.000000 |
| 7728 | *ZNF175* | 3.068703 | 0.000000 | 0.000000 | 0.000000 |
| 260294 | *NSUN5C* | 3.064449 | 0.121488 | 0.000000 | 0.000000 |
| 1945 | *EFNA4* | 3.059910 | 0.000000 | 0.000000 | 0.000000 |
| 55073 | *FLJ10120* | 3.053398 | 0.000000 | 0.000000 | 0.000000 |
| 10608 | *MXD4* | 3.049176 | 0.269448 | 0.000000 | 0.000000 |
| 79809 | *TTC21B* | 3.024469 | 0.000000 | 0.000000 | 0.000000 |
| 113178 | *SCAMP4* | 3.011526 | 0.000000 | 0.000000 | 0.000000 |
| 80868 | *HCG4P6* | 3.010924 | 0.000000 | 0.000000 | 0.000000 |
| 7561 | *ZNF14* | 3.005763 | 0.000000 | 0.000000 | 0.000000 |
| 441024 | *MTHFD2L* | 3.004234 | 0.000000 | 0.000000 | 0.000000 |
| 196 | *AHR* | 2.991171 | 1.387382 | 0.000000 | 0.000000 |
| 5616 | *PRKY* | 2.981993 | 0.970863 | 1.724291 | 0.846891 |
| 23284 | *LPHN3* | 2.979303 | 0.000000 | 0.000000 | 0.000000 |
| 55092 | *TMEM51* | 2.975696 | 0.000000 | 0.000000 | 0.000000 |
| 4313 | *MMP2* | 2.969265 | 0.000000 | 0.000000 | 0.000000 |
| 5793 | *PTPRG* | 2.964438 | 0.000000 | 0.000000 | 0.000000 |
| 27130 | *INVS* | 2.960717 | 0.000000 | 0.000000 | 0.000000 |
| 8581 | *LY6D* | 2.956474 | 0.000000 | 0.000000 | 0.000000 |
| 80212 | *CCDC92* | 2.927247 | 0.000000 | 0.000000 | 0.000000 |
| 79143 | *LENG4* | 2.924069 | 0.586005 | 0.000000 | 0.000000 |
| 2983 | *GUCY1B3* | 2.920002 | 0.000000 | 0.000000 | 0.000000 |
| 23322 | *KIAA1005* | 2.905886 | 0.000000 | 0.000000 | 0.000000 |
| 55277 | *FLJ10986* | 2.899839 | 0.000000 | 0.000000 | 0.000000 |
| 857 | *CAV1* | 2.890977 | 0.000000 | 0.000000 | 0.000000 |
| 9945 | *GFPT2* | 2.881805 | 0.000000 | 0.000000 | 0.000000 |
| 54457 | *TAF7L* | 2.855680 | 0.000000 | 0.000000 | 0.000000 |

| | | | | | |
|---|---|---|---|---|---|
| 5168 | *ENPP2* | 2.852433 | 0.000000 | 0.000000 | 0.000000 |
| 1038 | *CDR1* | 2.846525 | 0.000000 | 0.000000 | 0.000000 |
| 126070 | *ZNF440* | 2.833368 | 0.000000 | 0.000000 | 0.000000 |
| 57579 | *KIAA1411* | 2.827417 | 0.000000 | 0.000000 | 0.000000 |
| 228 | *ALDOAP2* | 2.826921 | 0.000000 | 0.000000 | 0.000000 |
| 29935 | *RPA4* | 2.820457 | 0.000000 | 0.000000 | 0.000000 |
| 9172 | *MYOM2* | 2.794114 | 0.000000 | 0.000000 | 0.000000 |
| 2845 | *GPR22* | 2.792783 | 0.000000 | 0.000000 | 0.000000 |
| 79611 | *FLJ21963* | 2.774706 | 0.000000 | 0.000000 | 0.000000 |
| 54622 | *ARL15* | 2.761455 | 0.000000 | 0.000000 | 0.000000 |
| 54662 | *TBC1D13* | 2.755103 | 0.086444 | 0.000000 | 0.000000 |
| 9170 | *EDG4* | 2.738271 | 0.000000 | 0.000000 | 0.000000 |
| 26575 | *RGS17* | 2.734961 | 0.000000 | 0.000000 | 0.000000 |
| 2294 | *FOXF1* | 2.719591 | 0.000000 | 0.000000 | 0.000000 |
| 1591 | *CYP24A1* | 2.691529 | 0.000000 | 0.000000 | 0.000000 |
| 2516 | *NR5A1* | 2.691114 | 0.000000 | 0.000000 | 0.000000 |
| 25780 | *RASGRP3* | 2.689911 | 0.000000 | 0.000000 | 0.000000 |
| 80255 | *SLC35F5* | 2.688316 | 0.000000 | 0.000000 | 0.000000 |
| 6285 | *S100B* | 2.685775 | 0.000000 | 0.000000 | 0.000000 |
| 9966 | *TNFSF15* | 2.666355 | 0.000000 | 0.000000 | 0.000000 |
| 80099 | *FLJ21075* | 2.666263 | 0.000000 | 0.000000 | 0.000000 |
| 9087 | *TMSB4Y* | 2.646962 | 0.155354 | 0.000000 | 0.000000 |
| 987 | *LRBA* | 2.637022 | 0.255945 | 0.000000 | 0.000000 |
| 904 | *CCNT1* | 2.629812 | 0.000000 | 0.000000 | 0.000000 |
| 59335 | *PRDM12* | 2.607304 | 0.000000 | 0.000000 | 0.000000 |
| 2162 | *F13A1* | 2.604718 | 0.000000 | 0.000000 | 0.000000 |
| 23138 | *N4BP3* | 2.593892 | 0.000000 | 0.000000 | 0.000000 |
| 5330 | *PLCB2* | 2.588669 | 0.000000 | 0.000000 | 0.000000 |
| 1787 | *TRDMT1* | 2.584654 | 0.000000 | 0.000000 | 0.000000 |
| 8787 | *RGS9* | 2.578545 | 0.000000 | 0.000000 | 0.000000 |
| 6450 | *SH3BGR* | 2.578456 | 0.000000 | 0.000000 | 0.000000 |
| 5998 | *RGS3* | 2.570832 | 0.000000 | 0.000000 | 0.000000 |
| 5646 | *PRSS3* | 2.562335 | 0.000000 | 0.000000 | 0.000000 |
| 1828 | *DSG1* | 2.553157 | 0.000000 | 0.000000 | 0.000000 |
| 394 | *ARHGAP5* | 2.536289 | 0.000000 | 0.000000 | 0.000000 |
| 55030 | *FBXO34* | 2.533400 | 0.000000 | 0.000000 | 0.000000 |
| 2241 | *FER* | 2.530328 | 0.000000 | 0.000000 | 0.000000 |
| 5763 | *PTMS* | 2.525169 | 0.000000 | 0.000000 | 0.000000 |
| 8854 | *ALDH1A2* | 2.519087 | 0.000000 | 0.000000 | 0.000000 |
| 4649 | *MYO9A* | 2.513479 | 0.000000 | 0.000000 | 0.000000 |
| 4808 | *NHLH2* | 2.510860 | 0.000000 | 0.000000 | 0.000000 |
| 6678 | *SPARC* | 2.498857 | 0.000000 | 0.000000 | 0.000000 |
| 7693 | *ZNF134* | 2.494784 | 0.000000 | 0.000000 | 0.000000 |
| 2015 | *EMR1* | 2.493159 | 0.000000 | 0.000000 | 0.000000 |
| 605 | *BCL7A* | 2.490750 | 0.000000 | 0.000000 | 0.000000 |
| 3680 | *ITGA9* | 2.490533 | 0.000000 | 0.000000 | 0.000000 |
| 657 | *BMPR1A* | 2.490032 | 0.000000 | 0.000000 | 0.000000 |
| 27330 | *RPS6KA6* | 2.487684 | 0.000000 | 0.000000 | 0.000000 |

| | | | | | |
|---|---|---|---|---|---|
| 7857 | *SCG2* | 2.480522 | 0.000000 | 0.000000 | 0.000000 |
| 10787 | *NCKAP1* | 2.478467 | 0.000000 | 0.000000 | 0.000000 |
| 367 | *AR* | 2.459779 | 0.000000 | 0.000000 | 0.000000 |
| 7718 | *ZNF165* | 2.459387 | 0.000000 | 0.000000 | 0.000000 |
| 79819 | *WDR78* | 2.458839 | 0.000000 | 0.000000 | 0.000000 |
| 118491 | *TTC18* | 2.444163 | 0.000000 | 0.000000 | 0.000000 |
| 9746 | *CLSTN3* | 2.439216 | 0.000000 | 0.000000 | 0.000000 |
| 64577 | *ALDH8A1* | 2.425182 | 0.000000 | 0.000000 | 0.000000 |
| 24138 | *IFIT5* | 2.382181 | 0.000000 | 0.000000 | 0.000000 |
| 64083 | *GOLPH3* | 2.376927 | 0.547517 | 0.000000 | 0.000000 |
| 6570 | *SLC18A1* | 2.345410 | 0.000000 | 0.000000 | 0.000000 |
| 10804 | *IGKC* | 2.335413 | 0.000000 | 0.000000 | 0.000000 |
| 55777 | *MBD5* | 2.334254 | 0.000000 | 0.000000 | 0.000000 |
| 54661 | *PCDHB17* | 2.322961 | 0.000000 | 0.000000 | 0.000000 |
| 5321 | *PLA2G4A* | 2.321953 | 0.000000 | 0.000000 | 0.000000 |
| 3087 | *HHEX* | 2.306967 | 0.000000 | 0.000000 | 0.000000 |
| 11250 | *GPR45* | 2.302369 | 0.000000 | 0.000000 | 0.000000 |
| 7404 | *UTY* | 2.301349 | 1.737227 | 9.501554 | 11.479673 |
| 26509 | *FER1L3* | 2.300189 | 0.000000 | 0.000000 | 0.000000 |
| 23221 | *RHOBTB2* | 2.295106 | 0.000000 | 0.000000 | 0.000000 |
| 202018 | *FLJ90013* | 2.272408 | 0.000000 | 0.000000 | 0.000000 |
| 90957 | *DHX57* | 2.265672 | 0.000000 | 0.000000 | 0.000000 |
| 3633 | *INPP5B* | 2.261962 | 0.000000 | 0.000000 | 0.000000 |
| 79828 | *METTL8* | 2.251128 | 0.000000 | 0.000000 | 0.000000 |
| 3670 | *ISL1* | 2.245597 | 0.000000 | 0.000000 | 0.000000 |
| 57228 | *LOC57228* | 2.230775 | 0.000000 | 0.000000 | 0.000000 |
| 51205 | *ACP6* | 2.230314 | 0.000000 | 0.000000 | 0.000000 |
| 7730 | *ZNF177* | 2.225524 | 0.000000 | 0.000000 | 0.000000 |
| 127703 | *FLJ38984* | 2.217717 | 0.000000 | 0.000000 | 0.000000 |
| 8558 | *CDK10* | 2.207604 | 0.000000 | 0.000000 | 0.000000 |
| 10896 | *OCLM* | 2.203800 | 0.000000 | 0.000000 | 0.000000 |
| 64787 | *EPS8L2* | 2.199246 | 0.000000 | 0.000000 | 0.000000 |
| 3995 | *FADS3* | 2.198226 | 0.000000 | 0.000000 | 0.000000 |
| 256949 | *RPS28* | 2.193702 | 0.000000 | 0.000000 | 0.000000 |
| 5342 | *PLGLB2* | 2.188973 | 0.000000 | 0.000000 | 0.000000 |
| 26032 | *SUSD5* | 2.180901 | 0.000000 | 0.000000 | 0.000000 |
| 3732 | *CD82* | 2.171006 | 0.389489 | 0.000000 | 0.000000 |
| 9499 | *MYOT* | 2.169974 | 0.000000 | 0.000000 | 0.000000 |
| 23060 | *ZNF609* | 2.166286 | 0.000000 | 0.000000 | 0.000000 |
| 150967 | *DKFZp434H1419* | 2.164947 | 0.000000 | 0.000000 | 0.000000 |
| 5095 | *PCCA* | 2.158584 | 0.000000 | 0.000000 | 0.000000 |
| 79719 | *FLJ11506* | 2.157900 | 0.000000 | 0.000000 | 0.000000 |
| 25992 | *SNED1* | 2.154052 | 0.000000 | 0.000000 | 0.000000 |
| 64859 | *OBFC2A* | 2.154033 | 0.000000 | 0.000000 | 0.000000 |
| 8658 | *TNKS* | 2.151881 | 0.000000 | 0.000000 | 0.000000 |
| 5585 | *PKN1* | 2.149184 | 0.000000 | 0.000000 | 0.000000 |
| 1847 | *DUSP5* | 2.134718 | 0.000000 | 0.000000 | 0.000000 |
| 8906 | *AP1G2* | 2.130041 | 0.335020 | 0.000000 | 0.000000 |

| | | | | | |
|---|---|---|---|---|---|
| 114044 | *MCM3APAS* | 2.129686 | 0.000000 | 0.000000 | 0.000000 |
| 5178 | *PEG3* | 2.124634 | 0.000000 | 0.000000 | 0.000000 |
| 55150 | *FLJ10490* | 2.109344 | 0.000000 | 0.000000 | 0.000000 |
| 9840 | *KIAA0748* | 2.096646 | 0.000000 | 0.000000 | 0.000000 |
| 2346 | *FOLH1* | 2.092567 | 0.000000 | 0.000000 | 0.000000 |
| 79904 | *FLJ11710* | 2.092190 | 0.000000 | 0.000000 | 0.000000 |
| 7799 | *PRDM2* | 2.078781 | 0.000000 | 0.000000 | 0.000000 |
| 51365 | *PLA1A* | 2.072672 | 0.000000 | 0.000000 | 0.000000 |
| 151230 | *KLHL23* | 2.065646 | 0.000000 | 0.000000 | 0.000000 |
| 27239 | *GPR162* | 2.062017 | 0.000000 | 0.000000 | 0.000000 |
| 5304 | *PIP* | 2.032198 | 0.000000 | 0.000000 | 0.000000 |
| 9108 | *MTMR7* | 2.031439 | 0.000000 | 0.000000 | 0.000000 |
| 55610 | *CCDC132* | 2.023376 | 0.000000 | 0.000000 | 0.000000 |
| 10497 | *UNC13B* | 2.018726 | 0.000000 | 0.000000 | 0.000000 |
| 55472 | *C8orf39* | 2.009006 | 0.000000 | 0.000000 | 0.000000 |
| 5567 | *PRKACB* | 1.999950 | 0.000000 | 0.000000 | 0.000000 |
| 9703 | *KIAA0100* | 1.986557 | 0.000000 | 0.000000 | 0.000000 |
| 10752 | *CHL1* | 1.983868 | 0.000000 | 0.000000 | 0.000000 |
| 9854 | *TMEM24* | 1.982934 | 0.000000 | 0.000000 | 0.000000 |
| 116828 | *CG030* | 1.978130 | 0.000000 | 0.000000 | 0.000000 |
| 7188 | *TRAF5* | 1.971735 | 0.000000 | 0.000000 | 0.000000 |
| 7479 | *WNT8B* | 1.970194 | 0.000000 | 0.000000 | 0.000000 |
| 10905 | *MAN1A2* | 1.964345 | 0.000000 | 0.000000 | 0.000000 |
| 9666 | *DZIP3* | 1.956953 | 0.000000 | 0.000000 | 0.000000 |
| 11223 | *MSTP9* | 1.948627 | 0.000000 | 0.000000 | 0.000000 |
| 23132 | *RAD54L2* | 1.945579 | 0.000000 | 0.000000 | 0.000000 |
| 4680 | *CEACAM6* | 1.944139 | 0.109510 | 0.000000 | 0.000000 |
| 79884 | *MAP9* | 1.934856 | 0.000000 | 0.000000 | 0.000000 |
| 55089 | *SLC38A4* | 1.929209 | 0.000000 | 0.000000 | 0.000000 |
| 3199 | *HOXA2* | 1.927482 | 0.000000 | 0.000000 | 0.000000 |
| 57473 | *GM632* | 1.923573 | 0.000000 | 0.000000 | 0.000000 |
| 2863 | *GPR39* | 1.921794 | 0.000000 | 0.000000 | 0.000000 |
| 26038 | *CHD5* | 1.907388 | 0.000000 | 0.000000 | 0.000000 |
| 55851 | *PSENEN* | 1.907351 | 0.000000 | 0.000000 | 0.000000 |
| 57728 | *WDR19* | 1.906992 | 0.000000 | 0.000000 | 0.000000 |
| 3092 | *HIP1* | 1.896949 | 0.000000 | 0.000000 | 0.000000 |
| 7252 | *TSHB* | 1.896604 | 0.000000 | 0.000000 | 0.000000 |
| 196743 | *PAOX* | 1.887434 | 0.000000 | 0.000000 | 0.000000 |
| 57720 | *GPR107* | 1.881258 | 0.000000 | 0.000000 | 0.000000 |
| 2529 | *FUT7* | 1.876921 | 0.000000 | 0.000000 | 0.000000 |
| 3696 | *ITGB8* | 1.867475 | 0.000000 | 0.000000 | 0.000000 |
| 5295 | *PIK3R1* | 1.860787 | 0.000000 | 0.000000 | 0.000000 |
| 9828 | *ARHGEF17* | 1.859009 | 0.000000 | 0.000000 | 0.000000 |
| 7673 | *ZNF222* | 1.858769 | 0.000000 | 0.000000 | 0.000000 |
| 10824 | *EPAG* | 1.853213 | 0.000000 | 0.000000 | 0.000000 |
| 4625 | *MYH7* | 1.838800 | 0.000000 | 0.000000 | 0.000000 |
| 90 | *ACVR1* | 1.833636 | 0.000000 | 0.000000 | 0.000000 |
| 2199 | *FBLN2* | 1.824230 | 0.000000 | 0.000000 | 0.000000 |

| | | | | | |
|---|---|---|---|---|---|
| 54332 | *GDAP1* | 1.823024 | 0.000000 | 0.000000 | 0.000000 |
| 26999 | *CYFIP2* | 1.816512 | 0.000000 | 0.000000 | 0.000000 |
| 22852 | *ANKRD26* | 1.807645 | 0.000000 | 0.000000 | 0.000000 |
| 8324 | *FZD7* | 1.805014 | 0.000000 | 0.000000 | 0.000000 |
| 79823 | *C2orf34* | 1.804257 | 0.000000 | 0.000000 | 0.000000 |
| 6925 | *TCF4* | 1.802251 | 0.000000 | 0.000000 | 0.000000 |
| 9892 | *SNAP91* | 1.789206 | 0.000000 | 0.000000 | 0.000000 |
| 6847 | *SYCP1* | 1.778453 | 0.000000 | 0.000000 | 0.000000 |
| 256227 | *MGC87042* | 1.767614 | 0.000000 | 0.000000 | 0.000000 |
| 4958 | *OMD* | 1.767199 | 0.000000 | 0.000000 | 0.000000 |
| 55526 | *DHTKD1* | 1.765774 | 0.000000 | 0.000000 | 0.000000 |
| 64167 | *LRAP* | 1.758764 | 0.000000 | 0.000000 | 0.000000 |
| 793 | *CALB1* | 1.746383 | 0.000000 | 0.000000 | 0.000000 |
| 55617 | *TASP1* | 1.736844 | 0.000000 | 0.000000 | 0.000000 |
| 79822 | *ARHGAP28* | 1.718074 | 0.000000 | 0.000000 | 0.000000 |
| 9230 | *RAB11B* | 1.709647 | 0.000000 | 0.000000 | 0.000000 |
| 7105 | *TSPAN6* | 1.698809 | 0.000000 | 0.000000 | 0.000000 |
| 9890 | *LPPR4* | 1.697537 | 0.000000 | 0.000000 | 0.000000 |
| 22871 | *NLGN1* | 1.688484 | 0.000000 | 0.000000 | 0.000000 |
| 3385 | *ICAM3* | 1.684317 | 0.425073 | 0.000000 | 0.000000 |
| 79800 | *ALS2CR8* | 1.678519 | 0.000000 | 0.000000 | 0.000000 |
| 2051 | *EPHB6* | 1.676929 | 0.000000 | 0.000000 | 0.000000 |
| 26027 | *ACOT11* | 1.675551 | 0.000000 | 0.000000 | 0.000000 |
| 3201 | *HOXA4* | 1.669135 | 0.000000 | 0.000000 | 0.000000 |
| 51177 | *PLEKHO1* | 1.668902 | 0.456908 | 0.000000 | 0.000000 |
| 1962 | *EHHADH* | 1.667525 | 0.000000 | 0.000000 | 0.000000 |
| 399 | *RHOH* | 1.663131 | 0.000000 | 0.000000 | 0.000000 |
| 29940 | *SART2* | 1.651897 | 0.000000 | 0.000000 | 0.000000 |
| 51134 | *CCDC41* | 1.643294 | 0.000000 | 0.000000 | 0.000000 |
| 642559 | *LOC642559* | 1.637745 | 0.000000 | 0.000000 | 0.000000 |
| 7704 | *ZBTB16* | 1.633607 | 0.000000 | 0.000000 | 0.000000 |
| 27039 | *PKD2L2* | 1.625831 | 0.000000 | 0.000000 | 0.000000 |
| 9824 | *ARHGAP11A* | 1.624899 | 0.000000 | 0.000000 | 0.000000 |
| 65082 | *VPS33A* | 1.622536 | 0.000000 | 0.000000 | 0.000000 |
| 5777 | *PTPN6* | 1.606754 | 0.000000 | 0.000000 | 0.000000 |
| 22837 | *COBLL1* | 1.578767 | 0.000000 | 0.000000 | 0.000000 |
| 10962 | *MLLT11* | 1.578563 | 0.000000 | 0.000000 | 0.000000 |
| 83990 | *BRIP1* | 1.576453 | 0.000000 | 0.000000 | 0.000000 |
| 114899 | *C1QTNF3* | 1.575280 | 0.000000 | 0.000000 | 0.000000 |
| 8350 | *HIST1H3A* | 1.557430 | 0.000000 | 0.000000 | 0.000000 |
| 765 | *CA6* | 1.539880 | 0.000000 | 0.000000 | 0.000000 |
| 2868 | *GRK4* | 1.537700 | 0.000000 | 0.000000 | 0.000000 |
| 53371 | *NUP54* | 1.536874 | 0.000000 | 0.000000 | 0.000000 |
| 55711 | *MLSTD1* | 1.535538 | 0.000000 | 0.000000 | 0.000000 |
| 7739 | *ZNF185* | 1.520660 | 0.698621 | 0.000000 | 0.000000 |
| 5444 | *PON1* | 1.520412 | 0.000000 | 0.000000 | 0.000000 |
| 51054 | *PLEKHA9* | 1.520326 | 0.000000 | 0.000000 | 0.000000 |
| 8542 | *APOL1* | 1.500456 | 0.000000 | 0.000000 | 0.000000 |

| | | | | | |
|---|---|---|---|---|---|
| 3221 | *HOXC4* | 1.499985 | 0.000000 | 0.000000 | 0.000000 |
| 9209 | *LRRFIP2* | 1.498926 | 0.000000 | 0.000000 | 0.000000 |
| 9823 | *ARMCX2* | 1.497907 | 0.000000 | 0.000000 | 0.000000 |
| 80221 | *FLJ20920* | 1.492929 | 0.000000 | 0.000000 | 0.000000 |
| 10331 | *B3GNT3* | 1.492538 | 0.000000 | 0.000000 | 0.000000 |
| 7171 | *TPM4* | 1.491696 | 0.000000 | 0.000000 | 0.000000 |
| 134728 | *IRAK1BP1* | 1.487889 | 0.000000 | 0.000000 | 0.000000 |
| 5289 | *PIK3C3* | 1.485107 | 0.000000 | 0.000000 | 0.000000 |
| 5774 | *PTPN3* | 1.483017 | 0.000000 | 0.000000 | 0.000000 |
| 23033 | *DOPEY1* | 1.482796 | 0.000000 | 0.000000 | 0.000000 |
| 5583 | *PRKCH* | 1.478958 | 0.000000 | 0.000000 | 0.000000 |
| 54477 | *PLEKHA5* | 1.465901 | 0.000000 | 0.000000 | 0.000000 |
| 3204 | *HOXA7* | 1.463620 | 0.000000 | 0.000000 | 0.000000 |
| 230 | *ALDOC* | 1.462756 | 0.000000 | 0.000000 | 0.000000 |
| 10090 | *UST* | 1.462402 | 0.000000 | 0.000000 | 0.000000 |
| 23201 | *KIAA0280* | 1.460027 | 0.000000 | 0.000000 | 0.000000 |
| 11127 | *KIF3A* | 1.457810 | 0.000000 | 0.000000 | 0.000000 |
| 9500 | *MAGED1* | 1.455772 | 0.463329 | 0.000000 | 0.000000 |
| 9727 | *RAB11FIP3* | 1.446464 | 0.000000 | 0.000000 | 0.000000 |
| 1021 | *CDK6* | 1.439250 | 0.000000 | 0.000000 | 0.000000 |
| 142 | *PARP1* | 1.433731 | 0.068201 | 0.000000 | 0.000000 |
| 2053 | *EPHX2* | 1.429994 | 0.000000 | 0.000000 | 0.000000 |
| 26984 | *SEC22A* | 1.413114 | 0.000000 | 0.000000 | 0.000000 |
| 26048 | *ZNF500* | 1.400862 | 0.000000 | 0.000000 | 0.000000 |
| 51351 | *H-plk* | 1.394730 | 0.000000 | 0.000000 | 0.000000 |
| 66035 | *SLC2A11* | 1.383136 | 0.000000 | 0.000000 | 0.000000 |
| 57091 | *C20orf32* | 1.379758 | 0.000000 | 0.000000 | 0.000000 |
| 5480 | *PPIC* | 1.369111 | 0.000000 | 0.000000 | 0.000000 |
| 7696 | *ZNF137* | 1.366738 | 0.000000 | 0.000000 | 0.000000 |
| 221830 | *TWISTNB* | 1.354937 | 0.000000 | 0.000000 | 0.000000 |
| 7975 | *MAFK* | 1.353916 | 0.000000 | 0.000000 | 0.000000 |
| 4261 | *CIITA* | 1.347443 | 0.000000 | 0.000000 | 0.000000 |
| 262 | *AMD1* | 1.343062 | 0.000000 | 0.000000 | 0.000000 |
| 3326 | *HSP90AB1* | 1.338861 | 0.000000 | 0.000000 | 0.000000 |
| 10723 | *SLC12A7* | 1.336075 | 0.533872 | 0.000000 | 0.000000 |
| 5900 | *RALGDS* | 1.327906 | 0.000000 | 0.000000 | 0.000000 |
| 54554 | *WDR5B* | 1.326439 | 0.000000 | 0.000000 | 0.000000 |
| 80755 | *AARSD1* | 1.320378 | 0.000000 | 0.000000 | 0.000000 |
| 10077 | *TSPAN32* | 1.316591 | 0.000000 | 0.000000 | 0.000000 |
| 930 | *CD19* | 1.311199 | 0.000000 | 0.000000 | 0.000000 |
| 7851 | *MALL* | 1.310824 | 0.000000 | 0.000000 | 0.000000 |
| 84796 | *MGC13053* | 1.309796 | 0.000000 | 0.000000 | 0.000000 |
| 89874 | *SLC25A21* | 1.307252 | 0.000000 | 0.000000 | 0.000000 |
| 10585 | *POMT1* | 1.303744 | 0.000000 | 0.000000 | 0.000000 |
| 11221 | *DUSP10* | 1.295019 | 0.000000 | 0.000000 | 0.000000 |
| 79600 | *FLJ21127* | 1.289404 | 0.000000 | 0.000000 | 0.000000 |
| 22902 | *RUFY3* | 1.286586 | 0.000000 | 0.000000 | 0.000000 |
| 85477 | *SCIN* | 1.284692 | 0.000000 | 0.000000 | 0.000000 |

| | | | | | |
|---|---|---|---|---|---|
| 5054 | *SERPINE1* | 1.282190 | 0.000000 | 0.000000 | 0.000000 |
| 395 | *ARHGAP6* | 1.279573 | 0.000000 | 0.000000 | 0.000000 |
| 23175 | *LPIN1* | 1.264174 | 0.000000 | 0.000000 | 0.000000 |
| 54923 | *LIME1* | 1.263769 | 0.000000 | 0.000000 | 0.000000 |
| 20 | *ABCA2* | 1.261733 | 0.000000 | 0.000000 | 0.000000 |
| 56605 | *ERO1LB* | 1.259814 | 0.000000 | 0.000000 | 0.000000 |
| 5190 | *PEX6* | 1.256236 | 0.000000 | 0.000000 | 0.000000 |
| 7556 | *ZNF10* | 1.254846 | 0.000000 | 0.000000 | 0.000000 |
| 51157 | *ZNF580* | 1.253091 | 0.000000 | 0.000000 | 0.000000 |
| 6252 | *RTN1* | 1.250354 | 0.000000 | 0.000000 | 0.000000 |
| 55752 | *11-Sep* | 1.246030 | 0.000000 | 0.000000 | 0.000000 |
| 63893 | *UBE2O* | 1.239778 | 0.000000 | 0.000000 | 0.000000 |
| 7157 | *TP53* | 1.236770 | 0.000000 | 0.000000 | 0.000000 |
| 10803 | *CCR9* | 1.234099 | 0.000000 | 0.000000 | 0.000000 |
| 79991 | *OBFC1* | 1.219634 | 0.000000 | 0.000000 | 0.000000 |
| 2747 | *GLUD2* | 1.218675 | 0.000000 | 0.000000 | 0.000000 |
| 22937 | *SCAP* | 1.216123 | 0.000000 | 0.000000 | 0.000000 |
| 5939 | *RBMS2* | 1.207704 | 0.000000 | 0.000000 | 0.000000 |
| 8526 | *DGKE* | 1.189114 | 0.000000 | 0.000000 | 0.000000 |
| 5978 | *REST* | 1.184814 | 0.000000 | 0.000000 | 0.000000 |
| 23180 | *RFTN1* | 1.180366 | 0.000000 | 0.000000 | 0.000000 |
| 2026 | *ENO2* | 1.176599 | 0.000000 | 0.000000 | 0.000000 |
| 10400 | *PEMT* | 1.170741 | 0.000000 | 0.000000 | 0.000000 |
| 8187 | *ZNF239* | 1.167476 | 0.000000 | 0.000000 | 0.000000 |
| 54793 | *KCTD9* | 1.164994 | 0.000000 | 0.000000 | 0.000000 |
| 7268 | *TTC4* | 1.162009 | 0.000000 | 0.000000 | 0.000000 |
| 10634 | *GAS2L1* | 1.161470 | 0.000000 | 0.000000 | 0.000000 |
| 85002 | *FAM86B1* | 1.159578 | 0.000000 | 0.000000 | 0.000000 |
| 57149 | *LYRM1* | 1.156830 | 0.000000 | 0.000000 | 0.000000 |
| 11247 | *NXPH4* | 1.141658 | 0.000000 | 0.000000 | 0.000000 |
| 1742 | *DLG4* | 1.140638 | 0.000000 | 0.000000 | 0.000000 |
| 2021 | *ENDOG* | 1.139483 | 0.000000 | 0.000000 | 0.000000 |
| 23523 | *CABIN1* | 1.138145 | 0.000000 | 0.000000 | 0.000000 |
| 823 | *CAPN1* | 1.136551 | 0.000000 | 0.000000 | 0.000000 |
| 23507 | *LRRC8B* | 1.136357 | 0.000000 | 0.000000 | 0.000000 |
| 79365 | *BHLHB3* | 1.135283 | 0.000000 | 0.000000 | 0.000000 |
| 5430 | *POLR2A* | 1.122144 | 0.000000 | 0.000000 | 0.000000 |
| 441204 | *LOC441204* | 1.121358 | 0.000000 | 0.000000 | 0.000000 |
| 8464 | *SUPT3H* | 1.115574 | 0.000000 | 0.000000 | 0.000000 |
| 10120 | *ACTR1B* | 1.111303 | 0.000000 | 0.000000 | 0.000000 |
| 54829 | *ASPN* | 1.108077 | 0.000000 | 0.000000 | 0.000000 |
| 10868 | *USP20* | 1.098175 | 0.000000 | 0.000000 | 0.000000 |
| 7554 | *ZNF8* | 1.090132 | 0.000000 | 0.000000 | 0.000000 |
| 2184 | *FAH* | 1.089733 | 0.000000 | 0.000000 | 0.000000 |
| 8653 | *DDX3Y* | 1.085470 | 1.181542 | 18.019358 | 25.764812 |
| 393 | *ARHGAP4* | 1.085241 | 0.000000 | 0.000000 | 0.000000 |
| 1521 | *CTSW* | 1.084480 | 0.893608 | 0.000000 | 0.000000 |
| 79956 | *KIAA1815* | 1.083423 | 0.000000 | 0.000000 | 0.000000 |

| | | | | | |
|---|---|---|---|---|---|
| 969 | *CD69* | 1.081601 | 0.613379 | 0.000000 | 0.000000 |
| 80184 | *CEP290* | 1.077368 | 0.000000 | 0.000000 | 0.000000 |
| 23108 | *GARNL4* | 1.067118 | 0.000000 | 0.000000 | 0.000000 |
| 408 | *ARRB1* | 1.057896 | 0.000000 | 0.000000 | 0.000000 |
| 2697 | *GJA1* | 1.057735 | 0.000000 | 0.000000 | 0.000000 |
| 4858 | *NOVA2* | 1.054483 | 0.000000 | 0.000000 | 0.000000 |
| 2077 | *ERF* | 1.046597 | 0.000000 | 0.000000 | 0.000000 |
| 3806 | *KIR2DS1* | 1.037691 | 0.000000 | 0.000000 | 0.000000 |
| 10352 | *WARS2* | 1.026628 | 0.000000 | 0.000000 | 0.000000 |
| 81848 | *SPRY4* | 1.016971 | 0.000000 | 0.000000 | 0.000000 |
| 4281 | *MID1* | 1.009281 | 0.000000 | 0.000000 | 0.000000 |
| 10773 | *ZBTB6* | 1.004462 | 0.000000 | 0.000000 | 0.000000 |
| 202559 | *KHDRBS2* | 0.999485 | 0.000000 | 0.000000 | 0.000000 |
| 861 | *RUNX1* | 0.998429 | 0.000000 | 0.000000 | 0.000000 |
| 55105 | *GPATCH2* | 0.996972 | 0.000000 | 0.000000 | 0.000000 |
| 7710 | *ZNF154* | 0.996805 | 0.000000 | 0.000000 | 0.000000 |
| 11057 | *ABHD2* | 0.996116 | 0.000000 | 0.000000 | 0.000000 |
| 9240 | *PNMA1* | 0.983399 | 0.000000 | 0.000000 | 0.000000 |
| 619426 | *C8orf60* | 0.982264 | 0.000000 | 0.000000 | 0.000000 |
| 54842 | *FLJ20160* | 0.980116 | 0.000000 | 0.000000 | 0.000000 |
| 574 | *BAGE* | 0.979960 | 0.000000 | 0.000000 | 0.000000 |
| 274 | *BIN1* | 0.974521 | 0.000000 | 0.000000 | 0.000000 |
| 10550 | *ARL6IP5* | 0.973653 | 0.000000 | 0.000000 | 0.000000 |
| 23329 | *KIAA0984* | 0.969202 | 0.000000 | 0.000000 | 0.000000 |
| 81873 | *ARPC5L* | 0.967062 | 0.411485 | 0.000000 | 0.000000 |
| 1875 | *E2F5* | 0.961138 | 0.000000 | 0.000000 | 0.000000 |
| 5287 | *PIK3C2B* | 0.956425 | 0.000000 | 0.000000 | 0.000000 |
| 1618 | *DAZL* | 0.951385 | 0.000000 | 0.000000 | 0.000000 |
| 8496 | *PPFIBP1* | 0.944134 | 0.000000 | 0.000000 | 0.000000 |
| 489 | *ATP2A3* | 0.943624 | 0.000000 | 0.000000 | 0.000000 |
| 9112 | *MTA1* | 0.934839 | 0.000000 | 0.000000 | 0.000000 |
| 4285 | *MIPEP* | 0.930868 | 0.000000 | 0.000000 | 0.000000 |
| 51701 | *NLK* | 0.927288 | 0.000000 | 0.000000 | 0.000000 |
| 56833 | *SLAMF8* | 0.914337 | 0.000000 | 0.000000 | 0.000000 |
| 80746 | *TSEN2* | 0.914297 | 0.000000 | 0.000000 | 0.000000 |
| 4669 | *NAGLU* | 0.910938 | 0.000000 | 0.000000 | 0.000000 |
| 29094 | *HSPC159* | 0.908306 | 0.000000 | 0.000000 | 0.000000 |
| 80129 | *C6orf97* | 0.905859 | 0.000000 | 0.000000 | 0.000000 |
| 120526 | *DPH4* | 0.903527 | 0.000000 | 0.000000 | 0.000000 |
| 2289 | *FKBP5* | 0.903161 | 0.000000 | 0.000000 | 0.000000 |
| 1804 | *DPP6* | 0.901471 | 0.000000 | 0.000000 | 0.000000 |
| 131544 | *DKFZp667G2110* | 0.891664 | 0.000000 | 0.000000 | 0.000000 |
| 79818 | *ZNF552* | 0.887879 | 0.000000 | 0.000000 | 0.000000 |
| 79000 | *C1orf135* | 0.880339 | 0.000000 | 0.000000 | 0.000000 |
| 27343 | *POLL* | 0.878064 | 0.000000 | 0.000000 | 0.000000 |
| 728411 | *LOC728411* | 0.868127 | 0.000000 | 0.000000 | 0.000000 |
| 5871 | *MAP4K2* | 0.868056 | 0.000000 | 0.000000 | 0.000000 |
| 10893 | *MMP24* | 0.867603 | 0.000000 | 0.000000 | 0.000000 |

| | | | | | |
|---|---|---|---|---|---|
| 4882 | *NPR2* | 0.866360 | 0.000000 | 0.000000 | 0.000000 |
| 55333 | *SYNJ2BP* | 0.865439 | 0.000000 | 0.000000 | 0.000000 |
| 400961 | *KIAA1155* | 0.864131 | 0.000000 | 0.000000 | 0.000000 |
| 80199 | *FUZ* | 0.862602 | 0.000000 | 0.000000 | 0.000000 |
| 10100 | *TSPAN2* | 0.858829 | 0.000000 | 0.000000 | 0.000000 |
| 51673 | *CGI-38* | 0.845245 | 0.000000 | 0.000000 | 0.000000 |
| 6477 | *SIAH1* | 0.840557 | 0.000000 | 0.000000 | 0.000000 |
| 161291 | *TMEM30B* | 0.838524 | 0.000000 | 0.000000 | 0.000000 |
| 55501 | *CHST12* | 0.836597 | 0.000000 | 0.000000 | 0.000000 |
| 6911 | *TBX6* | 0.831703 | 0.000000 | 0.000000 | 0.000000 |
| 1969 | *EPHA2* | 0.825048 | 0.000000 | 0.000000 | 0.000000 |
| 79830 | *ZMYM1* | 0.820074 | 0.000000 | 0.000000 | 0.000000 |
| 10911 | *UTS2* | 0.819225 | 0.000000 | 0.000000 | 0.000000 |
| 1373 | *CPS1* | 0.808854 | 0.000000 | 0.000000 | 0.000000 |
| 10620 | *ARID3B* | 0.800923 | 0.000000 | 0.000000 | 0.000000 |
| 8537 | *BCAS1* | 0.795566 | 0.000000 | 0.000000 | 0.000000 |
| 4040 | *LRP6* | 0.792976 | 0.000000 | 0.000000 | 0.000000 |
| 57634 | *EP400* | 0.786437 | 0.000000 | 0.000000 | 0.000000 |
| 23641 | *LDOC1* | 0.785847 | 0.000000 | 0.000000 | 0.000000 |
| 80095 | *ZNF606* | 0.782514 | 0.000000 | 0.000000 | 0.000000 |
| 307 | *ANXA4* | 0.781978 | 0.000000 | 0.000000 | 0.000000 |
| 51186 | *WBP5* | 0.780765 | 0.000000 | 0.000000 | 0.000000 |
| 11196 | *SEC23IP* | 0.780192 | 0.000000 | 0.000000 | 0.000000 |
| 26045 | *LRRTM2* | 0.779255 | 0.000000 | 0.000000 | 0.000000 |
| 9832 | *JAKMIP2* | 0.775551 | 0.000000 | 0.000000 | 0.000000 |
| 3912 | *LAMB1* | 0.769305 | 0.000000 | 0.000000 | 0.000000 |
| 9655 | *SOCS5* | 0.753611 | 0.000000 | 0.000000 | 0.000000 |
| 56953 | *NT5M* | 0.750062 | 0.000000 | 0.000000 | 0.000000 |
| 55359 | *STYK1* | 0.748668 | 0.000000 | 0.000000 | 0.000000 |
| 11072 | *DUSP14* | 0.743412 | 0.000000 | 0.000000 | 0.000000 |
| 3841 | *KPNA5* | 0.740589 | 0.000000 | 0.000000 | 0.000000 |
| 26052 | *DNM3* | 0.739923 | 0.000000 | 0.000000 | 0.000000 |
| 729164 | *LOC729164* | 0.735667 | 0.000000 | 0.000000 | 0.000000 |
| 4734 | *NEDD4* | 0.733173 | 0.000000 | 0.000000 | 0.000000 |
| 9421 | *HAND1* | 0.732023 | 0.000000 | 0.000000 | 0.000000 |
| 9086 | *EIF1AY* | 0.730849 | 4.919472 | 34.186362 | 47.835915 |
| 84617 | *TUBB6* | 0.719992 | 0.000000 | 0.000000 | 0.000000 |
| 5269 | *SERPINB6* | 0.719822 | 0.000000 | 0.000000 | 0.000000 |
| 157567 | *ANKRD46* | 0.718991 | 0.000000 | 0.000000 | 0.000000 |
| 11201 | *POLI* | 0.715387 | 0.000000 | 0.000000 | 0.000000 |
| 79783 | *C7orf10* | 0.712177 | 0.000000 | 0.000000 | 0.000000 |
| 7368 | *UGT8* | 0.703994 | 0.000000 | 0.000000 | 0.000000 |
| 7567 | *ZNF19* | 0.695740 | 0.000000 | 0.000000 | 0.000000 |
| 4218 | *RAB8A* | 0.694820 | 0.000000 | 0.000000 | 0.000000 |
| 2619 | *GAS1* | 0.684001 | 0.000000 | 0.000000 | 0.000000 |
| 2206 | *MS4A2* | 0.675589 | 0.000000 | 0.000000 | 0.000000 |
| 23285 | *KIAA1107* | 0.673970 | 0.000000 | 0.000000 | 0.000000 |
| 51626 | *DYNC2LI1* | 0.671814 | 0.000000 | 0.000000 | 0.000000 |

| | | | | | |
|---|---|---|---|---|---|
| 5205 | *ATP8B1* | 0.665265 | 0.000000 | 0.000000 | 0.000000 |
| 2261 | *FGFR3* | 0.658208 | 0.000000 | 0.000000 | 0.000000 |
| 9463 | *PICK1* | 0.657694 | 0.000000 | 0.000000 | 0.000000 |
| 23544 | *SEZ6L* | 0.637708 | 0.000000 | 0.000000 | 0.000000 |
| 1571 | *CYP2E1* | 0.632991 | 0.000000 | 0.000000 | 0.000000 |
| 23157 | *6-Sep* | 0.630249 | 0.000000 | 0.000000 | 0.000000 |
| 6558 | *SLC12A2* | 0.618326 | 0.000000 | 0.000000 | 0.000000 |
| 29965 | *C16orf5* | 0.613050 | 0.000000 | 0.000000 | 0.000000 |
| 389906 | *LOC389906* | 0.607102 | 0.000000 | 0.000000 | 0.000000 |
| 51393 | *TRPV2* | 0.604481 | 0.000000 | 0.000000 | 0.000000 |
| 10908 | *PNPLA6* | 0.599317 | 0.000000 | 0.000000 | 0.000000 |
| 3215 | *HOXB5* | 0.577378 | 0.000000 | 0.000000 | 0.000000 |
| 81698 | *C15orf5* | 0.576950 | 0.000000 | 0.000000 | 0.000000 |
| 10079 | *ATP9A* | 0.565877 | 0.000000 | 0.000000 | 0.000000 |
| 54874 | *FNBP1L* | 0.559609 | 0.000000 | 0.000000 | 0.000000 |
| 93594 | *WDR67* | 0.556477 | 0.000000 | 0.000000 | 0.000000 |
| 738 | *C11orf2* | 0.555517 | 0.000000 | 0.000000 | 0.000000 |
| 22859 | *LPHN1* | 0.552997 | 0.000000 | 0.000000 | 0.000000 |
| 7746 | *ZNF193* | 0.549297 | 0.000000 | 0.000000 | 0.000000 |
| 120227 | *CYP2R1* | 0.543984 | 0.000000 | 0.000000 | 0.000000 |
| 9215 | *LARGE* | 0.543739 | 0.000000 | 0.000000 | 0.000000 |
| 8099 | *CDK2AP1* | 0.543702 | 0.000000 | 0.000000 | 0.000000 |
| 55626 | *FLJ20294* | 0.541166 | 0.000000 | 0.000000 | 0.000000 |
| 7075 | *TIE1* | 0.538245 | 0.000000 | 0.000000 | 0.000000 |
| 79860 | *FLJ21369* | 0.519157 | 0.000000 | 0.000000 | 0.000000 |
| 4221 | *MEN1* | 0.514839 | 0.000000 | 0.000000 | 0.000000 |
| 820 | *CAMP* | 0.510996 | 0.000000 | 0.000000 | 0.000000 |
| 83988 | *NCALD* | 0.502300 | 0.000000 | 0.000000 | 0.000000 |
| 8420 | *SNHG3* | 0.498199 | 0.000000 | 0.000000 | 0.000000 |
| 1108 | *CHD4* | 0.494903 | 0.000000 | 0.000000 | 0.000000 |
| 11136 | *SLC7A9* | 0.487247 | 0.000000 | 0.000000 | 0.000000 |
| 192669 | *EIF2C3* | 0.478581 | 0.000000 | 0.000000 | 0.000000 |
| 26094 | *WDR21A* | 0.477995 | 0.000000 | 0.000000 | 0.000000 |
| 84914 | *ZNF587* | 0.475086 | 0.000000 | 0.000000 | 0.000000 |
| 3423 | *IDS* | 0.474274 | 0.000000 | 0.000000 | 0.000000 |
| 4778 | *NFE2* | 0.472232 | 0.000000 | 0.000000 | 0.000000 |
| 57669 | *EPB41L5* | 0.469381 | 0.000000 | 0.000000 | 0.000000 |
| 286343 | *C9orf150* | 0.467241 | 0.000000 | 0.000000 | 0.000000 |
| 55917 | *CTTNBP2NL* | 0.467218 | 0.000000 | 0.000000 | 0.000000 |
| 3615 | *IMPDH2* | 0.466308 | 0.000000 | 0.000000 | 0.000000 |
| 23363 | *OBSL1* | 0.462594 | 0.000000 | 0.000000 | 0.000000 |
| 55740 | *ENAH* | 0.458798 | 0.000000 | 0.000000 | 0.000000 |
| 9456 | *HOMER1* | 0.458509 | 0.000000 | 0.000000 | 0.000000 |
| 79646 | *PANK3* | 0.449312 | 0.000000 | 0.000000 | 0.000000 |
| 185 | *AGTR1* | 0.446305 | 0.000000 | 0.000000 | 0.000000 |
| 7042 | *TGFB2* | 0.432881 | 0.000000 | 0.000000 | 0.000000 |
| 6038 | *RNASE4* | 0.430534 | 0.000000 | 0.000000 | 0.000000 |
| 64864 | *RFXDC2* | 0.430301 | 0.000000 | 0.000000 | 0.000000 |

| | | | | | |
|---|---|---|---|---|---|
| 5581 | *PRKCE* | 0.417846 | 0.000000 | 0.000000 | 0.000000 |
| 84823 | *LMNB2* | 0.407471 | 0.000000 | 0.000000 | 0.000000 |
| 7029 | *TFDP2* | 0.405875 | 0.000000 | 0.000000 | 0.000000 |
| 8514 | *KCNAB2* | 0.401350 | 0.000000 | 0.000000 | 0.000000 |
| 54800 | *KLHL24* | 0.401067 | 0.000000 | 0.000000 | 0.000000 |
| 170626 | *PSMB1* | 0.399308 | 0.000000 | 0.000000 | 0.000000 |
| 7536 | *SF1* | 0.397583 | 0.000000 | 0.000000 | 0.000000 |
| 80055 | *PGAP1* | 0.395315 | 0.000000 | 0.000000 | 0.000000 |
| 396 | *ARHGDIA* | 0.395153 | 0.000000 | 0.000000 | 0.000000 |
| 9586 | *CREB5* | 0.394526 | 0.000000 | 0.000000 | 0.000000 |
| 51276 | *ZNF571* | 0.393981 | 0.000000 | 0.000000 | 0.000000 |
| 4782 | *NFIC* | 0.387729 | 0.000000 | 0.000000 | 0.000000 |
| 165 | *AEBP1* | 0.382559 | 0.000000 | 0.000000 | 0.000000 |
| 57707 | *KIAA1609* | 0.382018 | 0.000000 | 0.000000 | 0.000000 |
| 79724 | *ZNF768* | 0.376620 | 0.000000 | 0.000000 | 0.000000 |
| 623 | *BDKRB1* | 0.376217 | 0.000000 | 0.000000 | 0.000000 |
| 51195 | *RAPGEFL1* | 0.372735 | 0.000000 | 0.000000 | 0.000000 |
| 154661 | *RPIB9* | 0.364547 | 0.000000 | 0.000000 | 0.000000 |
| 10807 | *SDCCAG3* | 0.363353 | 0.000000 | 0.000000 | 0.000000 |
| 29760 | *BLNK* | 0.355190 | 0.000000 | 0.000000 | 0.000000 |
| 28981 | *IFT81* | 0.354988 | 0.000000 | 0.000000 | 0.000000 |
| 4287 | *ATXN3* | 0.341796 | 0.000000 | 0.000000 | 0.000000 |
| 51594 | *NAG* | 0.340113 | 0.000000 | 0.000000 | 0.000000 |
| 54014 | *BRWD1* | 0.340020 | 0.000000 | 0.000000 | 0.000000 |
| 7916 | *BAT2* | 0.339378 | 0.000000 | 0.000000 | 0.000000 |
| 2060 | *EPS15* | 0.338459 | 0.000000 | 0.000000 | 0.000000 |
| 6776 | *STAT5A* | 0.330994 | 0.000000 | 0.000000 | 0.000000 |
| 80263 | *TRIM45* | 0.321231 | 0.000000 | 0.000000 | 0.000000 |
| 23299 | *BICD2* | 0.320125 | 0.000000 | 0.000000 | 0.000000 |
| 9844 | *ELMO1* | 0.317100 | 0.000000 | 0.000000 | 0.000000 |
| 9760 | *TOX* | 0.311868 | 0.000000 | 0.000000 | 0.000000 |
| 29956 | *LASS2* | 0.308634 | 0.000000 | 0.000000 | 0.000000 |
| 57596 | *KIAA1446* | 0.302966 | 0.000000 | 0.000000 | 0.000000 |
| 1812 | *DRD1* | 0.301553 | 0.000000 | 0.000000 | 0.000000 |
| 29766 | *TMOD3* | 0.299153 | 0.000000 | 0.000000 | 0.000000 |
| 2664 | *GDI1* | 0.298769 | 0.000000 | 0.000000 | 0.000000 |
| 5257 | *PHKB* | 0.298091 | 0.000000 | 0.000000 | 0.000000 |
| 150726 | *FBXO41* | 0.284188 | 0.000000 | 0.000000 | 0.000000 |
| 3758 | *KCNJ1* | 0.284119 | 0.000000 | 0.000000 | 0.000000 |
| 80762 | *NDFIP1* | 0.276398 | 0.000000 | 0.000000 | 0.000000 |
| 80215 | *C21orf96* | 0.266367 | 0.000000 | 0.000000 | 0.000000 |
| 10842 | *C7orf16* | 0.258263 | 0.000000 | 0.000000 | 0.000000 |
| 10224 | *ZNF443* | 0.256692 | 0.000000 | 0.000000 | 0.000000 |
| 26015 | *RPAP1* | 0.253108 | 0.000000 | 0.000000 | 0.000000 |
| 4046 | *LSP1* | 0.248220 | 0.000000 | 0.000000 | 0.000000 |
| 5194 | *PEX13* | 0.239187 | 0.000000 | 0.000000 | 0.000000 |
| 4208 | *MEF2C* | 0.237710 | 0.000000 | 0.000000 | 0.000000 |
| 8302 | *KLRC4* | 0.232107 | 0.000000 | 0.000000 | 0.000000 |

| | | | | | |
|---|---|---|---|---|---|
| 11230 | *PRAF2* | 0.230619 | 0.000000 | 0.000000 | 0.000000 |
| 9410 | *WDR57* | 0.223586 | 0.000000 | 0.000000 | 0.000000 |
| 79848 | *CSPP1* | 0.222927 | 0.000000 | 0.000000 | 0.000000 |
| 79649 | *RP11-535K18.3* | 0.203996 | 0.000000 | 0.000000 | 0.000000 |
| 51561 | *IL23A* | 0.201782 | 0.000000 | 0.000000 | 0.000000 |
| 64223 | *GBL* | 0.197733 | 0.000000 | 0.000000 | 0.000000 |
| 390940 | *LOC390940* | 0.196630 | 0.000000 | 0.000000 | 0.000000 |
| 5898 | *RALA* | 0.190380 | 0.000000 | 0.000000 | 0.000000 |
| 201562 | *PTPLB* | 0.189994 | 0.000000 | 0.000000 | 0.000000 |
| 63897 | *ABC1* | 0.188853 | 0.000000 | 0.000000 | 0.000000 |
| 8706 | *B3GALNT1* | 0.179983 | 0.000000 | 0.000000 | 0.000000 |
| 8986 | *RPS6KA4* | 0.176393 | 0.000000 | 0.000000 | 0.000000 |
| 3249 | *HPN* | 0.169428 | 0.000000 | 0.000000 | 0.000000 |
| 9677 | *HISPPD2A* | 0.165114 | 0.000000 | 0.000000 | 0.000000 |
| 442240 | *ZNF259* | 0.161855 | 0.000000 | 0.000000 | 0.000000 |
| 11162 | *NUDT6* | 0.155794 | 0.000000 | 0.000000 | 0.000000 |
| 25788 | *RAD54B* | 0.152878 | 0.000000 | 0.000000 | 0.000000 |
| 10521 | *DDX17* | 0.135964 | 0.000000 | 0.000000 | 0.000000 |
| 55700 | *RPRC1* | 0.133322 | 0.000000 | 0.000000 | 0.000000 |
| 7639 | *ZNF85* | 0.132390 | 0.000000 | 0.000000 | 0.000000 |
| 54919 | *HEATR2* | 0.124359 | 0.000000 | 0.000000 | 0.000000 |
| 10336 | *PCGF3* | 0.123820 | 0.000000 | 0.000000 | 0.000000 |
| 9024 | *BRSK2* | 0.103164 | 0.000000 | 0.000000 | 0.000000 |
| 1296 | *COL8A2* | 0.101318 | 0.000000 | 0.000000 | 0.000000 |
| 10247 | *HRSP12* | 0.097055 | 0.000000 | 0.000000 | 0.000000 |
| 11186 | *RASSF1* | 0.092959 | 0.000000 | 0.000000 | 0.000000 |
| 64847 | *SPATA20* | 0.085086 | 0.000000 | 0.000000 | 0.000000 |
| 11272 | *PRR4* | 0.083523 | 0.000000 | 0.000000 | 0.000000 |
| 54020 | *SLC37A1* | 0.082077 | 0.000000 | 0.000000 | 0.000000 |
| 10940 | *POP1* | 0.080013 | 0.000000 | 0.000000 | 0.000000 |
| 3561 | *IL2RG* | 0.079096 | 0.000000 | 0.000000 | 0.000000 |
| 79921 | *TCEAL4* | 0.078029 | 0.000000 | 0.000000 | 0.000000 |
| 7041 | *TGFB1I1* | 0.076474 | 0.000000 | 0.000000 | 0.000000 |
| 60625 | *DHX35* | 0.075818 | 0.000000 | 0.000000 | 0.000000 |
| 81578 | *COL21A1* | 0.071200 | 0.000000 | 0.000000 | 0.000000 |
| 4057 | *LTF* | 0.065426 | 0.000000 | 0.000000 | 0.000000 |
| 51447 | *IHPK2* | 0.065092 | 0.000000 | 0.000000 | 0.000000 |
| 5607 | *MAP2K5* | 0.063751 | 0.000000 | 0.000000 | 0.000000 |
| 22870 | *SAPS1* | 0.058998 | 0.000000 | 0.000000 | 0.000000 |
| 643641 | *LOC643641* | 0.054239 | 0.000000 | 0.000000 | 0.000000 |
| 9891 | *NUAK1* | 0.051969 | 0.000000 | 0.000000 | 0.000000 |
| 55208 | *DCUN1D2* | 0.051614 | 0.000000 | 0.000000 | 0.000000 |
| 26037 | *SIPA1L1* | 0.048060 | 0.000000 | 0.000000 | 0.000000 |
| 63926 | *ANKRD5* | 0.040577 | 0.000000 | 0.000000 | 0.000000 |
| 290 | *ANPEP* | 0.039263 | 0.000000 | 0.000000 | 0.000000 |
| 254896 | *TNFRSF10C* | 0.038540 | 0.000000 | 0.000000 | 0.000000 |
| 6785 | *ELOVL4* | 0.036328 | 0.000000 | 0.000000 | 0.000000 |
| 8418 | *CMAH* | 0.035871 | 0.000000 | 0.000000 | 0.000000 |

| | | | | | |
|---|---|---|---|---|---|
| 6494 | *SIPA1* | 0.034864 | 0.000000 | 0.000000 | 0.000000 |
| 57658 | *CALCOCO1* | 0.033850 | 0.000000 | 0.000000 | 0.000000 |
| 3674 | *ITGA2B* | 0.032500 | 0.000000 | 0.000000 | 0.000000 |
| 3698 | *ITIH2* | 0.031420 | 0.000000 | 0.000000 | 0.000000 |
| 4086 | *SMAD1* | 0.029602 | 0.000000 | 0.000000 | 0.000000 |
| 9187 | *SLC24A1* | 0.029306 | 0.000000 | 0.000000 | 0.000000 |
| 1497 | *CTNS* | 0.025182 | 0.000000 | 0.000000 | 0.000000 |
| 51409 | *HEMK1* | 0.024152 | 0.000000 | 0.000000 | 0.000000 |
| 9877 | *ZC3H11A* | 0.023875 | 0.000000 | 0.000000 | 0.000000 |
| 5830 | *PEX5* | 0.018141 | 0.000000 | 0.000000 | 0.000000 |
| 22984 | *PDCD11* | 0.017613 | 0.000000 | 0.000000 | 0.000000 |
| 2527 | *FUT5* | 0.016784 | 0.000000 | 0.000000 | 0.000000 |
| 80742 | *PRR3* | 0.016362 | 0.000000 | 0.000000 | 0.000000 |
| 26275 | *HIBCH* | 0.015783 | 0.000000 | 0.000000 | 0.000000 |
| 55179 | *FAIM* | 0.015773 | 0.000000 | 0.000000 | 0.000000 |
| 4907 | *NT5E* | 0.011091 | 0.000000 | 0.000000 | 0.000000 |
| 65094 | *JMJD4* | 0.006146 | 0.000000 | 0.000000 | 0.000000 |
| 80224 | *NUBPL* | 0.001829 | 0.000000 | 0.000000 | 0.000000 |
| 60526 | *C2orf43* | 0.000835 | 0.000000 | 0.000000 | 0.000000 |
| 5683 | *PSMA2* | -0.000003 | 0.000000 | 0.000000 | 0.000000 |
| 27101 | *CACYBP* | -0.000952 | 0.000000 | 0.000000 | 0.000000 |
| 5937 | *RBMS1* | -0.008107 | 0.000000 | 0.000000 | 0.000000 |
| 339287 | *MSL-1* | -0.009395 | 0.000000 | 0.000000 | 0.000000 |
| 22833 | *KIAA0894* | -0.018683 | 0.000000 | 0.000000 | 0.000000 |
| 22908 | *SACM1L* | -0.020801 | 0.000000 | 0.000000 | 0.000000 |
| 8087 | *FXR1* | -0.023472 | 0.000000 | 0.000000 | 0.000000 |
| 5934 | *RBL2* | -0.027806 | 0.000000 | 0.000000 | 0.000000 |
| 2287 | *FKBP3* | -0.028846 | 0.000000 | 0.000000 | 0.000000 |
| 29086 | *HSPC142* | -0.032591 | 0.000000 | 0.000000 | 0.000000 |
| 11222 | *MRPL3* | -0.033122 | 0.000000 | 0.000000 | 0.000000 |
| 59338 | *PLEKHA1* | -0.033374 | 0.000000 | 0.000000 | 0.000000 |
| 10559 | *SLC35A1* | -0.035315 | 0.000000 | 0.000000 | 0.000000 |
| 406991 | *TMEM49* | -0.037999 | 0.000000 | 0.000000 | 0.000000 |
| 5437 | *POLR2H* | -0.041093 | 0.000000 | 0.000000 | 0.000000 |
| 7272 | *TTK* | -0.046437 | 0.000000 | 0.000000 | 0.000000 |
| 57235 | *KIAA0485* | -0.056029 | 0.000000 | 0.000000 | 0.000000 |
| 10933 | *MORF4L1* | -0.057403 | 0.000000 | 0.000000 | 0.000000 |
| 3507 | *IGHM* | -0.063909 | 0.000000 | 0.000000 | 0.000000 |
| 10170 | *DHRS9* | -0.065613 | 0.000000 | 0.000000 | 0.000000 |
| 4171 | *MCM2* | -0.068310 | 0.000000 | 0.000000 | 0.000000 |
| 57001 | *ACN9* | -0.072032 | 0.000000 | 0.000000 | 0.000000 |
| 6880 | *TAF9* | -0.072713 | 0.000000 | 0.000000 | 0.000000 |
| 5209 | *PFKFB3* | -0.075562 | 0.000000 | 0.000000 | 0.000000 |
| 5724 | *PTAFR* | -0.079309 | 0.000000 | 0.000000 | 0.000000 |
| 5236 | *PGM1* | -0.086457 | 0.000000 | 0.000000 | 0.000000 |
| 5176 | *SERPINF1* | -0.093572 | 0.000000 | 0.000000 | 0.000000 |
| 683 | *BST1* | -0.094302 | 0.000000 | 0.000000 | 0.000000 |
| 79145 | *CHCHD7* | -0.097435 | 0.000000 | 0.000000 | 0.000000 |

| | | | | | |
|---|---|---|---|---|---|
| 9435 | *CHST2* | -0.097552 | 0.000000 | 0.000000 | 0.000000 |
| 8535 | *CBX4* | -0.098570 | 0.000000 | 0.000000 | 0.000000 |
| 476 | *ATP1A1* | -0.098888 | 0.000000 | 0.000000 | 0.000000 |
| 92856 | *IMP4* | -0.104565 | 0.000000 | 0.000000 | 0.000000 |
| 7295 | *TXN* | -0.110185 | 0.000000 | 0.000000 | 0.000000 |
| 5476 | *CTSA* | -0.110451 | 0.000000 | 0.000000 | 0.000000 |
| 9592 | *IER2* | -0.114203 | 0.000000 | 0.000000 | 0.000000 |
| 10981 | *RAB32* | -0.114887 | 0.000000 | 0.000000 | 0.000000 |
| 55625 | *ZDHHC7* | -0.120394 | 0.000000 | 0.000000 | 0.000000 |
| 4809 | *NHP2L1* | -0.126783 | 0.000000 | 0.000000 | 0.000000 |
| 9562 | *MINPP1* | -0.130496 | 0.000000 | 0.000000 | 0.000000 |
| 55916 | *NXT2* | -0.141194 | 0.000000 | 0.000000 | 0.000000 |
| 55190 | *NUDT11* | -0.143005 | 0.000000 | 0.000000 | 0.000000 |
| 4292 | *MLH1* | -0.148426 | 0.000000 | 0.000000 | 0.000000 |
| 6446 | *SGK* | -0.149327 | 0.000000 | 0.000000 | 0.000000 |
| 55819 | *RNF130* | -0.153564 | 0.000000 | 0.000000 | 0.000000 |
| 2993 | *GYPA* | -0.157374 | 0.000000 | 0.000000 | 0.000000 |
| 6993 | *DYNLT1* | -0.158557 | 0.000000 | 0.000000 | 0.000000 |
| 23136 | *EPB41L3* | -0.161964 | 0.000000 | 0.000000 | 0.000000 |
| 3113 | *HLA-DPA1* | -0.164555 | 0.000000 | 0.000000 | 0.000000 |
| 1033 | *CDKN3* | -0.164840 | 0.000000 | 0.000000 | 0.000000 |
| 5695 | *PSMB7* | -0.168140 | 0.000000 | 0.000000 | 0.000000 |
| 10602 | *CDC42EP3* | -0.174721 | 0.000000 | 0.000000 | 0.000000 |
| 1545 | *CYP1B1* | -0.177745 | 0.000000 | 0.000000 | 0.000000 |
| 11157 | *LSM6* | -0.179506 | 0.000000 | 0.000000 | 0.000000 |
| 22978 | *NT5C2* | -0.185123 | 0.000000 | 0.000000 | 0.000000 |
| 963 | *CD53* | -0.187112 | 0.000000 | 0.000000 | 0.000000 |
| 9184 | *BUB3* | -0.191499 | 0.000000 | 0.000000 | 0.000000 |
| 55577 | *NAGK* | -0.207652 | 0.000000 | 0.000000 | 0.000000 |
| 6283 | *S100A12* | -0.207683 | 0.000000 | 0.000000 | 0.000000 |
| 27257 | *LSM1* | -0.209628 | 0.000000 | 0.000000 | 0.000000 |
| 7443 | *VRK1* | -0.210937 | 0.000000 | 0.000000 | 0.000000 |
| 114882 | *OSBPL8* | -0.215661 | 0.000000 | 0.000000 | 0.000000 |
| 57018 | *CCNL1* | -0.218230 | 0.000000 | 0.000000 | 0.000000 |
| 4717 | *NDUFC1* | -0.219672 | 0.000000 | 0.000000 | 0.000000 |
| 51596 | *CUTA* | -0.220240 | 0.000000 | 0.000000 | 0.000000 |
| 334 | *APLP2* | -0.222581 | 0.000000 | 0.000000 | 0.000000 |
| 9334 | *B4GALT5* | -0.225888 | 0.000000 | 0.000000 | 0.000000 |
| 3735 | *KARS* | -0.226730 | 0.000000 | 0.000000 | 0.000000 |
| 1130 | *LYST* | -0.230831 | 0.000000 | 0.000000 | 0.000000 |
| 9551 | *ATP5J2* | -0.231497 | 0.000000 | 0.000000 | 0.000000 |
| 5230 | *PGK1* | -0.238560 | 0.000000 | 0.000000 | 0.000000 |
| 8826 | *IQGAP1* | -0.238751 | 0.000000 | 0.000000 | 0.000000 |
| 64116 | *SLC39A8* | -0.241045 | 0.000000 | 0.000000 | 0.000000 |
| 51567 | *TTRAP* | -0.242813 | 0.000000 | 0.000000 | 0.000000 |
| 221960 | *C7orf28A* | -0.255914 | 0.000000 | 0.000000 | 0.000000 |
| 80273 | *GRPEL1* | -0.257055 | 0.000000 | 0.000000 | 0.000000 |
| 6629 | *SNRPB2* | -0.259340 | 0.000000 | 0.000000 | 0.000000 |

| | | | | | |
|---|---|---|---|---|---|
| 2672 | *GFI1* | -0.259744 | 0.000000 | 0.000000 | 0.000000 |
| 4125 | *MAN2B1* | -0.260256 | 0.000000 | 0.000000 | 0.000000 |
| 6631 | *SNRPC* | -0.261500 | 0.000000 | 0.000000 | 0.000000 |
| 5094 | *PCBP2* | -0.265932 | 0.000000 | 0.000000 | 0.000000 |
| 93974 | *ATPIF1* | -0.283520 | 0.000000 | 0.000000 | 0.000000 |
| 51593 | *ARS2* | -0.290435 | 0.000000 | 0.000000 | 0.000000 |
| 50809 | *HP1BP3* | -0.293994 | 0.000000 | 0.000000 | 0.000000 |
| 10327 | *AKR1A1* | -0.305007 | 0.000000 | 0.000000 | 0.000000 |
| 1649 | *DDIT3* | -0.305547 | 0.000000 | 0.000000 | 0.000000 |
| 518 | *ATP5G3* | -0.305667 | 0.000000 | 0.000000 | 0.000000 |
| 83440 | *ADPGK* | -0.314721 | 0.000000 | 0.000000 | 0.000000 |
| 4711 | *NDUFB5* | -0.315069 | 0.000000 | 0.000000 | 0.000000 |
| 8533 | *COPS3* | -0.325954 | 0.000000 | 0.000000 | 0.000000 |
| 2908 | *NR3C1* | -0.335953 | 0.000000 | 0.000000 | 0.000000 |
| 2791 | *GNG11* | -0.339936 | 0.000000 | 0.000000 | 0.000000 |
| 4282 | *MIF* | -0.341489 | 0.000000 | 0.000000 | 0.000000 |
| 55207 | *ARL8B* | -0.344230 | 0.000000 | 0.000000 | 0.000000 |
| 26519 | *TIMM10* | -0.348456 | 0.000000 | 0.000000 | 0.000000 |
| 22998 | *DKFZP686A0124* | -0.348505 | 0.000000 | 0.000000 | 0.000000 |
| 404636 | *FAM45B* | -0.352058 | 0.000000 | 0.000000 | 0.000000 |
| 90410 | *IFT20* | -0.352744 | 0.000000 | 0.000000 | 0.000000 |
| 51024 | *FIS1* | -0.356505 | 0.000000 | 0.000000 | 0.000000 |
| 3920 | *LAMP2* | -0.358961 | 0.000000 | 0.000000 | 0.000000 |
| 10160 | *FARP1* | -0.364585 | 0.000000 | 0.000000 | 0.000000 |
| 59342 | *SCPEP1* | -0.369902 | 0.000000 | 0.000000 | 0.000000 |
| 11191 | *PTENP1* | -0.370642 | 0.000000 | 0.000000 | 0.000000 |
| 4063 | *LY9* | -0.375642 | 0.000000 | 0.000000 | 0.000000 |
| 10389 | *SCML2* | -0.379062 | 0.000000 | 0.000000 | 0.000000 |
| 10672 | *GNA13* | -0.381399 | 0.000000 | 0.000000 | 0.000000 |
| 6646 | *SOAT1* | -0.382246 | 0.000000 | 0.000000 | 0.000000 |
| 11027 | *LILRA2* | -0.382905 | 0.000000 | 0.000000 | 0.000000 |
| 3512 | *IGJ* | -0.390183 | 0.000000 | 0.000000 | 0.000000 |
| 9775 | *EIF4A3* | -0.393348 | 0.000000 | 0.000000 | 0.000000 |
| 79801 | *SHCBP1* | -0.395195 | 0.000000 | 0.000000 | 0.000000 |
| 9352 | *TXNL1* | -0.398889 | 0.000000 | 0.000000 | 0.000000 |
| 7371 | *UCK2* | -0.405120 | 0.000000 | 0.000000 | 0.000000 |
| 5324 | *PLAG1* | -0.413193 | 0.000000 | 0.000000 | 0.000000 |
| 7555 | *CNBP* | -0.421344 | 0.000000 | 0.000000 | 0.000000 |
| 152100 | *MGC61571* | -0.428375 | 0.000000 | 0.000000 | 0.000000 |
| 529 | *ATP6V1E1* | -0.428439 | 0.000000 | 0.000000 | 0.000000 |
| 28974 | *C19orf53* | -0.429520 | 0.000000 | 0.000000 | 0.000000 |
| 8721 | *EDF1* | -0.431214 | 0.000000 | 0.000000 | 0.000000 |
| 54930 | *C14orf94* | -0.431770 | 0.000000 | 0.000000 | 0.000000 |
| 10402 | *ST3GAL6* | -0.437512 | 0.000000 | 0.000000 | 0.000000 |
| 81688 | *C6orf62* | -0.445741 | 0.000000 | 0.000000 | 0.000000 |
| 6708 | *SPTA1* | -0.451958 | 0.000000 | 0.000000 | 0.000000 |
| 6745 | *SSR1* | -0.466405 | 0.000000 | 0.000000 | 0.000000 |
| 23593 | *HEBP2* | -0.470192 | 0.000000 | 0.000000 | 0.000000 |

| | | | | | |
|---|---|---|---|---|---|
| 51309 | *ARMCX1* | -0.473686 | 0.000000 | 0.000000 | 0.000000 |
| 23370 | *ARHGEF18* | -0.485058 | 0.000000 | 0.000000 | 0.000000 |
| 347902 | *AMIGO2* | -0.491980 | 0.000000 | 0.000000 | 0.000000 |
| 3925 | *STMN1* | -0.500335 | 0.000000 | 0.000000 | 0.000000 |
| 917 | *CD3G* | -0.512986 | 0.000000 | 0.000000 | 0.000000 |
| 4082 | *MARCKS* | -0.520029 | 0.000000 | 0.000000 | 0.000000 |
| 8031 | *NCOA4* | -0.523594 | 0.000000 | 0.000000 | 0.000000 |
| 5110 | *PCMT1* | -0.528253 | 0.000000 | 0.000000 | 0.000000 |
| 51696 | *HECA* | -0.528535 | 0.000000 | 0.000000 | 0.000000 |
| 81614 | *NIPA2* | -0.530683 | 0.000000 | 0.000000 | 0.000000 |
| 29074 | *MRPL18* | -0.540473 | 0.000000 | 0.000000 | 0.000000 |
| 3182 | *HNRPAB* | -0.549991 | 0.000000 | 0.000000 | 0.000000 |
| 10661 | *KLF1* | -0.553146 | 0.000000 | 0.000000 | 0.000000 |
| 994 | *CDC25B* | -0.573259 | 0.000000 | 0.000000 | 0.000000 |
| 11340 | *EXOSC8* | -0.578548 | 0.000000 | 0.000000 | 0.000000 |
| 80145 | *THOC7* | -0.579262 | 0.000000 | 0.000000 | 0.000000 |
| 604 | *BCL6* | -0.579600 | 0.000000 | 0.000000 | 0.000000 |
| 7529 | *YWHAB* | -0.581988 | 0.000000 | 0.000000 | 0.000000 |
| 4635 | *MYL4* | -0.583989 | 0.000000 | 0.000000 | 0.000000 |
| 51601 | *LIPT1* | -0.591343 | 0.000000 | 0.000000 | 0.000000 |
| 200734 | *SPRED2* | -0.594153 | 0.000000 | 0.000000 | 0.000000 |
| 11328 | *FKBP9* | -0.595344 | 0.000000 | 0.000000 | 0.000000 |
| 4092 | *SMAD7* | -0.598992 | 0.000000 | 0.000000 | 0.000000 |
| 10095 | *ARPC1B* | -0.606094 | 0.000000 | 0.000000 | 0.000000 |
| 6622 | *SNCA* | -0.613310 | 0.000000 | 0.000000 | 0.000000 |
| 9377 | *COX5A* | -0.618615 | 0.000000 | 0.000000 | 0.000000 |
| 3398 | *ID2* | -0.621248 | 0.000000 | 0.000000 | 0.000000 |
| 10541 | *ANP32B* | -0.623124 | 0.000000 | 0.000000 | 0.000000 |
| 10023 | *FRAT1* | -0.624715 | 0.000000 | 0.000000 | 0.000000 |
| 3858 | *KRT10* | -0.626758 | 0.000000 | 0.000000 | 0.000000 |
| 6635 | *SNRPE* | -0.641773 | 0.000000 | 0.000000 | 0.000000 |
| 27440 | *CECR5* | -0.643373 | 0.000000 | 0.000000 | 0.000000 |
| 6583 | *SLC22A4* | -0.646499 | 0.000000 | 0.000000 | 0.000000 |
| 3903 | *LAIR1* | -0.648372 | 0.000000 | 0.000000 | 0.000000 |
| 991 | *CDC20* | -0.650442 | 0.000000 | 0.000000 | 0.000000 |
| 28831 | *IGLJ3* | -0.650969 | 0.000000 | 0.000000 | 0.000000 |
| 7514 | *XPO1* | -0.673133 | 0.000000 | 0.000000 | 0.000000 |
| 5688 | *PSMA7* | -0.693174 | 0.000000 | 0.000000 | 0.000000 |
| 51371 | *POMP* | -0.694314 | 0.000000 | 0.000000 | 0.000000 |
| 55619 | *DOCK10* | -0.695530 | 0.000000 | 0.000000 | 0.000000 |
| 6396 | *SEC13* | -0.697082 | 0.000000 | 0.000000 | 0.000000 |
| 65117 | *FLJ11021* | -0.701221 | 0.000000 | 0.000000 | 0.000000 |
| 10480 | *PCID1* | -0.702733 | 0.000000 | 0.000000 | 0.000000 |
| 9935 | *MAFB* | -0.711028 | 0.000000 | 0.000000 | 0.000000 |
| 7535 | *ZAP70* | -0.713241 | 0.000000 | 0.000000 | 0.000000 |
| 3937 | *LCP2* | -0.725562 | 0.000000 | 0.000000 | 0.000000 |
| 64061 | *TSPYL2* | -0.725656 | 0.000000 | 0.000000 | 0.000000 |
| 7378 | *UPP1* | -0.732942 | 0.000000 | 0.000000 | 0.000000 |

| | | | | | |
|---|---|---|---|---|---|
| 81552 | *ECOP* | -0.747411 | 0.000000 | 0.000000 | 0.000000 |
| 51430 | *C1orf9* | -0.749749 | 0.000000 | 0.000000 | 0.000000 |
| 27069 | *GHITM* | -0.752784 | 0.000000 | 0.000000 | 0.000000 |
| 4493 | *MT1E* | -0.754781 | 0.000000 | 0.000000 | 0.000000 |
| 51504 | *HSPC152* | -0.756432 | 0.000000 | 0.000000 | 0.000000 |
| 7913 | *DEK* | -0.767007 | 0.000000 | 0.000000 | 0.000000 |
| 5879 | *RAC1* | -0.768939 | 0.000000 | 0.000000 | 0.000000 |
| 9804 | *TOMM20* | -0.777347 | 0.000000 | 0.000000 | 0.000000 |
| 154 | *ADRB2* | -0.784663 | 0.000000 | 0.000000 | 0.000000 |
| 1052 | *CEBPD* | -0.786872 | 0.000000 | 0.000000 | 0.000000 |
| 3516 | *RBPSUH* | -0.787021 | 0.000000 | 0.000000 | 0.000000 |
| 527 | *ATP6V0C* | -0.792049 | 0.000000 | 0.000000 | 0.000000 |
| 3115 | *HLA-DPB1* | -0.799092 | 0.000000 | 0.000000 | 0.000000 |
| 25801 | *GCA* | -0.815084 | 0.000000 | 0.000000 | 0.000000 |
| 55596 | *ZCCHC8* | -0.816689 | 0.000000 | 0.000000 | 0.000000 |
| 64231 | *MS4A6A* | -0.827097 | 0.000000 | 0.000000 | 0.000000 |
| 9806 | *SPOCK2* | -0.833187 | 0.000000 | 0.000000 | 0.000000 |
| 4708 | *NDUFB2* | -0.834617 | 0.000000 | 0.000000 | 0.000000 |
| 4496 | *MT1H* | -0.840844 | 0.000000 | 0.000000 | 0.000000 |
| 328 | *APEX1* | -0.841496 | 0.000000 | 0.000000 | 0.000000 |
| 51278 | *IER5* | -0.843601 | 0.000000 | 0.000000 | 0.000000 |
| 4060 | *LUM* | -0.857643 | -1.272871 | -0.800792 | 0.000000 |
| 6277 | *S100A6* | -0.860518 | 0.000000 | 0.000000 | 0.000000 |
| 6596 | *HLTF* | -0.860548 | 0.000000 | 0.000000 | 0.000000 |
| 51176 | *LEF1* | -0.864049 | -0.381568 | 0.000000 | 0.000000 |
| 51622 | *C7orf28A* | -0.875326 | 0.000000 | 0.000000 | 0.000000 |
| 4709 | *NDUFB3* | -0.877128 | 0.000000 | 0.000000 | 0.000000 |
| 11260 | *XPOT* | -0.884556 | 0.000000 | 0.000000 | 0.000000 |
| 6632 | *SNRPD1* | -0.886100 | 0.000000 | 0.000000 | 0.000000 |
| 3720 | *JARID2* | -0.889696 | 0.000000 | 0.000000 | 0.000000 |
| 55651 | *NOLA2* | -0.892126 | 0.000000 | 0.000000 | 0.000000 |
| 55454 | *GALNACT-2* | -0.897174 | 0.000000 | 0.000000 | 0.000000 |
| 10092 | *ARPC5* | -0.899824 | 0.000000 | 0.000000 | 0.000000 |
| 8140 | *SLC7A5* | -0.913602 | 0.000000 | 0.000000 | 0.000000 |
| 2533 | *FYB* | -0.924508 | 0.000000 | 0.000000 | 0.000000 |
| 26234 | *FBXL5* | -0.926535 | 0.000000 | 0.000000 | 0.000000 |
| 79772 | *MCTP1* | -0.935385 | 0.000000 | 0.000000 | 0.000000 |
| 1983 | *EIF5* | -0.938273 | 0.000000 | 0.000000 | 0.000000 |
| 51429 | *SNX9* | -0.939763 | 0.000000 | 0.000000 | 0.000000 |
| 509 | *ATP5C1* | -0.943435 | 0.000000 | 0.000000 | 0.000000 |
| 4300 | *MLLT3* | -0.952193 | 0.000000 | 0.000000 | 0.000000 |
| 3183 | *HNRPC* | -0.953932 | 0.000000 | 0.000000 | 0.000000 |
| 3184 | *HNRPD* | -0.958656 | 0.000000 | 0.000000 | 0.000000 |
| 23312 | *DMXL2* | -0.959344 | 0.000000 | 0.000000 | 0.000000 |
| 5293 | *PIK3CD* | -0.960543 | 0.000000 | 0.000000 | 0.000000 |
| 79023 | *NUP37* | -0.961781 | 0.000000 | 0.000000 | 0.000000 |
| 51691 | *LSM8* | -0.969561 | 0.000000 | 0.000000 | 0.000000 |
| 824 | *CAPN2* | -0.969798 | 0.000000 | 0.000000 | 0.000000 |

| | | | | | |
|---|---|---|---|---|---|
| 6383 | *SDC2* | -0.973792 | 0.000000 | 0.000000 | 0.000000 |
| 55799 | *CACNA2D3* | -0.979391 | 0.000000 | 0.000000 | 0.000000 |
| 5436 | *POLR2G* | -0.986477 | 0.000000 | 0.000000 | 0.000000 |
| 811 | *CALR* | -0.987585 | 0.000000 | 0.000000 | 0.000000 |
| 115207 | *KCTD12* | -0.988375 | 0.000000 | 0.000000 | 0.000000 |
| 7167 | *TPI1* | -0.989121 | 0.000000 | 0.000000 | 0.000000 |
| 5196 | *PF4* | -0.990667 | 0.000000 | 0.000000 | 0.000000 |
| 9987 | *HNRPDL* | -1.025766 | 0.000000 | 0.000000 | 0.000000 |
| 7150 | *TOP1* | -1.029270 | 0.000000 | 0.000000 | 0.000000 |
| 5557 | *PRIM1* | -1.031372 | 0.000000 | 0.000000 | 0.000000 |
| 6430 | *SFRS5* | -1.060838 | 0.000000 | 0.000000 | 0.000000 |
| 7353 | *UFD1L* | -1.062673 | 0.000000 | 0.000000 | 0.000000 |
| 10175 | *CNIH* | -1.070887 | 0.000000 | 0.000000 | 0.000000 |
| 7323 | *UBE2D3* | -1.088258 | 0.000000 | 0.000000 | 0.000000 |
| 2113 | *ETS1* | -1.088698 | 0.000000 | 0.000000 | 0.000000 |
| 8661 | *EIF3S10* | -1.091211 | 0.000000 | 0.000000 | 0.000000 |
| 79005 | *SCNM1* | -1.092553 | 0.000000 | 0.000000 | 0.000000 |
| 6434 | *SFRS10* | -1.095761 | 0.000000 | 0.000000 | 0.000000 |
| 682 | *BSG* | -1.101360 | 0.000000 | 0.000000 | 0.000000 |
| 1039 | *CDR2* | -1.103145 | 0.000000 | 0.000000 | 0.000000 |
| 51094 | *ADIPOR1* | -1.108078 | 0.000000 | 0.000000 | 0.000000 |
| 64092 | *SAMSN1* | -1.113049 | 0.000000 | 0.000000 | 0.000000 |
| 10975 | *UQCR* | -1.113277 | 0.000000 | 0.000000 | 0.000000 |
| 81892 | *C14orf156* | -1.115346 | 0.000000 | 0.000000 | 0.000000 |
| 5143 | *PDE4C* | -1.116141 | 0.000000 | 0.000000 | 0.000000 |
| 3094 | *HINT1* | -1.123297 | 0.000000 | 0.000000 | 0.000000 |
| 22919 | *MAPRE1* | -1.123610 | 0.000000 | 0.000000 | 0.000000 |
| 10438 | *C1D* | -1.129834 | 0.000000 | 0.000000 | 0.000000 |
| 223 | *ALDH9A1* | -1.133718 | 0.000000 | 0.000000 | 0.000000 |
| 50807 | *DDEF1* | -1.135894 | 0.000000 | 0.000000 | 0.000000 |
| 26521 | *TIMM8B* | -1.138007 | 0.000000 | 0.000000 | 0.000000 |
| 6647 | *SOD1* | -1.146739 | 0.000000 | 0.000000 | 0.000000 |
| 3303 | *HSPA1A* | -1.147279 | 0.000000 | 0.000000 | 0.000000 |
| 7905 | *REEP5* | -1.150650 | 0.000000 | 0.000000 | 0.000000 |
| 4043 | *LRPAP1* | -1.161333 | 0.000000 | 0.000000 | 0.000000 |
| 4001 | *LMNB1* | -1.174198 | 0.000000 | 0.000000 | 0.000000 |
| 1200 | *TPP1* | -1.175675 | 0.000000 | 0.000000 | 0.000000 |
| 54407 | *SLC38A2* | -1.176939 | 0.000000 | 0.000000 | 0.000000 |
| 2091 | *FBL* | -1.178982 | 0.000000 | 0.000000 | 0.000000 |
| 11235 | *PDCD10* | -1.189315 | 0.000000 | 0.000000 | 0.000000 |
| 6636 | *SNRPF* | -1.191828 | 0.000000 | 0.000000 | 0.000000 |
| 27102 | *EIF2AK1* | -1.192437 | 0.000000 | 0.000000 | 0.000000 |
| 2879 | *GPX4* | -1.196254 | 0.000000 | 0.000000 | 0.000000 |
| 2098 | *ESD* | -1.198356 | 0.000000 | 0.000000 | 0.000000 |
| 5984 | *RFC4* | -1.201443 | 0.000000 | 0.000000 | 0.000000 |
| 226 | *ALDOA* | -1.205282 | 0.000000 | 0.000000 | 0.000000 |
| 55332 | *DRAM* | -1.208370 | 0.000000 | 0.000000 | 0.000000 |
| 60559 | *SPCS3* | -1.211818 | 0.000000 | 0.000000 | 0.000000 |

| | | | | | |
|---|---|---|---|---|---|
| 51065 | *RPS27L* | -1.231627 | 0.000000 | 0.000000 | 0.000000 |
| 517 | *ATP5G2* | -1.232104 | 0.000000 | 0.000000 | 0.000000 |
| 2534 | *FYN* | -1.232141 | 0.000000 | 0.000000 | 0.000000 |
| 56005 | *C19orf10* | -1.241227 | 0.000000 | 0.000000 | 0.000000 |
| 25793 | *FBXO7* | -1.244451 | 0.000000 | 0.000000 | 0.000000 |
| 78988 | *MRP63* | -1.245470 | 0.000000 | 0.000000 | 0.000000 |
| 10159 | *ATP6AP2* | -1.246465 | 0.000000 | 0.000000 | 0.000000 |
| 8727 | *CTNNAL1* | -1.246919 | 0.000000 | 0.000000 | 0.000000 |
| 4714 | *NDUFB8* | -1.247048 | 0.000000 | 0.000000 | 0.000000 |
| 6386 | *SDCBP* | -1.252275 | 0.000000 | 0.000000 | 0.000000 |
| 8665 | *EIF3S5* | -1.268982 | 0.000000 | 0.000000 | 0.000000 |
| 23753 | *SDF2L1* | -1.276541 | 0.000000 | 0.000000 | 0.000000 |
| 9997 | *SCO2* | -1.279216 | 0.000000 | 0.000000 | 0.000000 |
| 55771 | *PRR11* | -1.280899 | 0.000000 | 0.000000 | 0.000000 |
| 2841 | *GPR18* | -1.291652 | -0.390647 | 0.000000 | 0.000000 |
| 8848 | *TSC22D1* | -1.309616 | 0.000000 | 0.000000 | 0.000000 |
| 2647 | *BLOC1S1* | -1.316603 | 0.000000 | 0.000000 | 0.000000 |
| 80231 | *CXorf21* | -1.332875 | 0.000000 | 0.000000 | 0.000000 |
| 6275 | *S100A4* | -1.334947 | 0.000000 | 0.000000 | 0.000000 |
| 10140 | *TOB1* | -1.339014 | 0.000000 | 0.000000 | 0.000000 |
| 3301 | *DNAJA1* | -1.369961 | 0.000000 | 0.000000 | 0.000000 |
| 3162 | *HMOX1* | -1.374325 | 0.000000 | 0.000000 | 0.000000 |
| 1973 | *EIF4A1* | -1.377592 | 0.000000 | 0.000000 | 0.000000 |
| 8780 | *RIOK3* | -1.381180 | 0.000000 | 0.000000 | 0.000000 |
| 51669 | *TMEM66* | -1.386440 | 0.000000 | 0.000000 | 0.000000 |
| 50852 | *TRAT1* | -1.389158 | -1.018124 | -0.080597 | 0.000000 |
| 27258 | *LSM3* | -1.391751 | 0.000000 | 0.000000 | 0.000000 |
| 4696 | *NDUFA3* | -1.393046 | 0.000000 | 0.000000 | 0.000000 |
| 5730 | *PTGDS* | -1.407981 | 0.000000 | 0.000000 | 0.000000 |
| 22918 | *CD93* | -1.434421 | 0.000000 | 0.000000 | 0.000000 |
| 8804 | *CREG1* | -1.436174 | 0.000000 | 0.000000 | 0.000000 |
| 4700 | *NDUFA6* | -1.443726 | 0.000000 | 0.000000 | 0.000000 |
| 378 | *ARF4* | -1.464731 | 0.000000 | 0.000000 | 0.000000 |
| 4615 | *MYD88* | -1.464959 | 0.000000 | 0.000000 | 0.000000 |
| 3001 | *GZMA* | -1.468251 | 0.000000 | 0.000000 | 0.000000 |
| 4609 | *MYC* | -1.481369 | -0.282827 | 0.000000 | 0.000000 |
| 388677 | *NOTCH2NL* | -1.484569 | 0.000000 | 0.000000 | 0.000000 |
| 22934 | *RPIA* | -1.484857 | 0.000000 | 0.000000 | 0.000000 |
| 11258 | *DCTN3* | -1.491243 | 0.000000 | 0.000000 | 0.000000 |
| 3945 | *LDHB* | -1.495198 | 0.000000 | 0.000000 | 0.000000 |
| 51170 | *HSD17B11* | -1.518083 | 0.000000 | 0.000000 | 0.000000 |
| 1880 | *EBI2* | -1.523751 | 0.000000 | 0.000000 | 0.000000 |
| 6923 | *TCEB2* | -1.543355 | 0.000000 | 0.000000 | 0.000000 |
| 8668 | *EIF3S2* | -1.545996 | 0.000000 | 0.000000 | 0.000000 |
| 5861 | *RAB1A* | -1.547650 | 0.000000 | 0.000000 | 0.000000 |
| 972 | *CD74* | -1.551733 | 0.000000 | 0.000000 | 0.000000 |
| 8531 | *CSDA* | -1.552165 | 0.000000 | 0.000000 | 0.000000 |
| 3091 | *HIF1A* | -1.552777 | 0.000000 | 0.000000 | 0.000000 |

| | | | | | |
|---|---|---|---|---|---|
| 539 | *ATP5O* | -1.555199 | 0.000000 | 0.000000 | 0.000000 |
| 7037 | *TFRC* | -1.558606 | 0.000000 | 0.000000 | 0.000000 |
| 3059 | *HCLS1* | -1.559004 | 0.000000 | 0.000000 | 0.000000 |
| 10383 | *TUBB2C* | -1.569878 | 0.000000 | 0.000000 | 0.000000 |
| 5228 | *PGF* | -1.571303 | 0.000000 | 0.000000 | 0.000000 |
| 5111 | *PCNA* | -1.573690 | 0.000000 | 0.000000 | 0.000000 |
| 5721 | *PSME2* | -1.590199 | 0.000000 | 0.000000 | 0.000000 |
| 9936 | *CD302* | -1.594276 | 0.000000 | 0.000000 | 0.000000 |
| 55303 | *GIMAP4* | -1.598399 | 0.000000 | 0.000000 | 0.000000 |
| 6314 | *ATXN7* | -1.603227 | 0.000000 | 0.000000 | 0.000000 |
| 26034 | *PIP3-E* | -1.618231 | 0.000000 | 0.000000 | 0.000000 |
| 962 | *CD48* | -1.628415 | 0.000000 | 0.000000 | 0.000000 |
| 3560 | *IL2RB* | -1.639427 | 0.000000 | 0.000000 | 0.000000 |
| 3459 | *IFNGR1* | -1.645758 | 0.000000 | 0.000000 | 0.000000 |
| 1520 | *CTSS* | -1.650933 | 0.000000 | 0.000000 | 0.000000 |
| 23658 | *LSM5* | -1.704046 | 0.000000 | 0.000000 | 0.000000 |
| 2876 | *GPX1* | -1.705942 | 0.000000 | 0.000000 | 0.000000 |
| 2872 | *MKNK2* | -1.712153 | 0.000000 | 0.000000 | 0.000000 |
| 51121 | *RPL26L1* | -1.713980 | 0.000000 | 0.000000 | 0.000000 |
| 1431 | *CS* | -1.716725 | 0.000000 | 0.000000 | 0.000000 |
| 3133 | *HLA-E* | -1.721164 | 0.000000 | 0.000000 | 0.000000 |
| 4175 | *MCM6* | -1.725495 | 0.000000 | 0.000000 | 0.000000 |
| 7417 | *VDAC2* | -1.728283 | 0.000000 | 0.000000 | 0.000000 |
| 10289 | *EIF1B* | -1.745868 | 0.000000 | 0.000000 | 0.000000 |
| 6500 | *SKP1A* | -1.756645 | 0.000000 | 0.000000 | 0.000000 |
| 11177 | *BAZ1A* | -1.757487 | 0.000000 | 0.000000 | 0.000000 |
| 1175 | *AP2S1* | -1.761838 | 0.000000 | 0.000000 | 0.000000 |
| 54951 | *COMMD8* | -1.764045 | 0.000000 | 0.000000 | 0.000000 |
| 8992 | *ATP6V0E1* | -1.789308 | 0.000000 | 0.000000 | 0.000000 |
| 3002 | *GZMB* | -1.809057 | 0.000000 | 0.000000 | 0.000000 |
| 6351 | *CCL4* | -1.817220 | 0.000000 | 0.000000 | 0.000000 |
| 1337 | *COX6A1* | -1.825909 | 0.000000 | 0.000000 | 0.000000 |
| 8813 | *DPM1* | -1.833603 | 0.000000 | 0.000000 | 0.000000 |
| 51002 | *TPRKB* | -1.834480 | 0.000000 | 0.000000 | 0.000000 |
| 10972 | *TMED10* | -1.837042 | 0.000000 | 0.000000 | 0.000000 |
| 5901 | *RAN* | -1.857214 | 0.000000 | 0.000000 | 0.000000 |
| 4688 | *NCF2* | -1.860495 | 0.000000 | 0.000000 | 0.000000 |
| 2023 | *ENO1* | -1.866718 | -0.057435 | 0.000000 | 0.000000 |
| 51108 | *METTL9* | -1.872578 | 0.000000 | 0.000000 | 0.000000 |
| 1958 | *EGR1* | -1.889080 | 0.000000 | 0.000000 | 0.000000 |
| 29127 | *RACGAP1* | -1.889987 | 0.000000 | 0.000000 | 0.000000 |
| 55340 | *GIMAP5* | -1.897357 | 0.000000 | 0.000000 | 0.000000 |
| 3916 | *LAMP1* | -1.905594 | 0.000000 | 0.000000 | 0.000000 |
| 9847 | *KIAA0528* | -1.907690 | 0.000000 | 0.000000 | 0.000000 |
| 23480 | *SEC61G* | -1.930282 | 0.000000 | 0.000000 | 0.000000 |
| 79811 | *SLTM* | -1.943382 | 0.000000 | 0.000000 | 0.000000 |
| 9550 | *ATP6V1G1* | -1.950776 | 0.000000 | 0.000000 | 0.000000 |
| 56935 | *C11orf75* | -1.957374 | 0.000000 | 0.000000 | 0.000000 |

| | | | | | |
|---|---|---|---|---|---|
| 8843 | *GPR109B* | -1.959762 | 0.000000 | 0.000000 | 0.000000 |
| 9911 | *TMCC2* | -1.972353 | 0.000000 | 0.000000 | 0.000000 |
| 1072 | *CFL1* | -1.975071 | 0.000000 | 0.000000 | 0.000000 |
| 10169 | *SERF2* | -1.976055 | 0.000000 | 0.000000 | 0.000000 |
| 94239 | *H2AFV* | -1.986437 | 0.000000 | 0.000000 | 0.000000 |
| 10983 | *CCNI* | -1.995151 | 0.000000 | 0.000000 | 0.000000 |
| 2992 | *GYG1* | -2.003166 | 0.000000 | 0.000000 | 0.000000 |
| 51643 | *TMBIM4* | -2.021967 | 0.000000 | 0.000000 | 0.000000 |
| 427 | *ASAH1* | -2.024975 | 0.000000 | 0.000000 | 0.000000 |
| 29796 | *UCRC* | -2.043301 | 0.000000 | 0.000000 | 0.000000 |
| 3460 | *IFNGR2* | -2.055752 | 0.000000 | 0.000000 | 0.000000 |
| 126003 | *KIAA0256* | -2.056588 | 0.000000 | 0.000000 | 0.000000 |
| 6187 | *RPS2* | -2.059369 | 0.000000 | 0.000000 | 0.000000 |
| 7009 | *TEGT* | -2.062985 | 0.000000 | 0.000000 | 0.000000 |
| 4170 | *MCL1* | -2.064949 | 0.000000 | 0.000000 | 0.000000 |
| 3958 | *LGALS3* | -2.067700 | 0.000000 | 0.000000 | 0.000000 |
| 2923 | *PDIA3* | -2.075380 | 0.000000 | 0.000000 | 0.000000 |
| 4694 | *NDUFA1* | -2.077860 | 0.000000 | 0.000000 | 0.000000 |
| 10694 | *CCT8* | -2.094294 | 0.000000 | 0.000000 | 0.000000 |
| 5551 | *PRF1* | -2.094347 | 0.000000 | 0.000000 | 0.000000 |
| 9296 | *ATP6V1F* | -2.095502 | 0.000000 | 0.000000 | 0.000000 |
| 7097 | *TLR2* | -2.111238 | 0.000000 | 0.000000 | 0.000000 |
| 912 | *CD1D* | -2.118117 | -0.326607 | 0.000000 | 0.000000 |
| 8761 | *PABPC4* | -2.124236 | 0.000000 | 0.000000 | 0.000000 |
| 2203 | *FBP1* | -2.144255 | 0.000000 | 0.000000 | 0.000000 |
| 51650 | *MRPS33* | -2.163545 | 0.000000 | 0.000000 | 0.000000 |
| 4677 | *NARS* | -2.175795 | 0.000000 | 0.000000 | 0.000000 |
| 1535 | *CYBA* | -2.176093 | 0.000000 | 0.000000 | 0.000000 |
| 6427 | *SFRS2* | -2.187781 | 0.000000 | 0.000000 | 0.000000 |
| 59286 | *UBL5* | -2.194690 | 0.000000 | 0.000000 | 0.000000 |
| 10923 | *SUB1* | -2.204837 | 0.000000 | 0.000000 | 0.000000 |
| 9770 | *RASSF2* | -2.207409 | 0.000000 | 0.000000 | 0.000000 |
| 79682 | *MLF1IP* | -2.214578 | 0.000000 | 0.000000 | 0.000000 |
| 6612 | *SUMO3* | -2.222000 | 0.000000 | 0.000000 | 0.000000 |
| 6728 | *SRP19* | -2.238966 | 0.000000 | 0.000000 | 0.000000 |
| 79622 | *C16orf33* | -2.242247 | 0.000000 | 0.000000 | 0.000000 |
| 9055 | *PRC1* | -2.246947 | 0.000000 | 0.000000 | 0.000000 |
| 8743 | *TNFSF10* | -2.248009 | 0.000000 | 0.000000 | 0.000000 |
| 1665 | *DHX15* | -2.272001 | 0.000000 | 0.000000 | 0.000000 |
| 64757 | *MOSC1* | -2.272195 | 0.000000 | 0.000000 | 0.000000 |
| 79017 | *C7orf24* | -2.285002 | 0.000000 | 0.000000 | 0.000000 |
| 25984 | *KRT23* | -2.305236 | -0.317777 | 0.000000 | 0.000000 |
| 2717 | *GLA* | -2.307124 | 0.000000 | 0.000000 | 0.000000 |
| 9114 | *ATP6V0D1* | -2.310468 | 0.000000 | 0.000000 | 0.000000 |
| 3930 | *LBR* | -2.325176 | 0.000000 | 0.000000 | 0.000000 |
| 50486 | *G0S2* | -2.326231 | 0.000000 | 0.000000 | 0.000000 |
| 2219 | *FCN1* | -2.332227 | 0.000000 | 0.000000 | 0.000000 |
| 1315 | *COPB1* | -2.332419 | 0.000000 | 0.000000 | 0.000000 |

| | | | | | |
|---|---|---|---|---|---|
| 9315 | *C5orf13* | -2.333325 | -1.486585 | 0.000000 | 0.000000 |
| 7763 | *ZFAND5* | -2.339660 | 0.000000 | 0.000000 | 0.000000 |
| 10578 | *GNLY* | -2.352763 | 0.000000 | 0.000000 | 0.000000 |
| 23608 | *MKRN1* | -2.357541 | 0.000000 | 0.000000 | 0.000000 |
| 822 | *CAPG* | -2.362216 | 0.000000 | 0.000000 | 0.000000 |
| 5427 | *POLE2* | -2.370480 | 0.000000 | 0.000000 | 0.000000 |
| 6428 | *SFRS3* | -2.371885 | 0.000000 | 0.000000 | 0.000000 |
| 3682 | *ITGAE* | -2.380978 | 0.000000 | 0.000000 | 0.000000 |
| 387082 | *SUMO4* | -2.385770 | 0.000000 | 0.000000 | 0.000000 |
| 55013 | *CCDC109B* | -2.388287 | 0.000000 | 0.000000 | 0.000000 |
| 54107 | *POLE3* | -2.394571 | 0.000000 | 0.000000 | 0.000000 |
| 3838 | *KPNA2* | -2.397238 | 0.000000 | 0.000000 | 0.000000 |
| 50854 | *C6orf48* | -2.400012 | 0.000000 | 0.000000 | 0.000000 |
| 2752 | *GLUL* | -2.406938 | 0.000000 | 0.000000 | 0.000000 |
| 64919 | *TRA@* | -2.411374 | 0.000000 | 0.000000 | 0.000000 |
| 6091 | *ROBO1* | -2.425147 | -0.349784 | 0.000000 | 0.000000 |
| 6322 | *SCML1* | -2.441897 | -0.273806 | 0.000000 | 0.000000 |
| 51374 | *C2orf28* | -2.442867 | 0.000000 | 0.000000 | 0.000000 |
| 222161 | *DKFZp586I1420* | -2.467306 | 0.000000 | 0.000000 | 0.000000 |
| 10525 | *HYOU1* | -2.477327 | 0.000000 | 0.000000 | 0.000000 |
| 1655 | *DDX5* | -2.482575 | 0.000000 | 0.000000 | 0.000000 |
| 5789 | *PTPRD* | -2.491045 | -1.573493 | -0.011245 | 0.000000 |
| 4707 | *NDUFB1* | -2.507518 | 0.000000 | 0.000000 | 0.000000 |
| 805 | *CALM2* | -2.514088 | 0.000000 | 0.000000 | 0.000000 |
| 10875 | *FGL2* | -2.514126 | 0.000000 | 0.000000 | 0.000000 |
| 4494 | *MT1F* | -2.515389 | 0.000000 | 0.000000 | 0.000000 |
| 4085 | *MAD2L1* | -2.548710 | 0.000000 | 0.000000 | 0.000000 |
| 2217 | *FCGRT* | -2.552005 | 0.000000 | 0.000000 | 0.000000 |
| 313 | *AOAH* | -2.571060 | 0.000000 | 0.000000 | 0.000000 |
| 4601 | *MXI1* | -2.576819 | 0.000000 | 0.000000 | 0.000000 |
| 302 | *ANXA2* | -2.579344 | 0.000000 | 0.000000 | 0.000000 |
| 3021 | *H3F3B* | -2.581595 | 0.000000 | 0.000000 | 0.000000 |
| 56984 | *TNFSF5IP1* | -2.587984 | 0.000000 | 0.000000 | 0.000000 |
| 6727 | *SRP14* | -2.599303 | 0.000000 | 0.000000 | 0.000000 |
| 5720 | *PSME1* | -2.667895 | 0.000000 | 0.000000 | 0.000000 |
| 939 | *CD27* | -2.669589 | -0.867642 | 0.000000 | 0.000000 |
| 11345 | *GABARAPL2* | -2.672455 | 0.000000 | 0.000000 | 0.000000 |
| 8635 | *RNASET2* | -2.696252 | -0.537602 | 0.000000 | 0.000000 |
| 8908 | *GYG2* | -2.696356 | -2.455959 | -0.814591 | 0.000000 |
| 201725 | *TOMM7* | -2.699099 | 0.000000 | 0.000000 | 0.000000 |
| 10124 | *ARL4A* | -2.717765 | 0.000000 | 0.000000 | 0.000000 |
| 2235 | *FECH* | -2.718259 | 0.000000 | 0.000000 | 0.000000 |
| 1602 | *DACH1* | -2.721322 | -0.113022 | 0.000000 | 0.000000 |
| 7001 | *PRDX2* | -2.729110 | 0.000000 | 0.000000 | 0.000000 |
| 23479 | *ISCU* | -2.729637 | 0.000000 | 0.000000 | 0.000000 |
| 4118 | *MAL* | -2.752448 | -0.252069 | 0.000000 | 0.000000 |
| 11337 | *GABARAP* | -2.804327 | 0.000000 | 0.000000 | 0.000000 |
| 3149 | *HMGB3* | -2.807841 | -0.425883 | 0.000000 | 0.000000 |

| | | | | | |
|---|---|---|---|---|---|
| 6902 | *TBCA* | -2.815306 | 0.000000 | 0.000000 | 0.000000 |
| 4710 | *NDUFB4* | -2.821362 | 0.000000 | 0.000000 | 0.000000 |
| 1650 | *DDOST* | -2.824380 | 0.000000 | 0.000000 | 0.000000 |
| 84525 | *HOP* | -2.826259 | -0.756019 | 0.000000 | 0.000000 |
| 1786 | *DNMT1* | -2.827319 | 0.000000 | 0.000000 | 0.000000 |
| 5329 | *PLAUR* | -2.834112 | 0.000000 | 0.000000 | 0.000000 |
| 8277 | *TKTL1* | -2.838876 | 0.000000 | 0.000000 | 0.000000 |
| 5689 | *PSMB1* | -2.855248 | 0.000000 | 0.000000 | 0.000000 |
| 84790 | *TUBA6* | -2.861113 | 0.000000 | 0.000000 | 0.000000 |
| 1604 | *CD55* | -2.875967 | 0.000000 | 0.000000 | 0.000000 |
| 7940 | *LST1* | -2.920368 | 0.000000 | 0.000000 | 0.000000 |
| 1329 | *COX5B* | -2.946348 | 0.000000 | 0.000000 | 0.000000 |
| 4726 | *NDUFS6* | -2.951810 | 0.000000 | 0.000000 | 0.000000 |
| 10542 | *HBXIP* | -2.953858 | 0.000000 | 0.000000 | 0.000000 |
| 55969 | *C20orf24* | -2.962667 | 0.000000 | 0.000000 | 0.000000 |
| 7155 | *TOP2B* | -2.972653 | 0.000000 | 0.000000 | 0.000000 |
| 7419 | *VDAC3* | -2.981421 | 0.000000 | 0.000000 | 0.000000 |
| 400451 | *LOC400451* | -3.020655 | -1.536563 | 0.000000 | 0.000000 |
| 220988 | *HNRPA3* | -3.024955 | 0.000000 | 0.000000 | 0.000000 |
| 25907 | *TMEM158* | -3.030630 | -1.921186 | 0.000000 | 0.000000 |
| 6628 | *SNRPB* | -3.033574 | 0.000000 | 0.000000 | 0.000000 |
| 1163 | *CKS1B* | -3.035509 | 0.000000 | 0.000000 | 0.000000 |
| 468 | *ATF4* | -3.038890 | 0.000000 | 0.000000 | 0.000000 |
| 387 | *RHOA* | -3.064159 | 0.000000 | 0.000000 | 0.000000 |
| 51203 | *NUSAP1* | -3.074392 | 0.000000 | 0.000000 | 0.000000 |
| 7076 | *TIMP1* | -3.088943 | 0.000000 | 0.000000 | 0.000000 |
| 55505 | *NOLA3* | -3.109773 | 0.000000 | 0.000000 | 0.000000 |
| 304 | *ANXA2P2* | -3.116133 | 0.000000 | 0.000000 | 0.000000 |
| 7128 | *TNFAIP3* | -3.122360 | 0.000000 | 0.000000 | 0.000000 |
| 3535 | *SCGB2A2* | -3.174661 | -1.675545 | -0.187760 | 0.000000 |
| 7298 | *TYMS* | -3.189211 | 0.000000 | 0.000000 | 0.000000 |
| 821 | *CANX* | -3.204630 | 0.000000 | 0.000000 | 0.000000 |
| 64422 | *ATG3* | -3.228126 | 0.000000 | 0.000000 | 0.000000 |
| 3101 | *HK3* | -3.233719 | -0.360397 | 0.000000 | 0.000000 |
| 1508 | *CTSB* | -3.240003 | 0.000000 | 0.000000 | 0.000000 |
| 5250 | *SLC25A3* | -3.242213 | 0.000000 | 0.000000 | 0.000000 |
| 8836 | *GGH* | -3.242501 | 0.000000 | 0.000000 | 0.000000 |
| 54674 | *LRRN3* | -3.243912 | -1.182059 | 0.000000 | 0.000000 |
| 4154 | *MBNL1* | -3.305521 | -0.321151 | 0.000000 | 0.000000 |
| 51312 | *SLC25A37* | -3.319137 | -1.133385 | 0.000000 | 0.000000 |
| 1937 | *EEF1G* | -3.334259 | 0.000000 | 0.000000 | 0.000000 |
| 10096 | *ACTR3* | -3.345286 | 0.000000 | 0.000000 | 0.000000 |
| 1371 | *CPOX* | -3.377192 | 0.000000 | 0.000000 | 0.000000 |
| 54470 | *ARMCX6* | -3.378145 | 0.000000 | 0.000000 | 0.000000 |
| 1176 | *AP3S1* | -3.386323 | 0.000000 | 0.000000 | 0.000000 |
| 292 | *SLC25A5* | -3.389862 | 0.000000 | 0.000000 | 0.000000 |
| 2994 | *GYPB* | -3.402136 | -0.651392 | 0.000000 | 0.000000 |
| 2268 | *FGR* | -3.440166 | 0.000000 | 0.000000 | 0.000000 |

| | | | | | |
|---|---|---|---|---|---|
| 6185 | *RPN2* | -3.459849 | 0.000000 | 0.000000 | 0.000000 |
| 6119 | *RPA3* | -3.473089 | 0.000000 | 0.000000 | 0.000000 |
| 8655 | *DYNLL1* | -3.493778 | 0.000000 | 0.000000 | 0.000000 |
| 3181 | *HNRPA2B1* | -3.495778 | 0.000000 | 0.000000 | 0.000000 |
| 9802 | *DAZAP2* | -3.530402 | 0.000000 | 0.000000 | 0.000000 |
| 51097 | *SCCPDH* | -3.531832 | 0.000000 | 0.000000 | 0.000000 |
| 3956 | *LGALS1* | -3.541424 | 0.000000 | 0.000000 | 0.000000 |
| 1471 | *CST3* | -3.597429 | 0.000000 | 0.000000 | 0.000000 |
| 7538 | *ZFP36* | -3.605778 | 0.000000 | 0.000000 | 0.000000 |
| 10577 | *NPC2* | -3.618351 | 0.000000 | 0.000000 | 0.000000 |
| 2354 | *FOSB* | -3.639652 | 0.000000 | 0.000000 | 0.000000 |
| 3553 | *IL1B* | -3.639713 | 0.000000 | 0.000000 | 0.000000 |
| 9450 | *LY86* | -3.639992 | -0.813801 | 0.000000 | 0.000000 |
| 9978 | *RBX1* | -3.658831 | 0.000000 | 0.000000 | 0.000000 |
| 9214 | *FAIM3* | -3.672307 | -1.357744 | 0.000000 | 0.000000 |
| 5553 | *PRG2* | -3.673048 | 0.000000 | 0.000000 | 0.000000 |
| 5692 | *PSMB4* | -3.679756 | 0.000000 | 0.000000 | 0.000000 |
| 9920 | *KBTBD11* | -3.683285 | -1.198704 | 0.000000 | 0.000000 |
| 1441 | *CSF3R* | -3.687619 | 0.000000 | 0.000000 | 0.000000 |
| 1345 | *COX6C* | -3.687781 | 0.000000 | 0.000000 | 0.000000 |
| 10476 | *ATP5H* | -3.703596 | 0.000000 | 0.000000 | 0.000000 |
| 10628 | *TXNIP* | -3.730517 | -0.757724 | 0.000000 | 0.000000 |
| 51514 | *DTL* | -3.742015 | -0.211491 | 0.000000 | 0.000000 |
| 2778 | *GNAS* | -3.745816 | 0.000000 | 0.000000 | 0.000000 |
| 23787 | *MTCH1* | -3.746771 | 0.000000 | 0.000000 | 0.000000 |
| 27020 | *NPTN* | -3.840232 | 0.000000 | 0.000000 | 0.000000 |
| 1351 | *COX8A* | -3.855387 | 0.000000 | 0.000000 | 0.000000 |
| 10797 | *MTHFD2* | -3.858836 | 0.000000 | 0.000000 | 0.000000 |
| 22914 | *KLRK1* | -3.860056 | -1.237319 | 0.000000 | 0.000000 |
| 7277 | *TUBA1* | -3.869056 | 0.000000 | 0.000000 | 0.000000 |
| 3337 | *DNAJB1* | -3.876637 | 0.000000 | 0.000000 | 0.000000 |
| 28972 | *SPCS1* | -3.896429 | 0.000000 | 0.000000 | 0.000000 |
| 3587 | *IL10RA* | -3.905603 | 0.000000 | 0.000000 | 0.000000 |
| 29851 | *ICOS* | -3.921454 | -1.879515 | 0.000000 | 0.000000 |
| 916 | *CD3E* | -3.942904 | -1.439255 | 0.000000 | 0.000000 |
| 7386 | *UQCRFS1* | -3.960248 | 0.000000 | 0.000000 | 0.000000 |
| 498 | *ATP5A1* | -3.966577 | 0.000000 | 0.000000 | 0.000000 |
| 7873 | *ARMET* | -3.970728 | 0.000000 | 0.000000 | 0.000000 |
| 51187 | *C15orf15* | -3.982755 | 0.000000 | 0.000000 | 0.000000 |
| 6613 | *SUMO2* | -3.985738 | 0.000000 | 0.000000 | 0.000000 |
| 7357 | *UGCG* | -4.007079 | -1.117781 | 0.000000 | 0.000000 |
| 4501 | *MT1X* | -4.027102 | 0.000000 | 0.000000 | 0.000000 |
| 23204 | *ARL6IP1* | -4.042456 | 0.000000 | 0.000000 | 0.000000 |
| 919 | *CD247* | -4.072620 | -1.636376 | 0.000000 | 0.000000 |
| 8724 | *SNX3* | -4.088976 | 0.000000 | 0.000000 | 0.000000 |
| 7086 | *TKT* | -4.094839 | 0.000000 | 0.000000 | 0.000000 |
| 58526 | *MID1IP1* | -4.099627 | 0.000000 | 0.000000 | 0.000000 |
| 10397 | *NDRG1* | -4.105332 | 0.000000 | 0.000000 | 0.000000 |

| | | | | | |
|---|---|---|---|---|---|
| 9960 | *USP3* | -4.119186 | -0.294360 | 0.000000 | 0.000000 |
| 2038 | *EPB42* | -4.176079 | -0.538081 | 0.000000 | 0.000000 |
| 9446 | *GSTO1* | -4.214419 | 0.000000 | 0.000000 | 0.000000 |
| 199 | *AIF1* | -4.231407 | 0.000000 | 0.000000 | 0.000000 |
| 54539 | *NDUFB11* | -4.235702 | 0.000000 | 0.000000 | 0.000000 |
| 801 | *CALM1* | -4.246289 | -0.731078 | 0.000000 | 0.000000 |
| 8337 | *HIST2H2AA3* | -4.246380 | 0.000000 | 0.000000 | 0.000000 |
| 3187 | *HNRPH1* | -4.264599 | -0.158268 | 0.000000 | 0.000000 |
| 23765 | *IL17RA* | -4.266205 | -0.215856 | 0.000000 | 0.000000 |
| 1982 | *EIF4G2* | -4.318907 | -0.312587 | 0.000000 | 0.000000 |
| 6166 | *RPL36AL* | -4.352971 | 0.000000 | 0.000000 | 0.000000 |
| 10519 | *CIB1* | -4.375890 | 0.000000 | 0.000000 | 0.000000 |
| 103910 | *MRLC2* | -4.377004 | 0.000000 | 0.000000 | 0.000000 |
| 2180 | *ACSL1* | -4.444665 | -0.528089 | 0.000000 | 0.000000 |
| 399665 | *FAM102A* | -4.450466 | -3.075682 | 0.000000 | 0.000000 |
| 2745 | *GLRX* | -4.466248 | 0.000000 | 0.000000 | 0.000000 |
| 3003 | *GZMK* | -4.467223 | -2.155503 | 0.000000 | 0.000000 |
| 7381 | *UQCRB* | -4.468327 | -0.288234 | 0.000000 | 0.000000 |
| 7504 | *XK* | -4.470237 | -2.873600 | 0.000000 | 0.000000 |
| 533 | *ATP6V0B* | -4.488887 | 0.000000 | 0.000000 | 0.000000 |
| 8364 | *HIST1H4C* | -4.489980 | 0.000000 | 0.000000 | 0.000000 |
| 3936 | *LCP1* | -4.491365 | 0.000000 | 0.000000 | 0.000000 |
| 83443 | *SF3B5* | -4.505999 | 0.000000 | 0.000000 | 0.000000 |
| 2787 | *GNG5* | -4.549480 | 0.000000 | 0.000000 | 0.000000 |
| 51386 | *EIF3S6IP* | -4.569182 | 0.000000 | 0.000000 | 0.000000 |
| 6281 | *S100A10* | -4.584194 | -0.263034 | 0.000000 | 0.000000 |
| 5501 | *PPP1CC* | -4.615928 | 0.000000 | 0.000000 | 0.000000 |
| 9314 | *KLF4* | -4.616176 | -0.523872 | 0.000000 | 0.000000 |
| 9167 | *COX7A2L* | -4.629913 | 0.000000 | 0.000000 | 0.000000 |
| 2210 | *FCGR1B* | -4.655004 | -0.343708 | 0.000000 | 0.000000 |
| 3820 | *KLRB1* | -4.664435 | -1.927567 | 0.000000 | 0.000000 |
| 522 | *ATP5J* | -4.696250 | 0.000000 | 0.000000 | 0.000000 |
| 10135 | *PBEF1* | -4.743973 | -1.038474 | 0.000000 | 0.000000 |
| 10437 | *IFI30* | -4.746120 | -0.611160 | 0.000000 | 0.000000 |
| 301 | *ANXA1* | -4.751654 | -1.089112 | 0.000000 | 0.000000 |
| 1462 | *CSPG2* | -4.755344 | -2.728577 | 0.000000 | 0.000000 |
| 8091 | *HMGA2* | -4.757769 | -2.835858 | 0.000000 | 0.000000 |
| 6503 | *SLA* | -4.769731 | -0.768115 | 0.000000 | 0.000000 |
| 9882 | *TBC1D4* | -4.770290 | -3.175797 | -1.670430 | 0.000000 |
| 2971 | *GTF3A* | -4.780901 | 0.000000 | 0.000000 | 0.000000 |
| 51506 | *UFC1* | -4.793219 | 0.000000 | 0.000000 | 0.000000 |
| 6432 | *SFRS7* | -4.812831 | -0.628080 | 0.000000 | 0.000000 |
| 4830 | *NME1* | -4.837128 | 0.000000 | 0.000000 | 0.000000 |
| 3145 | *HMBS* | -4.841906 | 0.000000 | 0.000000 | 0.000000 |
| 10961 | *ERP29* | -4.846252 | 0.000000 | 0.000000 | 0.000000 |
| 925 | *CD8A* | -4.848936 | -2.431830 | 0.000000 | 0.000000 |
| 5687 | *PSMA6* | -4.869808 | 0.000000 | 0.000000 | 0.000000 |
| 5217 | *PFN2* | -4.887676 | -2.909191 | -1.060502 | 0.000000 |

| | | | | | |
|---|---|---|---|---|---|
| 2526 | *FUT4* | -4.922174 | -1.909268 | 0.000000 | 0.000000 |
| 27335 | *EIF3S12* | -4.923162 | 0.000000 | 0.000000 | 0.000000 |
| 51079 | *NDUFA13* | -4.957051 | 0.000000 | 0.000000 | 0.000000 |
| 11130 | *ZWINT* | -4.964823 | 0.000000 | 0.000000 | 0.000000 |
| 51218 | *GLRX5* | -4.993711 | 0.000000 | 0.000000 | 0.000000 |
| 8477 | *GPR65* | -5.013932 | -2.776514 | 0.000000 | 0.000000 |
| 2123 | *EVI2A* | -5.026945 | 0.000000 | 0.000000 | 0.000000 |
| 6282 | *S100A11* | -5.039967 | -0.043196 | 0.000000 | 0.000000 |
| 515 | *ATP5F1* | -5.096760 | 0.000000 | 0.000000 | 0.000000 |
| 25816 | *TNFAIP8* | -5.171258 | -2.296983 | 0.000000 | 0.000000 |
| 27242 | *TNFRSF21* | -5.201672 | -2.696264 | 0.000000 | 0.000000 |
| 5800 | *PTPRO* | -5.225824 | -1.793992 | 0.000000 | 0.000000 |
| 2069 | *EREG* | -5.267485 | -4.241950 | -0.479802 | 0.000000 |
| 8667 | *EIF3S3* | -5.278184 | 0.000000 | 0.000000 | 0.000000 |
| 11331 | *PHB2* | -5.312729 | 0.000000 | 0.000000 | 0.000000 |
| 5577 | *PRKAR2B* | -5.338753 | -2.249089 | 0.000000 | 0.000000 |
| 397 | *ARHGDIB* | -5.344016 | -0.268513 | 0.000000 | 0.000000 |
| 759 | *CA1* | -5.347738 | -2.911613 | 0.000000 | 0.000000 |
| 3490 | *IGFBP7* | -5.377769 | -0.967117 | 0.000000 | 0.000000 |
| 4023 | *LPL* | -5.412601 | -4.095376 | -1.968673 | 0.000000 |
| 3932 | *LCK* | -5.455262 | -4.069286 | 0.000000 | 0.000000 |
| 10412 | *TINP1* | -5.457591 | 0.000000 | 0.000000 | 0.000000 |
| 5691 | *PSMB3* | -5.484739 | 0.000000 | 0.000000 | 0.000000 |
| 6726 | *SRP9* | -5.485732 | 0.000000 | 0.000000 | 0.000000 |
| 706 | *TSPO* | -5.500875 | 0.000000 | 0.000000 | 0.000000 |
| 3394 | *IRF8* | -5.528965 | -2.587479 | 0.000000 | 0.000000 |
| 2171 | *FABP5* | -5.575313 | 0.000000 | 0.000000 | 0.000000 |
| 8409 | *UXT* | -5.581126 | 0.000000 | 0.000000 | 0.000000 |
| 10212 | *DDX39* | -5.584503 | 0.000000 | 0.000000 | 0.000000 |
| 4942 | *OAT* | -5.590434 | 0.000000 | 0.000000 | 0.000000 |
| 1843 | *DUSP1* | -5.672593 | -0.612111 | 0.000000 | 0.000000 |
| 22856 | *CHSY1* | -5.674875 | -0.616900 | 0.000000 | 0.000000 |
| 81611 | *ANP32E* | -5.691601 | -0.609927 | 0.000000 | 0.000000 |
| 3329 | *HSPD1* | -5.695750 | -2.049759 | 0.000000 | 0.000000 |
| 9555 | *H2AFY* | -5.719149 | -1.568495 | 0.000000 | 0.000000 |
| 29997 | *GLTSCR2* | -5.750382 | -0.948847 | 0.000000 | 0.000000 |
| 10094 | *ARPC3* | -5.759342 | 0.000000 | 0.000000 | 0.000000 |
| 6932 | *TCF7* | -5.789795 | -2.604480 | 0.000000 | 0.000000 |
| 8660 | *IRS2* | -5.824750 | -1.574624 | 0.000000 | 0.000000 |
| 4818 | *NKG7* | -5.839424 | -1.383600 | 0.000000 | 0.000000 |
| 4706 | *NDUFAB1* | -5.905269 | 0.000000 | 0.000000 | 0.000000 |
| 10627 | *MRCL3* | -5.938755 | 0.000000 | 0.000000 | 0.000000 |
| 1974 | *EIF4A2* | -6.041151 | -1.442696 | 0.000000 | 0.000000 |
| 5997 | *RGS2* | -6.081018 | -0.250703 | 0.000000 | 0.000000 |
| 28755 | *TRAC* | -6.083889 | -5.199883 | -1.213378 | 0.000000 |
| 6286 | *S100P* | -6.094713 | -3.839701 | -0.900819 | 0.000000 |
| 521 | *ATP5I* | -6.107640 | 0.000000 | 0.000000 | 0.000000 |
| 79630 | *C1orf54* | -6.132182 | -2.787389 | -0.075759 | 0.000000 |

| | | | | | |
|---|---|---|---|---|---|
| 10125 | *RASGRP1* | -6.148919 | -4.826082 | 0.000000 | 0.000000 |
| 23478 | *SEC11A* | -6.187897 | 0.000000 | 0.000000 | 0.000000 |
| 10109 | *ARPC2* | -6.212703 | 0.000000 | 0.000000 | 0.000000 |
| 6135 | *RPL11* | -6.256565 | 0.000000 | 0.000000 | 0.000000 |
| 1854 | *DUT* | -6.288733 | -0.254748 | 0.000000 | 0.000000 |
| 79668 | *PARP8* | -6.391602 | -1.913866 | 0.000000 | 0.000000 |
| 10613 | *SPFH1* | -6.426298 | -3.294605 | -0.190613 | 0.000000 |
| 1514 | *CTSL* | -6.525759 | -1.607367 | 0.000000 | 0.000000 |
| 914 | *CD2* | -6.562516 | -3.889621 | 0.000000 | 0.000000 |
| 6320 | *CLEC11A* | -6.584169 | -0.443805 | 0.000000 | 0.000000 |
| 3074 | *HEXB* | -6.643262 | -0.190075 | 0.000000 | 0.000000 |
| 10130 | *PDIA6* | -6.648757 | -0.572566 | 0.000000 | 0.000000 |
| 3135 | *HLA-G* | -6.671101 | -0.423837 | 0.000000 | 0.000000 |
| 27089 | *UQCRQ* | -6.689804 | 0.000000 | 0.000000 | 0.000000 |
| 10899 | *JTB* | -6.693234 | 0.000000 | 0.000000 | 0.000000 |
| 7305 | *TYROBP* | -6.731381 | 0.000000 | 0.000000 | 0.000000 |
| 1936 | *EEF1D* | -6.748821 | -1.716974 | 0.000000 | 0.000000 |
| 6167 | *RPL37* | -6.776287 | -1.129614 | 0.000000 | 0.000000 |
| 9535 | *GMFG* | -6.866004 | 0.000000 | 0.000000 | 0.000000 |
| 51053 | *GMNN* | -6.928895 | -1.054112 | 0.000000 | 0.000000 |
| 4904 | *YBX1* | -6.956631 | -1.698497 | 0.000000 | 0.000000 |
| 23406 | *COTL1* | -6.957187 | -1.281650 | 0.000000 | 0.000000 |
| 5538 | *PPT1* | -6.988325 | -0.144085 | 0.000000 | 0.000000 |
| 847 | *CAT* | -6.992064 | -2.785637 | 0.000000 | 0.000000 |
| 3150 | *HMGN1* | -7.002407 | 0.000000 | 0.000000 | 0.000000 |
| 6746 | *SSR2* | -7.184473 | 0.000000 | 0.000000 | 0.000000 |
| 6241 | *RRM2* | -7.194853 | -2.490224 | 0.000000 | 0.000000 |
| 3134 | *HLA-F* | -7.230095 | -2.188856 | 0.000000 | 0.000000 |
| 4860 | *NP* | -7.232026 | -1.331164 | 0.000000 | 0.000000 |
| 5358 | *PLS3* | -7.233104 | -5.117583 | -2.417202 | 0.000000 |
| 64747 | *MFSD1* | -7.303085 | 0.000000 | 0.000000 | 0.000000 |
| 2079 | *ERH* | -7.368767 | 0.000000 | 0.000000 | 0.000000 |
| 9235 | *IL32* | -7.505617 | -5.396581 | -0.676639 | 0.000000 |
| 144983 | *HNRPA1* | -7.569137 | -1.384468 | 0.000000 | 0.000000 |
| 2821 | *GPI* | -7.612209 | -1.828401 | 0.000000 | 0.000000 |
| 57570 | *TRMT5* | -7.669246 | -2.470649 | 0.000000 | 0.000000 |
| 1512 | *CTSH* | -7.680095 | -2.785040 | 0.000000 | 0.000000 |
| 6165 | *RPL35A* | -7.690887 | -2.001764 | 0.000000 | 0.000000 |
| 91353 | *CTA-246H3.1* | -7.762777 | -3.089102 | -0.001946 | 0.000000 |
| 4232 | *MEST* | -7.803542 | -4.928887 | -2.226133 | 0.000000 |
| 1327 | *COX4I1* | -7.838692 | -0.701564 | 0.000000 | 0.000000 |
| 4869 | *NPM1* | -7.841469 | 0.000000 | 0.000000 | 0.000000 |
| 11315 | *PARK7* | -7.845502 | 0.000000 | 0.000000 | 0.000000 |
| 9768 | *KIAA0101* | -7.878298 | -2.600625 | 0.000000 | 0.000000 |
| 689 | *BTF3* | -7.903117 | -2.744467 | 0.000000 | 0.000000 |
| 4953 | *ODC1* | -7.937681 | -0.711781 | 0.000000 | 0.000000 |
| 746 | *C11orf10* | -7.972265 | 0.000000 | 0.000000 | 0.000000 |
| 2495 | *FTH1* | -8.063371 | -0.671114 | 0.000000 | 0.000000 |

| | | | | | |
|---|---|---|---|---|---|
| 9556 | *C14orf2* | -8.088841 | -0.427805 | 0.000000 | 0.000000 |
| 6134 | *RPL10* | -8.145075 | -2.802903 | 0.000000 | 0.000000 |
| 7184 | *HSP90B1* | -8.150822 | -2.836618 | 0.000000 | 0.000000 |
| 6303 | *SAT1* | -8.184343 | -2.281829 | 0.000000 | 0.000000 |
| 5479 | *PPIB* | -8.194672 | -1.834800 | 0.000000 | 0.000000 |
| 7388 | *UQCRH* | -8.259499 | 0.000000 | 0.000000 | 0.000000 |
| 645 | *BLVRB* | -8.389441 | -1.067672 | 0.000000 | 0.000000 |
| 308 | *ANXA5* | -8.545533 | -2.772679 | 0.000000 | 0.000000 |
| 1347 | *COX7A2* | -8.579345 | -0.225680 | 0.000000 | 0.000000 |
| 441454 | *PTMA* | -8.602259 | 0.000000 | 0.000000 | 0.000000 |
| 55603 | *FAM46A* | -8.686905 | -5.537881 | -0.373180 | 0.000000 |
| 1066 | *CES1* | -8.765066 | -5.759753 | -1.509101 | 0.000000 |
| 10376 | *K-ALPHA-1* | -8.791910 | -1.503485 | 0.000000 | 0.000000 |
| 6210 | *RPS15A* | -8.836967 | -2.846474 | 0.000000 | 0.000000 |
| 4670 | *HNRPM* | -8.905446 | -2.689332 | 0.000000 | 0.000000 |
| 3122 | *HLA-DRA* | -9.046947 | -1.756368 | 0.000000 | 0.000000 |
| 6279 | *S100A8* | -9.074483 | -5.086002 | 0.000000 | 0.000000 |
| 60 | *ACTB* | -9.085821 | 0.000000 | 0.000000 | 0.000000 |
| 1350 | *COX7C* | -9.135334 | -1.113223 | 0.000000 | 0.000000 |
| 4753 | *NELL2* | -9.150172 | -7.670950 | -2.756024 | 0.000000 |
| 3146 | *HMGB1* | -9.168724 | -3.286960 | 0.000000 | 0.000000 |
| 1236 | *CCR7* | -9.219817 | -6.960453 | -0.432524 | 0.000000 |
| 9232 | *PTTG1* | -9.250713 | -2.603171 | 0.000000 | 0.000000 |
| 51142 | *CHCHD2* | -9.286454 | 0.000000 | 0.000000 | 0.000000 |
| 6156 | *RPL30* | -9.286574 | 0.000000 | 0.000000 | 0.000000 |
| 4613 | *MYCN* | -9.297111 | -6.582710 | -1.512191 | 0.000000 |
| 6039 | *RNASE6* | -9.324556 | -8.332271 | -3.984649 | 0.000000 |
| 3178 | *HNRPA1* | -9.340572 | -3.117126 | 0.000000 | 0.000000 |
| 7316 | *UBC* | -9.354553 | -1.101448 | 0.000000 | 0.000000 |
| 5204 | *PFDN5* | -9.401559 | 0.000000 | 0.000000 | 0.000000 |
| 6138 | *RPL15* | -9.404813 | -2.196310 | 0.000000 | 0.000000 |
| 2197 | *FAU* | -9.464537 | 0.000000 | 0.000000 | 0.000000 |
| 2207 | *FCER1G* | -9.470524 | -4.252769 | 0.000000 | 0.000000 |
| 24141 | *C20orf103* | -9.483314 | -7.490091 | -4.056357 | 0.000000 |
| 4697 | *NDUFA4* | -9.484173 | 0.000000 | 0.000000 | 0.000000 |
| 6146 | *RPL22* | -9.520043 | -4.177658 | 0.000000 | 0.000000 |
| 6203 | *RPS9* | -9.559504 | -3.634761 | 0.000000 | 0.000000 |
| 1340 | *COX6B1* | -9.563511 | 0.000000 | 0.000000 | 0.000000 |
| 7431 | *VIM* | -9.662770 | -4.170000 | 0.000000 | 0.000000 |
| 6637 | *SNRPG* | -9.772954 | -1.001352 | 0.000000 | 0.000000 |
| 6202 | *RPS8* | -9.822397 | -5.081495 | 0.000000 | 0.000000 |
| 760 | *CA2* | -9.822942 | -5.314717 | -0.828205 | 0.000000 |
| 202 | *AIM1* | -9.839355 | -5.040750 | -0.559519 | 0.000000 |
| 1051 | *CEBPB* | -9.873779 | -2.428561 | 0.000000 | 0.000000 |
| 10632 | *ATP5L* | -9.943671 | 0.000000 | 0.000000 | 0.000000 |
| 2509 | *FTHP1* | -9.950886 | -0.380386 | 0.000000 | 0.000000 |
| 3702 | *ITK* | -10.079002 | -8.645156 | -2.534076 | 0.000000 |
| 6181 | *RPLP2* | -10.115561 | -3.150508 | 0.000000 | 0.000000 |

| | | | | | |
|---|---|---|---|---|---|
| 6234 | *RPS28* | -10.129131 | -5.322690 | -0.022404 | 0.000000 |
| 514 | *ATP5E* | -10.233697 | 0.000000 | 0.000000 | 0.000000 |
| 6158 | *RPL28* | -10.307114 | -3.814474 | 0.000000 | 0.000000 |
| 2353 | *FOS* | -10.317573 | -2.253634 | 0.000000 | 0.000000 |
| 8673 | *VAMP8* | -10.432121 | -0.669456 | 0.000000 | 0.000000 |
| 9553 | *MRPL33* | -10.460546 | -2.779485 | 0.000000 | 0.000000 |
| 30846 | *EHD2* | -10.462750 | -7.472513 | -4.642830 | -2.872814 |
| 6352 | *CCL5* | -10.507117 | -4.619257 | -0.819479 | 0.000000 |
| 4946 | *OAZ1* | -10.576236 | -2.679115 | 0.000000 | 0.000000 |
| 3148 | *HMGB2* | -10.635271 | -4.822062 | -0.563263 | 0.000000 |
| 6125 | *RPL5* | -10.651211 | -4.155182 | 0.000000 | 0.000000 |
| 10209 | *EIF1* | -10.656947 | -5.613107 | -0.255408 | 0.000000 |
| 25873 | *RPL36* | -10.731355 | 0.000000 | 0.000000 | 0.000000 |
| 2896 | *GRN* | -10.770876 | -4.412922 | 0.000000 | 0.000000 |
| 915 | *CD3D* | -10.797929 | -6.930390 | -3.390074 | 0.000000 |
| 2124 | *EVI2B* | -10.825304 | -5.268876 | 0.000000 | 0.000000 |
| 6169 | *RPL38* | -10.827276 | -4.567137 | 0.000000 | 0.000000 |
| 6160 | *RPL31* | -10.829679 | -4.691652 | 0.000000 | 0.000000 |
| 967 | *CD63* | -10.852987 | -1.739577 | 0.000000 | 0.000000 |
| 6175 | *RPLP0* | -10.853851 | -0.115204 | 0.000000 | 0.000000 |
| 9045 | *RPL14* | -10.953021 | -3.287737 | 0.000000 | 0.000000 |
| 4925 | *NUCB2* | -10.961905 | -6.489966 | -2.572738 | 0.000000 |
| 3575 | *IL7R* | -10.983072 | -8.448246 | -2.056933 | 0.000000 |
| 26986 | *PABPC3* | -11.073365 | -6.533192 | -0.653548 | 0.000000 |
| 10075 | *HUWE1* | -11.104029 | -4.410072 | -0.746750 | 0.000000 |
| 9349 | *RPL23* | -11.125707 | -5.408854 | -0.616807 | 0.000000 |
| 6128 | *RPL6* | -11.174935 | -3.032958 | 0.000000 | 0.000000 |
| 5223 | *PGAM1* | -11.406106 | -3.861633 | 0.000000 | 0.000000 |
| 10399 | *GNB2L1* | -11.420632 | -5.416482 | -0.363508 | 0.000000 |
| 1829 | *DSG2* | -11.452505 | -10.229966 | -7.201186 | -2.431835 |
| 1164 | *CKS2* | -11.519116 | -4.211388 | 0.000000 | 0.000000 |
| 2597 | *GAPDH* | -11.669660 | -3.065763 | 0.000000 | 0.000000 |
| 51327 | *ERAF* | -11.713645 | -6.485393 | -1.444470 | 0.000000 |
| 4666 | *NACA* | -11.736722 | -2.447236 | 0.000000 | 0.000000 |
| 5660 | *PSAP* | -11.742829 | -4.155602 | 0.000000 | 0.000000 |
| 4602 | *MYB* | -11.767698 | -8.417980 | -3.416205 | -0.428223 |
| 6124 | *RPL4* | -11.834954 | 0.000000 | 0.000000 | 0.000000 |
| 5478 | *PPIA* | -11.859927 | -5.536818 | -0.783248 | 0.000000 |
| 4502 | *MT2A* | -11.911417 | -2.726813 | 0.000000 | 0.000000 |
| 3309 | *HSPA5* | -11.929098 | -5.820456 | -1.063182 | 0.000000 |
| 9709 | *HERPUD1* | -11.973175 | -5.481450 | -0.123120 | 0.000000 |
| 241 | *ALOX5AP* | -12.041402 | -7.534111 | -1.754784 | 0.000000 |
| 7311 | *UBA52* | -12.125992 | -3.296914 | 0.000000 | 0.000000 |
| 6748 | *SSR4* | -12.175992 | -3.656545 | 0.000000 | 0.000000 |
| 5757 | *PTMA* | -12.277341 | -1.531715 | 0.000000 | 0.000000 |
| 932 | *MS4A3* | -12.294124 | -10.458221 | -5.527925 | 0.000000 |
| 6132 | *RPL8* | -12.298303 | -0.078741 | 0.000000 | 0.000000 |
| 64852 | *TUT1* | -12.391937 | -7.494168 | -2.200654 | 0.000000 |

| | | | | | |
|---|---|---|---|---|---|
| 10370 | *CITED2* | -12.574052 | -6.467249 | -1.187449 | 0.000000 |
| 6231 | *RPS26* | -12.668389 | -0.403708 | 0.000000 | 0.000000 |
| 5873 | *RAB27A* | -12.808328 | -7.604228 | -2.874909 | 0.000000 |
| 6157 | *RPL27A* | -12.815162 | -5.873616 | -0.059168 | 0.000000 |
| 6228 | *RPS23* | -12.842386 | -7.862418 | -0.968028 | 0.000000 |
| 6173 | *RPL36A* | -12.876400 | -0.787356 | 0.000000 | 0.000000 |
| 6144 | *RPL21* | -12.966305 | -6.259079 | -0.162810 | 0.000000 |
| 3015 | *H2AFZ* | -13.059726 | -2.904019 | 0.000000 | 0.000000 |
| 3107 | *HLA-C* | -13.236621 | -7.669522 | -2.168232 | 0.000000 |
| 4352 | *MPL* | -13.306600 | -11.176914 | -7.091091 | -3.016208 |
| 5034 | *P4HB* | -13.334024 | -8.301861 | -2.722723 | 0.000000 |
| 6230 | *RPS25* | -13.488260 | -2.019585 | 0.000000 | 0.000000 |
| 6159 | *RPL29* | -13.511814 | -0.291015 | 0.000000 | 0.000000 |
| 4637 | *MYL6* | -13.525565 | -2.373155 | 0.000000 | 0.000000 |
| 4332 | *MNDA* | -13.619646 | -9.512613 | -2.847278 | 0.000000 |
| 6141 | *RPL18* | -13.766487 | -6.899957 | -0.825576 | 0.000000 |
| 6208 | *RPS14* | -13.800461 | -7.562223 | -0.783856 | 0.000000 |
| 51316 | *PLAC8* | -14.128131 | -5.965633 | 0.000000 | 0.000000 |
| 6129 | *RPL7* | -14.221095 | -7.323072 | -0.136481 | 0.000000 |
| 6130 | *RPL7A* | -14.578742 | -3.880537 | 0.000000 | 0.000000 |
| 6229 | *RPS24* | -14.603557 | -8.040045 | -0.710662 | 0.000000 |
| 10409 | *BASP1* | -15.280583 | -12.167716 | -7.321195 | -3.258970 |
| 6232 | *RPS27* | -15.383748 | -8.949784 | -2.484512 | 0.000000 |
| 6168 | *RPL37A* | -15.479430 | -8.385385 | -2.547111 | 0.000000 |
| 51816 | *CECR1* | -15.601857 | -9.709138 | -2.361217 | 0.000000 |
| 6223 | *RPS19* | -15.668252 | -8.215389 | -2.247944 | 0.000000 |
| 7314 | *UBB* | -15.675483 | -8.334118 | -2.159198 | 0.000000 |
| 57211 | *GPR126* | -15.741054 | -13.758150 | -11.042023 | -7.797849 |
| 1349 | *COX7B* | -15.822186 | -5.033441 | 0.000000 | 0.000000 |
| 1933 | *EEF1B2* | -16.089350 | -4.698821 | 0.000000 | 0.000000 |
| 5055 | *SERPINB2* | -16.154857 | -11.777563 | -3.537762 | 0.000000 |
| 6633 | *SNRPD2* | -16.236425 | -1.614238 | 0.000000 | 0.000000 |
| 7852 | *CXCR4* | -16.411939 | -12.976198 | -6.368529 | -0.711877 |
| 6227 | *RPS21* | -16.528616 | -5.794632 | -0.041904 | 0.000000 |
| 6233 | *RPS27A* | -16.675465 | -10.215265 | -2.803769 | 0.000000 |
| 6137 | *RPL13* | -16.724504 | -10.445005 | -4.297667 | -0.005309 |
| 6139 | *RPL17* | -17.003757 | -7.646118 | 0.000000 | 0.000000 |
| 11224 | *RPL35* | -17.050215 | -3.022827 | 0.000000 | 0.000000 |
| 6209 | *RPS15* | -17.114493 | -10.332511 | -3.655118 | -0.267045 |
| 6036 | *RNASE2* | -17.156764 | -11.949235 | -7.005911 | -0.512881 |
| 6176 | *RPLP1* | -17.403957 | -10.744868 | -5.271589 | -0.949695 |
| 4736 | *RPL10A* | -17.408054 | -9.260685 | -2.835765 | 0.000000 |
| 6191 | *RPS4X* | -17.420658 | -10.103028 | -4.404949 | -0.134653 |
| 694 | *BTG1* | -17.493455 | -12.909982 | -7.725790 | -1.958460 |
| 3151 | *HMGN2* | -17.559868 | -7.404462 | 0.000000 | 0.000000 |
| 51372 | *CCDC72* | -18.555720 | -6.261816 | 0.000000 | 0.000000 |
| 6280 | *S100A9* | -18.681380 | -8.366455 | -5.808097 | -1.275154 |
| 6143 | *RPL19* | -18.716587 | -6.187813 | 0.000000 | 0.000000 |

| | | | | | |
|---|---|---|---|---|---|
| 3921 | *RPSA* | -19.029806 | -6.943896 | 0.000000 | 0.000000 |
| 3106 | *HLA-B* | -19.306296 | -10.203045 | -2.210232 | 0.000000 |
| 55827 | *IQWD1* | -19.325902 | -10.590411 | -4.020718 | -0.761956 |
| 23521 | *RPL13A* | -19.465419 | -5.170585 | 0.000000 | 0.000000 |
| 6205 | *RPS11* | -19.817968 | -12.005081 | -4.769759 | -1.616462 |
| 6217 | *RPS16* | -21.005192 | -12.053854 | -4.761310 | 0.000000 |
| 8530 | *CST7* | -21.075012 | -15.085782 | -8.534416 | -2.213034 |
| 1475 | *CSTA* | -21.080991 | -12.492963 | -4.999834 | 0.000000 |
| 6152 | *RPL24* | -21.162243 | -6.937672 | 0.000000 | 0.000000 |
| 6194 | *RPS6* | -21.242554 | -13.354283 | -6.908761 | -0.699192 |
| 6206 | *RPS12* | -21.557374 | -4.753650 | 0.000000 | 0.000000 |
| 567 | *B2M* | -21.684923 | -13.994407 | -7.070947 | -2.484003 |
| 6201 | *RPS7* | -21.695832 | -8.005436 | 0.000000 | 0.000000 |
| 6136 | *RPL12* | -22.243141 | -7.686923 | 0.000000 | 0.000000 |
| 3105 | *HLA-A* | -22.494332 | -14.072490 | -7.212473 | -0.732498 |
| 1915 | *EEF1A1* | -22.894552 | -15.105348 | -7.819078 | -2.269781 |
| 6207 | *RPS13* | -23.158730 | -9.747921 | -0.307961 | 0.000000 |
| 6235 | *RPS29* | -23.751638 | -4.718924 | 0.000000 | 0.000000 |
| 6204 | *RPS10* | -25.160867 | -14.248573 | -6.072768 | 0.000000 |
| 1675 | *CFD* | -25.850743 | -18.159717 | -10.371001 | -2.754991 |
| 6218 | *RPS17* | -25.937502 | -13.062370 | 0.000000 | 0.000000 |
| 5552 | *PRG1* | -26.607427 | -19.883112 | -11.907298 | -4.341454 |
| 2512 | *FTL* | -27.042121 | -11.678799 | -2.401752 | 0.000000 |
| 6037 | *RNASE3* | -28.060542 | -24.501391 | -17.077363 | -9.639906 |
| 6161 | *RPL32* | -29.245661 | -15.483326 | -1.642399 | 0.000000 |
| 3045 | *HBD* | -30.848787 | -19.785336 | -11.430465 | -10.620286 |
| 6155 | *RPL27* | -31.072216 | -16.232074 | -5.260344 | 0.000000 |
| 9168 | *TMSB10* | -32.696140 | -16.793030 | -5.878550 | -0.265437 |
| 7178 | *TPT1* | -33.044862 | -16.171061 | -5.817343 | 0.000000 |
| 6189 | *RPS3A* | -33.796025 | -21.567861 | -8.129200 | -1.609656 |
| 1511 | *CTSG* | -33.821659 | -28.571328 | -22.926675 | -15.111896 |
| 6147 | *RPL23A* | -40.532033 | -29.020685 | -6.762963 | 0.000000 |
| 3043 | *HBB* | -40.704275 | -34.805679 | -25.589047 | -19.835775 |
| 6133 | *RPL9* | -43.812241 | -24.015539 | -9.258505 | 0.000000 |
| 6222 | *RPS18* | -45.801470 | -26.378625 | -8.907302 | 0.000000 |
| 6224 | *RPS20* | -46.676179 | -27.358997 | -10.259791 | -2.083969 |
| 4353 | *MPO* | -47.674276 | -37.205415 | -26.795725 | -15.388090 |
| 4069 | *LYZ* | -56.665828 | -50.772997 | -37.652102 | -27.727456 |
| 1991 | *ELA2* | -57.747273 | -51.209218 | -42.867886 | -35.774702 |
| 566 | *AZU1* | -60.087828 | -47.263396 | -36.021064 | -27.421245 |
| 6170 | *RPL39* | -61.904344 | -41.630507 | -27.663496 | -18.183008 |
| 6171 | *RPL41* | -67.709222 | -43.996932 | -22.909097 | -5.269906 |
