## Supplementary Methods for "Computational deconvolution of gene expression in leukemic cell hierarchies"

#### Deconvolution model

To detect cell type-specific changes in gene expression, we solved the optimization problem

$$\min_{W_1,W_2,H_1,H_2} ||A - W_1H_1^T - W_2H_2^T||_F^2 + \lambda||W_1 - \tilde{W}_1||_1, \text{ s.t. } H_1, H_2 \geq 0. \tag{1}$$

Here, $A$ is a $m \times n$ matrix containing $m$-dimensional gene expression profiles of $n$ unsorted samples from a given tumor type (thousands of samples). The $m \times k_1$ matrix $\tilde{W}_1$ contains reference gene expression profiles of $k_1$ normal cell types in the tissue-of-origin (these vectors can be estimated using samples from a limited number of healthy subjects). The $m \times k_1$ matrix $W_1$ and the $m \times k_2$ matrix $W_2$ represent unknown gene expression vectors. The $n \times k_1$ matrix $H_1$ and $n \times k_2$ matrix $H_2$ contain the weights of these in each sample in $A$.

The $\mathcal{L}^1$ norm in the second term plays an important role in our model. Firstly, this term imposes sparsity in the sense that $W_1$ and $\tilde{W}_1$ will differ only at a few elements. Secondly, it controls the distance between the estimated patterns in $W_1$ and reference patterns in $\tilde{W}_1$. When $\lambda \to 0$, the estimated patterns in $W_1$ are allowed to move farther from the reference patterns in $\tilde{W}_1$. This increases the freedom in the estimation, as the optimization will driven to a larger extent by statistical variation in $A$ and to a lesser extent by prior information. The results will approach those obtained with unsupervised decomposition methods[3,21,26,28]. Conversely, a large $\lambda$ will force $W_1$ towards $\tilde{W}_1$, while $W_1$ can still be estimated freely. In this case, our model approaches previously published deconvolution methods, which assume a fixed $W_1$ and only estimate $H_1$[1,4,7,9,11–13,19,20,23,24,30–32].

As for $W_2$, this matrix is estimated without proximity constraints. It serves to absorb additional variation, including variation in frequency and gene expression in additional cell types or batch effects (*i.e.*, technical differences between $A$ and $W_1$).

To calculate solutions to the optimization problem, we adopted a block-coordinate descent algorithm[26], where we alternatingly used an efficient non-negative least squares algorithm to update $H_1$ and $H_2$[2] and a robust rank-1 decomposition to update $W_1$ and $W_2$[26]. For robustness, we bootstrapped (resampled) the $A$ matrix 200 times, and used the element-wise median of $W_1$ in all downstream analyses.

#### Gene expression data sets

We used gene expression profiles of blood and bone marrow samples from a total of 2,799 patients with AML, previously generated in ten different studies[5,8,10,14,15,18,22,25,27,29], and were retrieved from the NCBI Gene Expression Omnibus (http://www.ncbi.nlm.nih.gov/gds; accession numbers GSE6891, GSE10358, GSE13159, GSE14468, GSE15434, GSE17855, GSE22845, GSE21261, GSE22056, GSE12417). Further, we used a control set of 1,554 gene expression of other hematologic malignancies extracted from the GSE13159 data set[8]. All data were generated on Affymetrix microarrays and quantile-normalized to a log-normal distribution.

#### Deconvolution experiments

We applied our deconvolution method to the pooled set of 2,799 AML gene expression profiles, as well as to the ten AML data sets separately and the control data for other hematologic malignancies. As reference gene expression vectors in the $\tilde{W}_1$ matrix, we used expression profiles of sorted blood cells from the Differentiation Map (DMAP; accession number GSE24759)[17]. While all analyses were carried out with a broad range of $\lambda$ values (0.1 to

1.0), we found empirically that $\lambda$ values in the lower range (order of 0.10 to 0.25) were needed to detect AML-associated signals for rare cell types like HSC1 (**Supplementary Figures 1** to **3**; **Supplementary Table 1**). The analyses shown were carried out with 2 unconstrained $W_2$ vectors. For completeness, the analyses were repeated with other reasonable numbers of unconstrained $W_2$ vectors (0 to 3), yielding results in broad agreement with those shown.

#### Gene expression data set for sorted AML cell fractions

To test the AML-LSC relevance of identified gene expression patterns, we used an external set of gene expression profiles of AML samples fractionated by fluorescence-activated cell sorting using the cell surface markers CD34 and CD38 into four fractions ($CD34^{+}38^{-}$, $CD34^{+}38^{+}$, $CD34^{-}38^{+}$ and $CD34^{-}38^{-}$ cells)[6]. From sorted cells, RNA was purified and analyzed on Affymetrix microarrays (NCBI Gene Expression Omnibus accession number GSE30375). In parallel, AML-LSC activity in each sorted fraction was assayed using a limited dilution assay based on *in vivo* xenotransplantation into NOD-scid gamma mice. The AML-LSC relevance of each gene was calculated by correlating AML-LSC activity with gene expression. Once these scores (correlation coefficients) had been calculated, we tested for enrichments of high AML-LSC relevance scores among genes that were up- or downregulated in AML cell types compared to normal blood cells. For enrichment testing, we used the program RenderCat[16]. Sets of up- and downregulated genes in different cell types in AML samples were defined by enumerating the genes with positive and negative values in each column of $W_1 - \tilde{W}_1$.

### References


1. Abbas, A. R., Wolslegel, K., Seshasayee, D., Modrusan, Z., and Clark, H. F. (2009). Deconvolution of blood microarray data identifies cellular activation patterns in systemic lupus erythematosus. *PLoS One*, **4**(7), e6098.

2. Benthem, M.H. Keenan, M. (2004). Fast algorithm for the solution of large-scale non-negativity-constrained least squares problems. *Journal of Chemometrics*, **18**, 441–450.

3. Brunet, J.-P., Tamayo, P., Golub, T. R., and Mesirov, J. P. (2004). Metagenes and molecular pattern discovery using matrix factorization. *Proc Natl Acad Sci U S A*, **101**(12), 4164–4169.

4. Clarke, J., Seo, P., and Clarke, B. (2010). Statistical expression deconvolution from mixed tissue samples. *Bioinformatics*, **26**(8), 1043–1049.

5. de Jonge, H. J. M., Valk, P. J. M., Veeger, N. J. G. M., ter Elst, A., den Boer, M. L., Cloos, J., de Haas, V., van den Heuvel-Eibrink, M. M., Kaspers, G. J. L., Zwaan, C. M., Kamps, W. A., Lowenberg, B., and de Bont, E. S. J. M. (2010). High vegfc expression is associated with unique gene expression profiles and predicts adverse prognosis in pediatric and adult acute myeloid leukemia. *Blood*, **116**(10), 1747–1754.

6. Eppert, K., Takenaka, K., Lechman, E. R., Waldron, L., Nilsson, B., van Galen, P., Metzeler, K. H., Poeppl, A., Ling, V., Beyene, J., Canty, A. J., Danska, J. S., Bohlander, S. K., Buske, C., Minden, M. D., Golub, T. R., Jurisica, I., Ebert, B. L., and Dick, J. E. (2011). Stem cell gene expression programs influence clinical outcome in human leukemia. *Nat Med*, **17**(9), 1086–1093.

7. Gong, T. and Szustakowski, J. D. (2013). Deconrnaseq: a statistical framework for deconvolution of heterogeneous tissue samples based on mrna-seq data. *Bioinformatics*, **29**(8), 1083–1085.

8. Haferlach, T. *et al.* (2010). Clinical utility of microarray-based gene expression profiling in the diagnosis and subclassification of leukemia: report from the international microarray innovations in leukemia study group. *J Clin Oncol*, **28**(15), 2529–2537.

9. Hoffman, E. (2004). Expression profiling–best practices for data generation and interpretation in clinical trials. *Nat Rev Genet*, **5**(3), 229–237.

10. Klein, H.-U., Ruckert, C., Kohlmann, A., Bullinger, L., Thiede, C., Haferlach, T., and Dugas, M. (2009). Quantitative comparison of microarray experiments with published leukemia related gene expression signatures. *BMC Bioinformatics*, **10**, 422.

11. Kuhn, A., Kumar, A., Beilina, A., Dillman, A., Cookson, M. R., and Singleton, A. B. (2012). Cell population-specific expression analysis of human cerebellum. *BMC Genomics*, **13**, 610.

12. Lahdesmaki, H., Shmulevich, L., Dunmire, V., Yli-Harja, O., and Zhang, W. (2005). In silico microdissection of microarray data from heterogeneous cell populations. *BMC Bioinformatics*, **6**, 54.

13. Lu, P., Nakorchevskiy, A., and Marcotte, E. M. (2003). Expression deconvolution: a reinterpretation of dna microarray data reveals dynamic changes in cell populations. *Proc Natl Acad Sci U S A*, **100**(18), 10370–10375.

14. Metzeler, K. H., Hummel, M., Bloomfield, C. D., Spiekermann, K., Braess, J., Sauerland, M.-C., Heinecke, A., Radmacher, M., Marcucci, G., Whitman, S. P., Maharry, K., Paschka, P., Larson, R. A., Berdel, W. E., Bochner, T., Wormann, B., Mansmann, U., Hiddemann, W., Bohlander, S. K., Buske, C., , C., B, L. G., and , G. A. M. L. C. G. (2008). An 86-probe-set gene-expression signature predicts survival in cytogenetically normal acute myeloid leukemia. *Blood*, **112**(10), 4193–4201.

15. Miesner, M., Haferlach, C., Bacher, U., Weiss, T., Macijewski, K., Kohlmann, A., Klein, H.-U., Dugas, M., Kern, W., Schnittger, S., and Haferlach, T. (2010). Multilineage dysplasia (MLD) in acute myeloid leukemia (AML) correlates with MDS-related cytogenetic abnormalities and a prior history of MDS or MDS/MPN but has no independent prognostic relevance: a comparison of 408 cases classified as "aml not otherwise specified" (AML-NOS) or "AML with myelodysplasia-related changes" (AML-MRC). *Blood*, **116**(15), 2742–2751.

16. Nilsson, L., Eden, P., Olsson, E., Mansson, R., Astrand-Grundstrom, I., Strombeck, B., Theilgaard-Monch, K., Anderson, K., Hast, R., Hellstrom-Lindberg, E., Samuelsson, J., Bergh, G., Nerlov, C., Johansson, B., Sigvardsson, M., Borg, A., and Jacobsen, S. E. W. (2007). The molecular signature of MDS stem cells supports a stem-cell origin of 5q myelodysplastic syndromes. *Blood*, **110**(8), 3005–3014.

17. Novershtern, N., Subramanian, A., Lawton, L. N., Mak, R. H., Haining, W. N., McConkey, M. E., Habib, N., Yosef, N., Chang, C. Y., Shay, T., Frampton, G. M., Drake, A. C. B., Leskov, I., Nilsson, B., Preffer, F., Dombkowski, D., Evans, J. W., Liefeld, T., Smutko, J. S., Chen, J., Friedman, N., Young, R. A., Golub, T. R., Regev, A., and Ebert, B. L. (2011). Densely interconnected transcriptional circuits control cell states in human hematopoiesis. *Cell*, **144**(2), 296–309.

18. Payton, J. E., Grieselhuber, N. R., Chang, L.-W., Murakami, M., Geiss, G. K., Link, D. C., Nagarajan, R., Watson, M. A., and Ley, T. J. (2009). High throughput digital quantification of mrna abundance in primary human acute myeloid leukemia samples. *J Clin Invest*, **119**(6), 1714–1726.

19. Qi, L., Li, B., Dong, Y., Xu, H., Chen, L., Wang, H., Li, P., Zhao, W., Gu, Y., Wang, C., and Guo, Z. (2014). Deconvolution of the gene expression profiles of valuable banked blood specimens for studying the prognostic values of altered peripheral immune cell proportions in cancer patients. *PLoS One*, **9**(6), e100934.

20. Qiao, W., Quon, G., Csaszar, E., Yu, M., Morris, Q., and Zandstra, P. W. (2012). PERT: a method for expression deconvolution of human blood samples from varied microenvironmental and developmental conditions. *PLoS Comput Biol*, **8**(12), e1002838.

21. Ringnér, M. (2008). What is principal component analysis? *Nat Biotechnol*, **26**(3), 303–304.

22. Sandahl, J. D., Coenen, E. A., Forestier, E., Harbott, J., Johansson, B., Kerndrup, G., Adachi, S., Auvrignon, A., Beverloo, H. B., Cayuela, J.-M., Chilton, L., Fornerod, M., de Haas, V., Harrison, C. J., Inaba, H., Kaspers, G. J. L., Liang, D.-C., Locatelli, F.,

Masetti, R., Perot, C., Raimondi, S. C., Reinhardt, K., Tomizawa, D., von Neuhoff, N., Zecca, M., Zwaan, C. M., van den Heuvel-Eibrink, M. M., and Hasle, H. (2014). t(6;9)(p22;q34)/DEK-NUP214-rearranged pediatric myeloid leukemia: an international study of 62 patients. *Haematologica*, **99**(5), 865–872.

23. Shannon, C. P., Balshaw, R., Ng, R. T., Wilson-McManus, J. E., Keown, P., McMaster, R., McManus, B. M., Landsberg, D., Isbel, N. M., Knoll, G., and Tebbutt, S. J. (2014). Two-stage, in silico deconvolution of the lymphocyte compartment of the peripheral whole blood transcriptome in the context of acute kidney allograft rejection. *PLoS One*, **9**(4), e95224.

24. Shen-Orr, S. S., Tibshirani, R., Khatri, P., Bodian, D. L., Staedtler, F., Perry, N. M., Hastie, T., Sarwal, M. M., Davis, M. M., and Butte, A. J. (2010). Cell type-specific gene expression differences in complex tissues. *Nat Methods*, **7**(4), 287–289.

25. Taskesen, E. *et al.* (2011). Prognostic impact, concurrent genetic mutations, and gene expression features of aml with cebpa mutations in a cohort of 1182 cytogenetically normal aml patients: further evidence for cebpa double mutant aml as a distinctive disease entity. *Blood*, **117**(8), 2469–2475.

26. Taslaman, L. and Nilsson, B. (2012). A framework for regularized non-negative matrix factorization, with application to the analysis of gene expression data. *PLoS One*, **7**(11), e46331.

27. Tomasson, M. H., Xiang, Z., Walgren, R., Zhao, Y., Kasai, Y., Miner, T., Ries, R. E., Lubman, O., Fremont, D. H., McLellan, M. D., Payton, J. E., Westervelt, P., DiPersio, J. F., Link, D. C., Walter, M. J., Graubert, T. A., Watson, M., Baty, J., Heath, S., Shannon, W. D., Nagarajan, R., Bloomfield, C. D., Mardis, E. R., Wilson, R. K., and Ley, T. J. (2008). Somatic mutations and germline sequence variants in the expressed tyrosine kinase genes of patients with de novo acute myeloid leukemia. *Blood*, **111**(9), 4797–4808.

28. Venet, D., Pecasse, F., Maenhaut, C., and Bersini, H. (2001). Separation of samples into their constituents using gene expression data. *Bioinformatics*, **17 Suppl 1**, S279–S287.

29. Verhaak, R. G. W., Wouters, B. J., Erpelinck, C. A. J., Abbas, S., Beverloo, H. B., Lugthart, S., Löwenberg, B., Delwel, R., and Valk, P. J. M. (2009). Prediction of molecular subtypes in acute myeloid leukemia based on gene expression profiling. *Haematologica*, **94**(1), 131–134.

30. Wang, M., Master, S. R., and Chodosh, L. A. (2006). Computational expression deconvolution in a complex mammalian organ. *BMC Bioinformatics*, **7**, 328.

31. Wang, N., Gong, T., Clarke, R., Chen, L., Shih, I.-M., Zhang, Z., Levine, D. A., Xuan, J., and Wang, Y. (2015). Undo: a bioconductor r package for unsupervised deconvolution of mixed gene expressions in tumor samples. *Bioinformatics*, **31**(1), 137–139.

32. Zhong, Y., Wan, Y.-W., Pang, K., Chow, L. M. L., and Liu, Z. (2013). Digital sorting of complex tissues for cell type-specific gene expression profiles. *BMC Bioinformatics*, **14**, 89.
